# Can we fish on stocks that need rebuilding? Illustrating the trade-offs between stock conservation and fisheries considerations

**DOI:** 10.1101/2021.02.25.432880

**Authors:** Vanessa Trijoulet, Casper W. Berg, Claus R. Sparrevohn, Anders Nielsen, Martin A. Pastoors, Henrik Mosegaard

## Abstract

In the Northeast Atlantic, advice for many fish stocks follows the ICES MSY approach, where a zero catch will be recommended if the stock is below its limit reference point, *B_lim_*, and cannot rebuild in the short-term. However, zero catch advice are rarely implemented by managers. This study used medium-term stochastic forecasts with harvest control rules (HCRs) to investigate the consequences of allowing reduced fishing below *B_lim_*. We applied the method to western Baltic herring and North Sea cod, two contrasting species currently estimated below *B_lim_*. We show that the minimum rebuilding probability of 95% required by the MSY approach could be impossible to reach in the short-to medium-term. When this is the case, a lower probability may need to be considered instead in the short-term. Recruitment is the largest source of uncertainty in stock response to management, and can exceed differences between HCRs. Reference points should be estimated in accordance with current recruitment levels if they are to be used for short-term advice or as realistic rebuilding targets. For both stocks, it is possible to keep fishing at reduced levels for similar cumulative catch, SSB and risk on the stock in the medium-term compared to no catch below *B_lim_*. Medium-term trade-offs between stock conservation and fisheries considerations may be needed when fishery closure cannot be implemented in practice.

## 1. Introduction

Globally, one third of the marine fish stocks in the world are fished unsustainably (FAO, 2018), i.e. fish abundance is below the abundance at maximum sustainable yield (MSY). While most of those stocks are not scientifically assessed or managed (Hilborn and Ovando, 2014), several managed stocks are still below their limit reference point in the Northeast Atlantic (ICES, 2019h,d,e,j,k,c) and the Baltic Sea (ICES, 2019b). Usually, fisheries management involves two different processes. First, a scientific process where the stocks are assessed and scientific advice are produced to inform managers on sustainable fishing levels. Second, management decisions are taken at the European or national level and, if relevant, total allowable catches (TACs) are given to fishers

There is currently an international endorsement of the precautionary approach to fishing (Garcia, 1994; Richards and Maguire, 1998), where fish stocks should be sustainably managed while accounting for uncertainty in stock perception, in management and in risk. Fish stocks are usually managed using reference points, which are used as target to get to or stay above (e.g. fishing mortality and biomass at the MSY equilibrium, *F_MSY_* and *B_MSY_* respectively) or trigger (threshold) or limit not to fall below (e.g. limit reference point, *B_lim_*). Harvest control rules (HCRs) are one management tool that is often used to produce advice for managers on fishing mortality (Kvamsdal et al., 2016) and their shapes are often defined using biological reference points. HCR shapes differ between countries but usually fishing mortality is constant and set to its target when stock biomass is above the trigger or threshold reference point. Fishing mortality then decreased linearly with stock biomass between the trigger and the limit reference point. Some HCRs decrease linearly to zero (New Zealand Ministry of Fisheries, 2011; ICES, 2019a) and others decrease to a lower level of fishing mortality (NRC, 2014; ICCAT, 2019).

According to the United Nations Fish Stocks Agreement (UN, 1995), the risk of the stock falling below its limit reference point should be very low. The precautionary approach below the limit reference point has been applied differently around the world and different levels of risk have also be taken. Risk on the stock, estimated as the probability to fall below the limit reference point, vary between jurisdictions and stocks from 5 to 50% (NRC, 2014; ICES, 2019a; ICCAT, 2019). According to Wetzel et al. (2016), rebuilding probability should be above 60% to allow successful rebuilding above the target within the necessary time frame. Some jurisdictions acknowledge that fishing mortality should be reduced as much as possible when stock biomass is below the limit reference point (NAFO, 2004; DFO, 2009; Australian Government, 2018; ICCAT, 2019) and others allow for fishery closure (New Zealand Ministry of Fisheries, 2011; ICES, 2017a). While precautionary considerations are necessary at low stock size since stock dynamics at these levels are usually unknown and poorly understood; reduced or no fishing below the limit reference point conflicts with fisheries socio-economic considerations.

In the Northeast Atlantic, the International Council for the Exploration of the Sea (ICES) provides advice on sustainable fishing opportunities for many fish stocks upon request from clients. Clients are mostly fisheries authorities at the international or national level, which use the advice to manage their stocks. Managers request single species advice that ensures sustainable fisheries management but while all clients are subject to the same international agreements, policies may differ at national levels. Many of the stocks are shared between different fisheries authorities so ICES develops advice consistent with common international agreements. Advice in ICES is given annually by performing short-term stock projections from the intermediate year to the advice year (intermediate year +1) under different fishing scenarios. The outputs of these scenarios are presented in the advice sheet but only one scenario is used as advice for managers (stated in the headline of the advice sheet). The advice follows the modeling and reference point assumptions that were agreed on during the last benchmark^1^ for the stock.

For stocks with reliable quantitative or qualitative assessments (ICES category 1 and 2 stocks), when an agreed and evaluated management plan is not already in place between the different fisheries authorities managing the stocks, ICES advice is based on the maximum sustainable yield (MSY) approach, a framework that is based on MSY considerations and consistent with the precautionary approach (ICES, 2019a). The MSY approach is based on an advice rule that uses reference points as thresholds and targets that define the shape of the HCR. The MSY approach notably stipulates that fishing mortality cannot exceed *F_MSY_*, becoming therefore the maximum *F* that would be advised on a stock. A reduction in *F* would be recommended if the stock is below the trigger reference point, *MSY B_trigger_* ^2^. Also, according to the MSY approach, the stocks should be above the biomass limit reference point, *B_lim_*^3^ with a 95% probability, meaning that the risk for the stock to fall below *B_lim_* should be below 5%. As a result, *F_MSY_* is estimated in ICES so that there is less than 5% chance that applying the HCR described above would lead to the stock falling below *B_lim_* in the long-term. The precautionary constraint in the estimation of *F_MSY_* by ICES differs from other approaches around the world, and necessitates an estimate for *B_lim_* before MSY reference points can be estimated. ICES includes an additional precautionary consideration in its advice rule that if a stock is below *B_lim_* and cannot rebuild above *B_lim_* in a two-year forecast when *F* = 0, zero catch for the following year would be recommended to allow for the stock to rapidly rebuild above *B_lim_* (ICES, 2019a).

In 2019, 18 stocks (category 1 and 2) assessed by ICES were estimated below *B_lim_* and of those, 13 got a zero catch advice for 2020. However, zero catch is rarely implemented by managers (Rosenberg et al., 2006; Cadrin and Pastoors, 2008; Kvamsdal et al., 2016). For instance, among these 13 stocks, at least 7 got a positive TAC in 2020 (EU, 2019b; EU and Norway, 2019, 2020). One possible reason is that, after implementation of the Landing Obligation (EU, 2013), zero catch cannot be realistically achieved in practice, because many stocks are fished by multiple fleets and sometimes in a mixed fishery . Therefore, a zero catch on a stock could mean an impossibility of fishing on other stocks in an entire geographical area and several fleets going out of business. In other jurisdictions such as in the US, zero catch is never advised as not realistically implementable due to socio-economic and mixed fisheries considerations (U.S. Government, 2007). In case of zero catch advice in ICES, there is little guidance for managers to set a TAC that balances biological and socio-economic sustainability considerations in the medium-term. The medium-to long-term consequences on fish stocks of not following zero catch advice are unclear. It seems therefore necessary to understand the trade-offs between keeping a yield to save a fishery and risking a stock collapse or slowing a rebuilding rate.

In this study, we investigated the consequences of allowing fishing when a stock is below *B_lim_* compared to the current MSY approach, by developing a stochastic forecast that tests different HCRs setting annual fishing quotas. We were notably interested in the effects induced by these HCRs, on the stock, in terms of stock size and risk, but also on the fishery, in terms of future cumulative catch. We chose to look at different time frames in the short- (0-5 years) to medium-term (5-10 years). Current scientific advice in ICES is largely based on deterministic short-term forecasts (2 years), we simply extended this period and included uncertainties in the key processes to investigate longer-term consequences. The forecast was built to account for uncertainty in the current perception of the stock but also in future dynamics. We did not look at long-term simulations as we were interested in the rebuilding period between unsustainable and sustainable stock levels (Gröger et al., 2007; Rosenberg and Brault, 1991).

The simulations were based on different recruitment assumptions, fishing scenarios and fishing targets. The different recruitment assumptions enable to investigate stock rebuilding for different plausible recruitment regimes. We focused on differentt fishing scenarios that include HCRs where fishing mortality varies with SSB according to limit and target reference points, where reduced fishing is possible or not below *B_lim_*. The forecast was developed as an extension to the `stockassessment R` package for the state-space assessment model (SAM) (Nielsen and Berg, 2014; Berg and Nielsen, 2016).

We applied the stochastic forecast to two fish stocks currently assessed with SAM: North Sea cod and western Baltic spring-spawning (WBSS) herring. These stocks were chosen because they are currently considered at risk according to the ICES precautionary approach but they contrast by the fact that WBSS herring is a pelagic species that has been recommended a zero catch for two years (ICES, 2018a, 2019i) while North Sea cod is a demersal fish stock that, despite being currently estimated below *B_lim_*, still is estimated to recover in the short-term in the 2019 advice (ICES, 2019f), and was therefore not subject to recommended fishing closure for 2020. WBSS herring is a good example of current failure of advice being implemented into management as the zero catch advice was never applied in practice. The European Union and Norway still allowed positive TACs to be taken in 2019 and 2020 in the western Baltic, Kattegat and Skagerrak (EU, 2018b, 2019a,b; EU and Norway, 2019, 2020), despite a zero catch advice for 2019 and 2020 (ICES, 2018a, 2019i). The response from managers was to reduce the TACs by 42% from 2018 to 2019 (EU, 2017, 2018a) and by 13% from 2019 to 2020, resulting in a current TAC in ICES Subdivisions 20-24 for 2020 of 34337 tonnes. WBSS herring is taken as bycatch in the industrial sprat fishery in ICES Division 3.a as well as in mixed stock catches in ICES Divisions 3.a and 4.a East (ICES, 2019l). A zero catch would mean closing the Norwegian herring fishery in the eastern North Sea and industrial fisheries operating in 3.a, and this can explain why it was never applied by managers. Cod is caught in mixed fisheries and is the choke species^4^ for many fleets in the North Sea (ICES, 2019g). Such as for WBSS herring, if in the future cod is not estimated to rebuild in the short-term forecasts used for scientific advice, closing the fishery for cod would have important consequences for the fishing industry and has therefore little probability of ever being applied in practice.

## 2. Methods

### 2.1. Stochastic forecasts

We developed the forecast function in the SAM package to enable the outcome of HCRs to be implemented in the assessment projections for both the single and multifleet versions of the package. The source code can be found at https://github.com/vtrijoulet/SAM/tree/master2 for the single fleet version and at https://github.com/vtrijoulet/SAM/tree/multi2 for the multifleet version.

#### 2.1.1. Forecast method

The stochastic forecast can be run for a specified number of years as a continuity of the assessment model. The process error assumptions from the assessment model are used in the forecast, e.g. process errors in fish numbers, except for process error on fishing mortality. The last year of the assessment model is used as starting point for the forecast. The estimates in the last year of the assessment and their uncertainty are used to simulate replicates of stock numbers at age and fishing mortality at age for each fleet This enables to account for the uncertainty in the perception of the current stock level. In this study, 5000 replicates were used after checking they were enough to get outputs that were not sensitive to the number of iterations.

Similarly to what is used in forecasts used for advice, in the intermediate year, catch is fixed to the agreed TAC for this fish stock. A management strategy is applied thereafter and until the end of the forecast period to all replicates independently.

For convenience, in this study, simulation trials (STs) are defined by a combination of recruitment assumption, fishing scenario or harvest control rule (HCR) and target and limit reference points that inform the shape of the HCR. The different possible ST options are described below.

#### 2.1.2. Recruitment assumptions

The forecast simulates population development and fishery under three contrasting recruitment assumptions to investigate the robustness of HCR performance to different productivity regimes:

*•* Recruitment is randomly sampled with replacement from the assessment model recruitment estimates in specified assessment years: this is the most commonly used assumption in short-term forecasts used for scientific advice. For instance, if recruitment is sampled in the final assessment years, this is equivalent to assuming recruitment stays at current levels.
*•* Hockey-stick stock recruitment relationship: recruitment varies with spawning stock biomass (SSB) following a segmented regression (hockey-stick) relationship (Barrowman and Myers, 2000). The parameters of the stock-recruitment relationship are estimated using the assessment model estimated pairs of SSB and recruitment in specified years. The standard deviation of the recruitment residuals was used to inform random errors around the recruitment estimates resulting from the hockey-stick relationship. This recruitment assumption is what resemble the most the assumption taken when estimating the MSY reference points for ICES fish stocks.
*•* Random walk with negative drift: log recruitment follows a random walk that decreases over time given a specific proportion of the variance estimated for recruitment in the assessment model (if random walk was assumed in the assessment model). Median recruitment decreases over time from the last year of assessment but the variance of log recruitment increases linearly with time whereas the first two recruitment assumptions have constant variance. It is important to note that while a random walk model is adequate for short-term forecasts, recruitment variations under this assumption could increase with time to unrealistic high values so it may not be realistic for longer forecasts.

#### 2.1.3. Harvest control rules

Management was assumed to control the average total fishing mortality (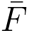) on a fish stock in a forecast year as a function of the SSB in the previous year via HCRs. The shape of the HCRs is informed by a reduced fishing mortality level and three reference points: (i) *MSY B_trigger_*, which is the biomass reference point that triggers a cautious response within the ICES MSY framework; (ii) *B_lim_*, the limit reference point and (iii) *F_target_* which is the maximum fishing mortality that is applied on the stock. In the ICES MSY approach, it is set to *F_MSY_* .

The HCRs were defined following discussion with a stakeholder representative and differed in how 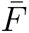 varied when SSB was below *MSY B_trigger_*as illustrated in Figure 1. HCR 1 corresponds to the ICES MSY approach

**Figure 1:**
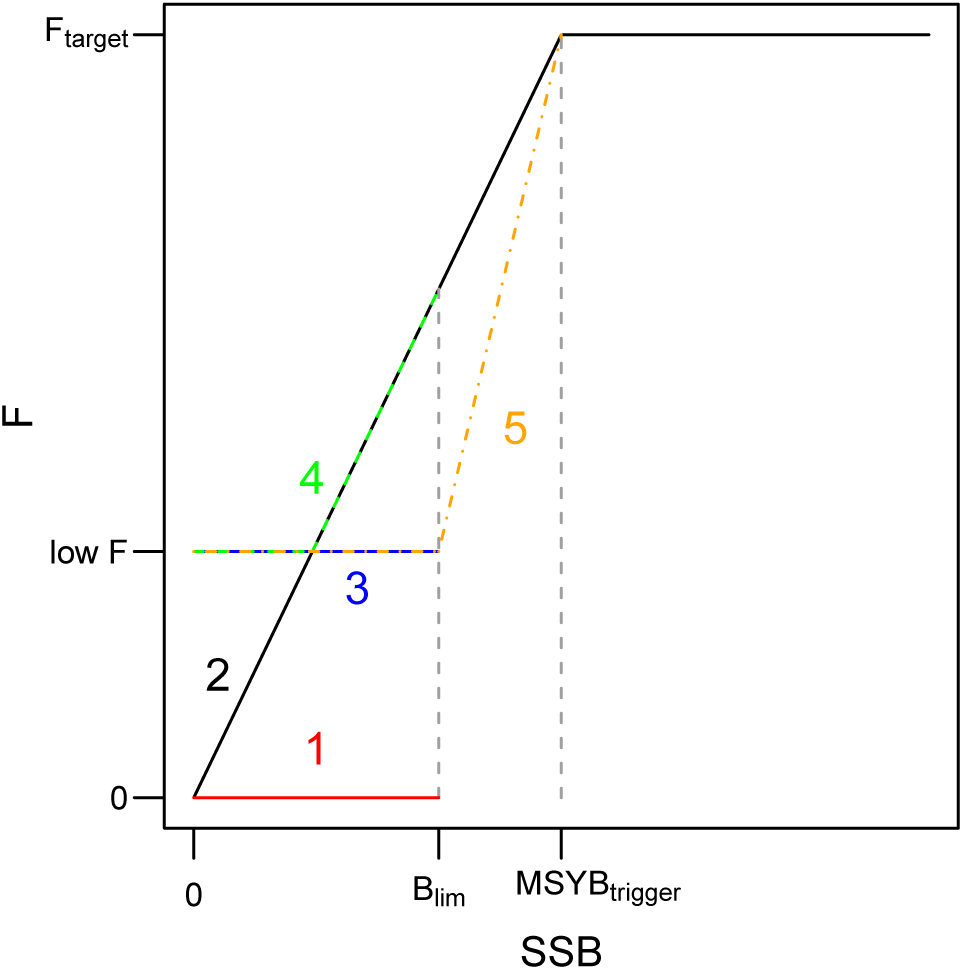
Harvest control rules (HCRs) developed in the stochastic forecast. Five HCRs can be chosen from (1-5) for any given set of reference points (*F_target_*, *B_lim_* and *MSY B_trigger_*) and *lowF* level.

(when the stock cannot rebuild in the short-term), where there is no fishing when SSB is below *B_lim_*. For HCR 2, 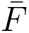 decreases linearly from the targeted fishing mortality (*F_target_*) at *MSY B_trigger_* to 0 when *SSB* = 0. HCR 3 is an option where 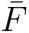 is reduced to a specified low value when SSB is below *B_lim_*. HCR 4 is similar to HCR 2 but 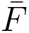 is fixed to this low level for low values of SSB. Finally, HCR 5 is the case where 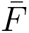 is reduced to a constant low level when SSB is below *B_lim_* and increases linearly to *F_target_* when SSB increases from *B_lim_* to *MSY B_trigger_* .

### 2.2. Application to WBSS herring and North Sea cod

For WBSS herring, the stochastic forecast used the results from the 2019 assessment (ICES, 2019l). Assessment model scripts, data files and results are publicly available on stockassessment.org under the name “WBSS HAWG 2019”. The assessment was run for the years 1991 to 2018. For North Sea cod, the 2019 assessment results were used (ICES, 2019m) and can be found on `stockassessment.org` under the name “nscod19 ass01 October”. Cod was assessed from 1963 to 2019.

#### 2.2.1. Simulation trials parameterization

The assumptions used for the recruitment assumptions for herring and cod are summarized in Table 1. The years used for the recruitment randomly sampled from the assessment estimates were the same as those used in the latest scientific advice for each species (ICES, 2019i,f). Recruitment being low at the end of the time series for both species, this assumption resulted in assuming recruitment in the forecast will stay at current low levels. For the hockey-stick recruitment assumption, the starting year used for the SSB and recruitment pairs was the same as the one used to estimate reference points for both stocks (ICES, 2018c, 2017b, 2015). This recruitment assumption is the one that resembles the reference point estimation assumptions the most. However, it is not identical as reference points for both stocks were estimated using earlier estimates of SSB and recruitment, available at the time of the benchmarks. Also, note that the reference point estimates were obtained using the “Eqsim” procedure based on a weighted average of Ricker, Beverton-Holt and hockey stick stock-recruitment curves. This recruitment assumption is the most optimistic as it assumes that recruitment may increase in the future if SSB increases but it is also the only assumption that can simulate stock collapse below *B_lim_*, as the curve decreases to zero. In the most recent assessment period, recruitment has been low but with a slight decreasing trend for herring and stable since 1998 for cod but slightly decreasing since 2010. Therefore, to cover a more pessimistic development for both stocks, a further decrease in median recruitment was implemented. Both stocks were assessed in 2019 assuming a random walk in recruitment in SAM. We used the estimated standard variation of the estimated recruitment for each stock to model a random walk with a negative drift of 10% these standard deviations. This proportion was chosen to get a smooth decrease in median recruitment over time for both species without producing a too unrealistic steep decline. The random walk assumption was the most pessimistic assumption since it assumed median recruitment decreased over time. The resulting recruitment for the three assumptions are illustrated in Appendix A1. Comparing the three recruitment option outputs should give a large range of plausible recruitment scenarios from pessimistic (random walk with drift), neutral (randomly sampled) to more optimistic (hockey-stick).

**Table 1:** Summary of recruitment assumptions for both stocks. The hockey-stick parameters *SSB_i_* and *R_i_* are the x and y coordinates respectively of the inflection point of the stock-recruitment curve with their coefficient of variation (CV) given in parenthesis and *sd* is the standard deviation around the recruitment residuals on log scale. *log*(*σ*) is the estimate of the log of the standard deviation for the random walk taken from the SAM fit (aka. *logSdLogN*) and used in the forecast. The *drift* parameter is the percentage of standard deviation used to decrease the mean of the random walk in the forecasts.

| Assumptions | Sampled |  | Hockey-stick |  | Random walk |  |
| --- | --- | --- | --- | --- | --- | --- |
| Stocks | Herring | Cod | Herring | Cod | Herring | Cod |
| Years | 2013-2017 | 1998-2018 | 1991-2018 | 1988-2019 |  |  |
| Parameters | | | $SSB_i = 218788$ (CV=0.38)<br>$R_i = 3923183$ (CV=0.35)<br>$sd = 0.35$ | $SSB_i = 81331$ (CV=0.05)<br>$R_i = 283433$ (CV=0.13)<br>$sd = 0.64$ | $\log(\sigma) = -1.33$<br>$drift = 10\%$ | $\log(\sigma) = -0.23$<br>$drift = 10\%$ |
| Figure | A1.1 | A1.2 | A1.3 | A1.3 | A1.1 | A1.2 |

Reference points and reduced fishing mortality value used to parameterize the HCRs are summarized in Table 2. Three *F_target_* levels were considered for the forecasts. *F_MSY_* is commonly used in management advice and is notably the current *F_target_* in the MSY approach used for management of both WBSS herring and North Sea cod. It was therefore chosen as main *F_target_* in the study. We also investigated the effect of reducing or increasing the *F_target_* to reflect its uncertainty or even a change in future *F_MSY_* . We used two other *F_target_* levels for each species defined as the EU Baltic Sea Multiannual Plan 2018 lower and upper reference points (*F_lower_* and *F_upper_* ^5^) for herring (European Commission, 2018a; ICES, 2019i) and the North Sea Multiannual Plan 2018 for cod (European Commission, 2018b; ICES, 2019f). The low fishing mortality (*lowF*) for HCRs 3-5 is set to 0.1 for both species, which corresponds to around one third of *F_MSY_* .

**Table 2:** Reference points and *low*F values used to parameterize the HCRs in this study.

| Stock | Herring | Cod | Reference |
| --- | --- | --- | --- |
| $MSYB_{trigger}$ (t) | 150000 | 150000 | ICES (2018c, 2019i, 2017b, 2019f) |
| $B_{lim}$ (t) | 120000 | 107000 | |
| $F_{target}$ ($y^{-1}$) | $F_{MSY} = 0.31$ | $F_{MSY} = 0.31$ | European Commission (2018a); ICES (2019i);<br>European Commission (2018b); ICES (2019f) |
| | $F_{lower} = 0.216$ | $F_{lower} = 0.198$ | |
| | $F_{upper} = 0.379$ | $F_{upper} = 0.46$ | |
| $lowF$ ($y^{-1}$) | 0.1 | 0.1 | |

The catch constraint used in the stochastic forecast for the intermediate year (2019) was the same as the one used in scientific advice for both species (ICES, 2019i,f) and is summarized in Table A2.1. The forecast was run for 10 years so to 2028 for both species. In this study, a total of 51 STs were considered. Two base cases are considered for each of the 3 recruitment options (so 6 STs): one where no fishing occurs for all forecast years after the intermediate year (option “*F* = 0”), and another where the catch by fleet in 2019 is kept constant for all remaining years (option “constant 2019 catch”). The other 45 STs were comprised of the 3 recruitment options, the 3 different values of target fishing mortality (*F_target_*) and the 5 HCRs.

#### 2.2.2. Performance metrics

Resulting forecast trajectories for SSB, average fishing mortality (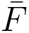_3_*_−_*_6_ for herring and 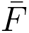_2_*_−_*_4_ for cod), total catch and cumulative catch were compared for the 51 STs by taking the median and 95% confidence interval of the 5000 replicates. We compared time for median stock recovery above *B_lim_* for the all options. The risk on the stock for every forecast year with management (2020-2028) is calculated as the probability of SSB falling below *B_lim_* using the proportion of replicates where *SSB < B_lim_*.

#### 2.2.3. Additional long-term forecast analysis

To investigate the impact of fishing at the different equilibrium *F_target_* on equilibrium stock size and rebuilding probability, we ran 6 additional stochastic long-term forecasts for both species where 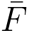 is fixed at the different *F_target_* values for the two recruitment assumptions where recruitment is randomly sampled in the assessment estimates and when recruitment follows the hockey-stick stock-recruitment relationship. The forecasts were run until the steady-state SSB was reached. We did not run long-term forecast for the random walk with negative drift recruitment assumption because this assumption is not adequate for long-term forecasts. We looked at equilibrium SSB and probability of falling below *B_lim_* for these results.

In addition, while the hockey-stick assumption resembled the assumption taken when estimating reference points for both stocks, it was not identical (cf. section 2.2.1). As a result, the equilibrium fishing and biomass corresponding to the recruitment assumptions taken in this study may differ from the reference points used in the HCRs. As a result, we calculated MSY reference points for the same two recruitment assumptions. Note that the MSY reference points estimated in this study should be only considered for illustration. More information on how these were estimated is given in Appendix A3. We did not consider the extra precautionary constraint implied by ICES (less than 5% probability of falling below *B_lim_*) in the estimation of *F_MSY_* . When recruitment is randomly sampled from the assessment estimates, *F_MSY_* corresponds in fact to *F* at maximum yield-per-recruit, *F_max_*.

## 3. Results

An example of forecast trajectories for SSB, 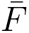 and total catch is given Figure 2 for HCR 1 (MSY approach with no fishing below *B_lim_*) with recruitment sampled from the assessment estimates, so staying at current low levels. Here, cod rebuilt rapidly above *B_lim_* after closure of the fishery for two years no matter the *F_target_* value, whereas herring necessitated three years of closure and presented oscillations around *B_lim_* when *F_target_ ≥ F_MSY_* . This resulted in consequent large changes in fishing mortality and total catch in the forecasts with 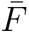 oscillating between 0 and large values over time.

**Figure 2:**
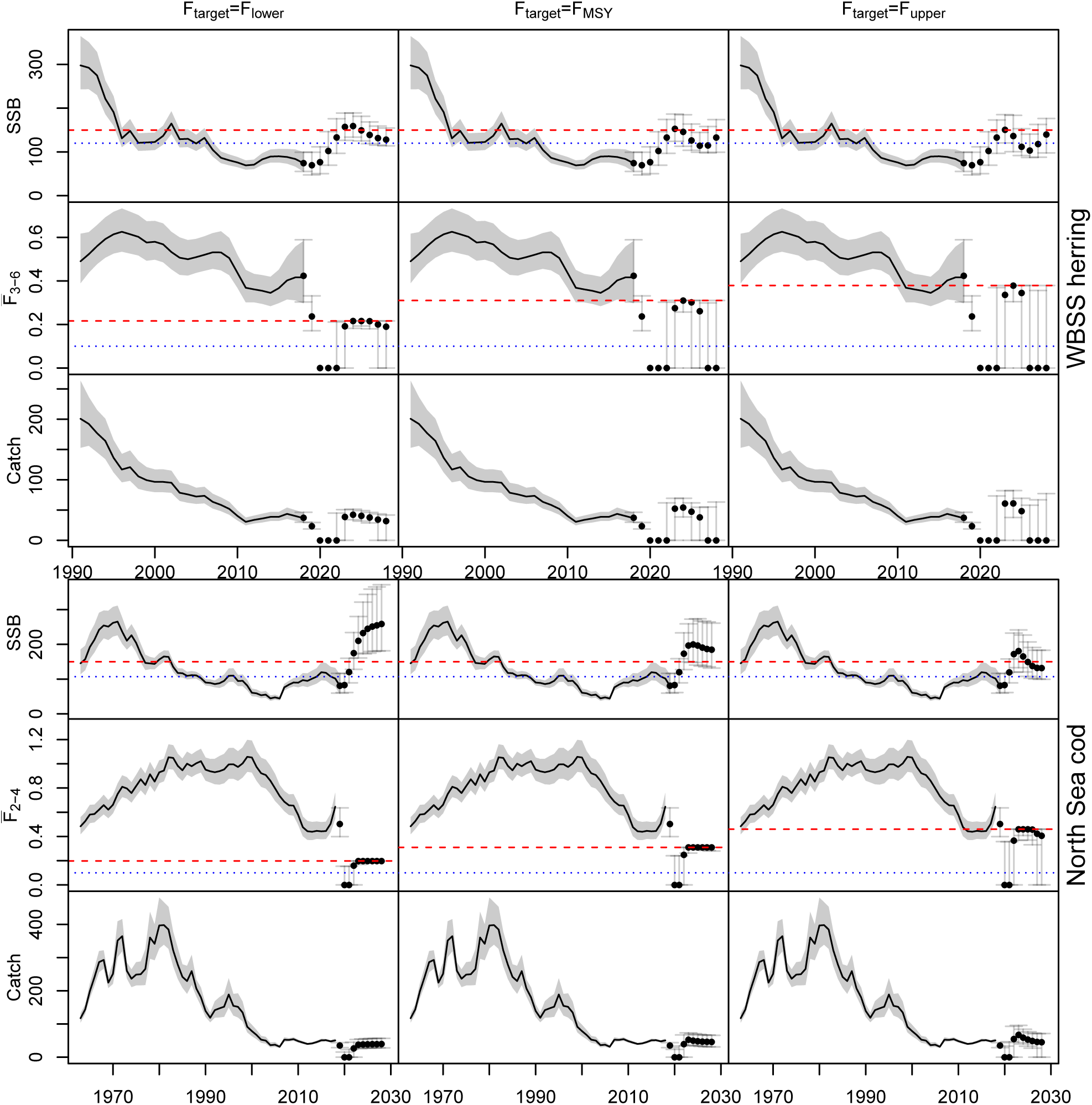
Historical and forecast spawning stock biomass (SSB), mean fishing mortality and total catch for HCR 1 (MSY approach) with recruitment sampled from the assessment estimates. The points are medians and the segments the 95% confidence intervals estimated from the 5000 replicates. For SSB, the red dashed line is *MSY B_trigger_* and the blue dotted line is *B_lim_*. For 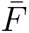, the red dashed line is the *F_target_* and the blue dotted line is the reduced *F* = 0.1.

The results for all STs (all recruitment assumptions, *F_target_*, HCRs and base cases) are given in A4. The results were very different between fish stocks and were sensitive to the recruitment assumption. WBSS herring showed difficulties to get a stable recovery above *B_lim_* over time for all HCRs and recruitment assumptions (Figures A4.3 and A4.19) except when recruitment was more optimistic and increased when SSB increased (hockey-stick assumption, Figure A4.11). For all recruitment scenarios, some big jumps in fishing mortality were observed notably for HCR 1, HCR 3, and HCR 5 where 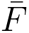 varied between its lower limit (0 for HCR 1 or 0.1 for HCRs 3 and 5) and higher values while the other HCRs had a smoother change in 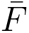 (Figures A4.4, A4.12 and A4.20). These changes induced subsequent variations in total catch (Figures A4.5, A4.13 and A4.21). North Sea cod rebuilt in few years above *B_lim_* and even sometimes *MSY B_trigger_* no matter the ST considered (Figures A4.2, A4.7, A4.15 and A4.23). Even if median rebuilding was successful in medium-term, for all recruitment and *F_target_* assumptions, it still necessitated reducing 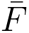 for 2 years for all HCRs notably to its lower limits 0 and 0.1 for HCR 1 and HCRs 3, 5 respectively (Figures A4.8, A4.16 and A4.24).

Figure 3 summarizes the time frame for median recovery for all STs resulting in rebuilding for WBSS herring and North Sea cod. For WBSS herring, about 73% (37 out of 51) of the STs allowed rebuilding above *B_lim_* but the recovery was often not stable to the end of the simulation with the median SSB declining below *B_lim_* after few years. Stable herring recovery by the end of the simulation (SSB went above and stayed above *B_lim_*) was reached for 51% (26 out of 51) of the STs. Having a low value for *F_target_* increased the probability for medium-term (2028) recovery. Only HCR 1 enabled stock rebuilding as quickly as the fishing closure base case *F* = 0 and most of the other options allowed rebuilding a year or two later. All STs applied to North Sea cod resulted in stable rebuilding of median SSB above *B_lim_* by the end of the forecast period. For 82% (42 out of 51) of the STs rebuilding above *B_lim_* started in 2021, time frame considered for current scientific advice. Keeping the 2019 catch constant was the ST showing the slowest rebuilding and rebuilding time was sensitive to the stock recruitment assumption.

**Figure 3:**
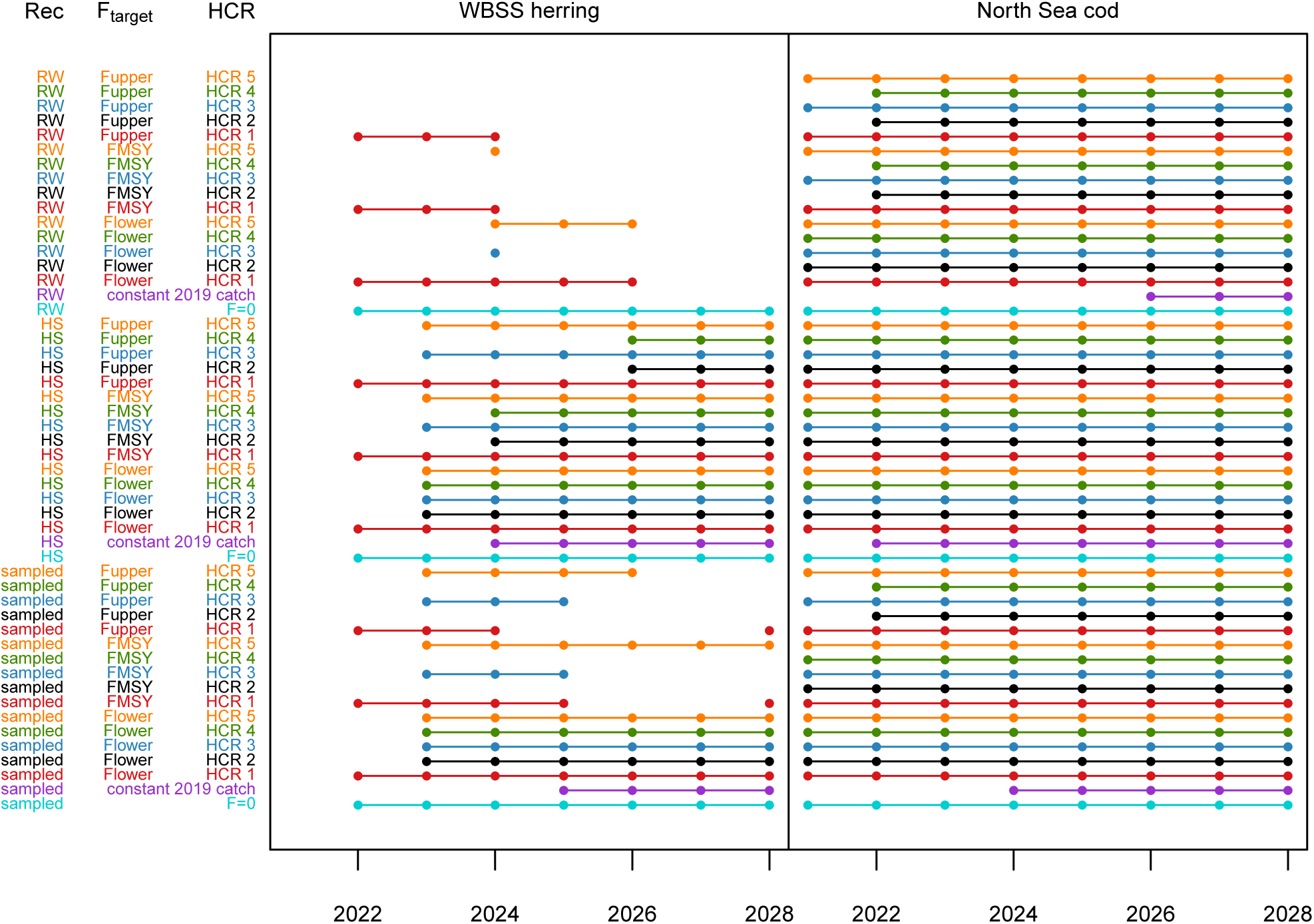
Simulation trials (STs) for which median SSB recovered above *B_lim_*. The points represent the years for which *SSB ≥ B_lim_* in the forecasts. The lines highlight the consecutive years where *SSB ≥ B_lim_*. Each row is a ST (set of recruitment (Rec), *F_target_* and HCR assumption) and 6 STs are base cases where *F* = 0 or the catch in 2019 is kept constant. The notation “sampled” is given when recruitment is randomly sampled from the assessment estimates, “HS” when log recruitment follows an hockey-stick stock-recruitment relationship, and “RW” stands for random walk with negative drift.

Figure 4 shows the median total catch against the probability of SSB falling below *B_lim_* for HCR 1. It highlights the oscillating behavior (cf. Figure 2) observed for WBSS herring when median recruitment was assumed to stay at the current low levels (sampled in 2013-2017) or decreased further (random walk with negative drift). Similar behavior was observed for HCRs 3 and 5 (Figures A4.6 and A4.22). For all recruitment assumptions and *F_target_* levels, the risk on herring decreased in the first forecast years (2020-2022) following the absence of fishing. Then median catch increased and risk stayed low as fishing started again. However when *F_target_ ≥ F_MSY_*, for all recruitment assumptions except the hockey-stick, median catch decreased and risk increased until the fishery was closed again until the end of the forecast period. When recruitment increased with increasing SSB, risk on herring stayed below 20% by 2028. Similar figures for all STs and for North Sea cod are given in Appendix A4. Similarly oscillations can be seen for cod, to a lesser degree, for the random walk assumption (Figure A4.26).

**Figure 4:**
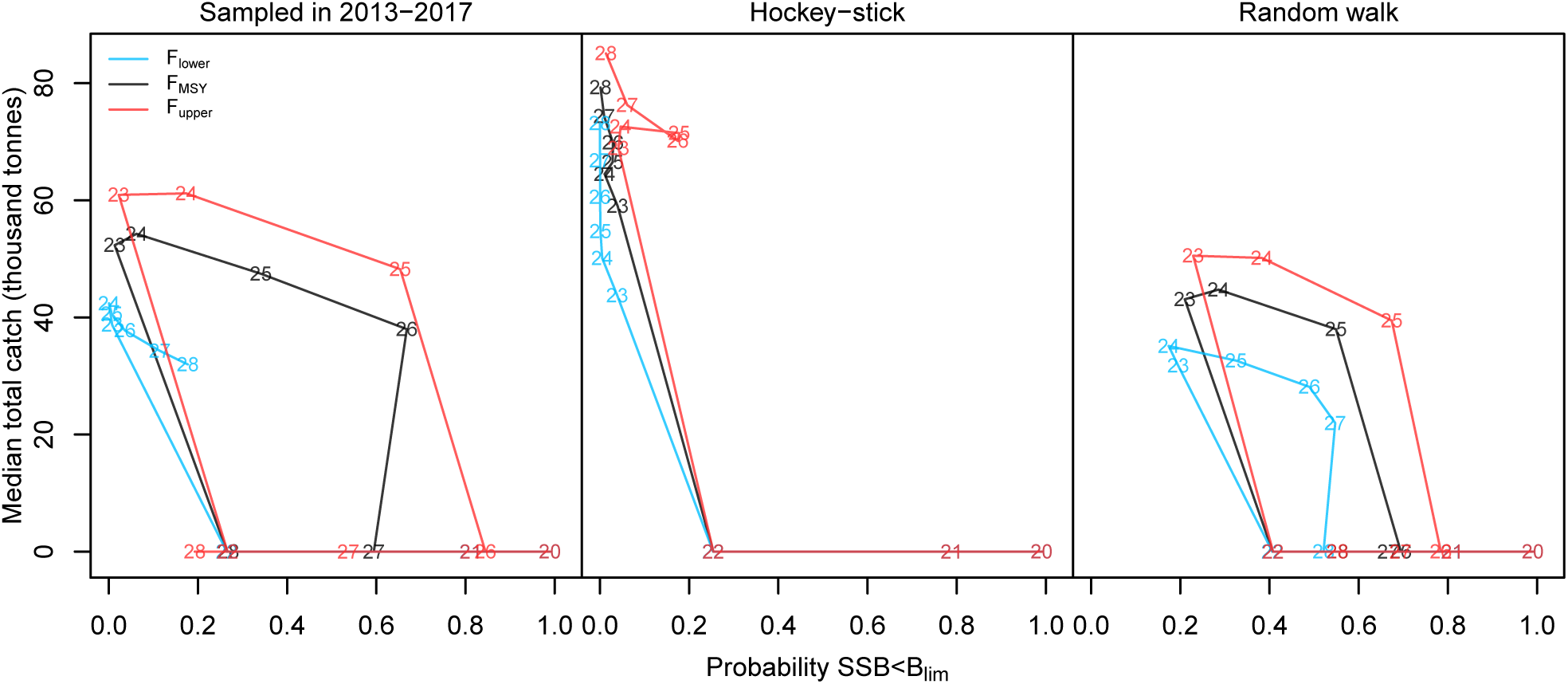
WBSS herring median total catch against probability of SSB falling below *B_lim_* for HCR 1 (MSY approach). The numbers correspond to the forecast years (2020-2028). Each color represents a *F_target_* option.

Figure 5 summarizes the change in WBSS herring median cumulative catch against median SSB in the forecast years for all STs except *F* = 0. It is clear that median recovery between *B_lim_* and *MSY B_trigger_* (yellow zone) or above *MSY B_trigger_* (green zone) is difficult for herring when median recruitment stayed at current low levels (sampled in 2013-2017) or decreased (random walk with negative drift). Overall, all the HCRs resulted in similar level of cumulative catch by the end of the forecast period but the path to get there and the final median SSB differed between HCRs.

**Figure 5:**
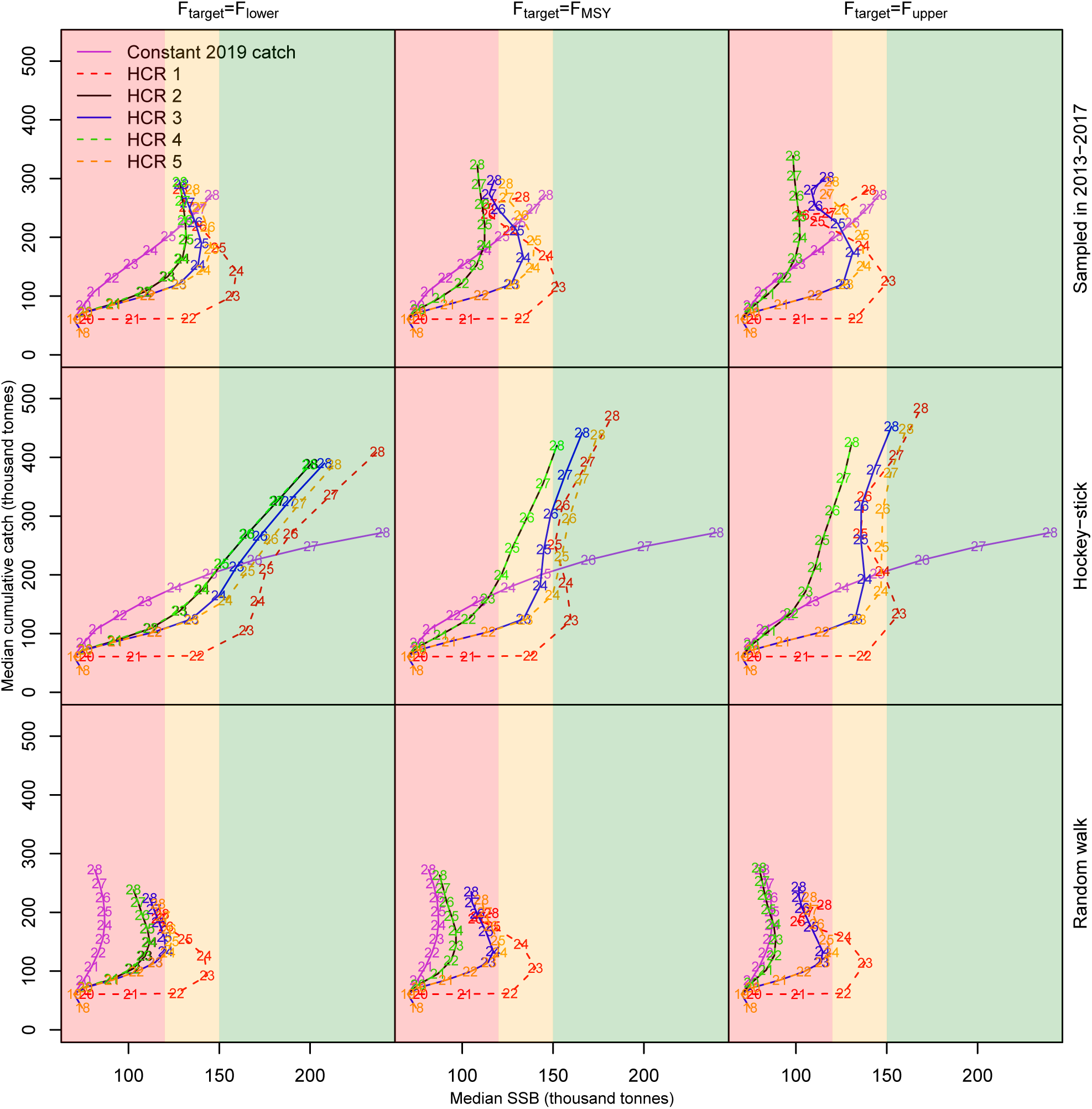
WBSS herring median cumulative catch against median SSB in the forecast period. The numbers correspond to the forecast years (2019-2028). The red zone is when *SSB < B_lim_*, the yellow zone is when *B_lim_ ≤ SSB ≤ MSY B_trigger_*, the green zone is when *SSB > MSY B_trigger_* .

Similar results including variation in replicates are presented in Figure 6 for *F_target_* = *F_MSY_*, current *F_target_* used in ICES, and for three different time frames (short-term 2022, early medium-term 2025 and late medium-term 2028). Overall, for herring, it was difficult to get to a probability of SSB falling below *B_lim_* of less than 5%, except in 2028 when recruitment was assumed to increase when SSB increases (hockey-stick). In the medium-term (2025 and 2028), HCR 1 and HCR 3 resulted in similar cumulative catch, SSB and risk on the stock but HCR 3 allowed for fishing at low level below *B_lim_* while HCR 1 assumed fishery closure below *B_lim_*. HCR 1 was overall more variable than the other STs, due to larger differences in 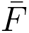 between replicates (cf. Figures A4.4, A4.12 and A4.20) .

**Figure 6:**
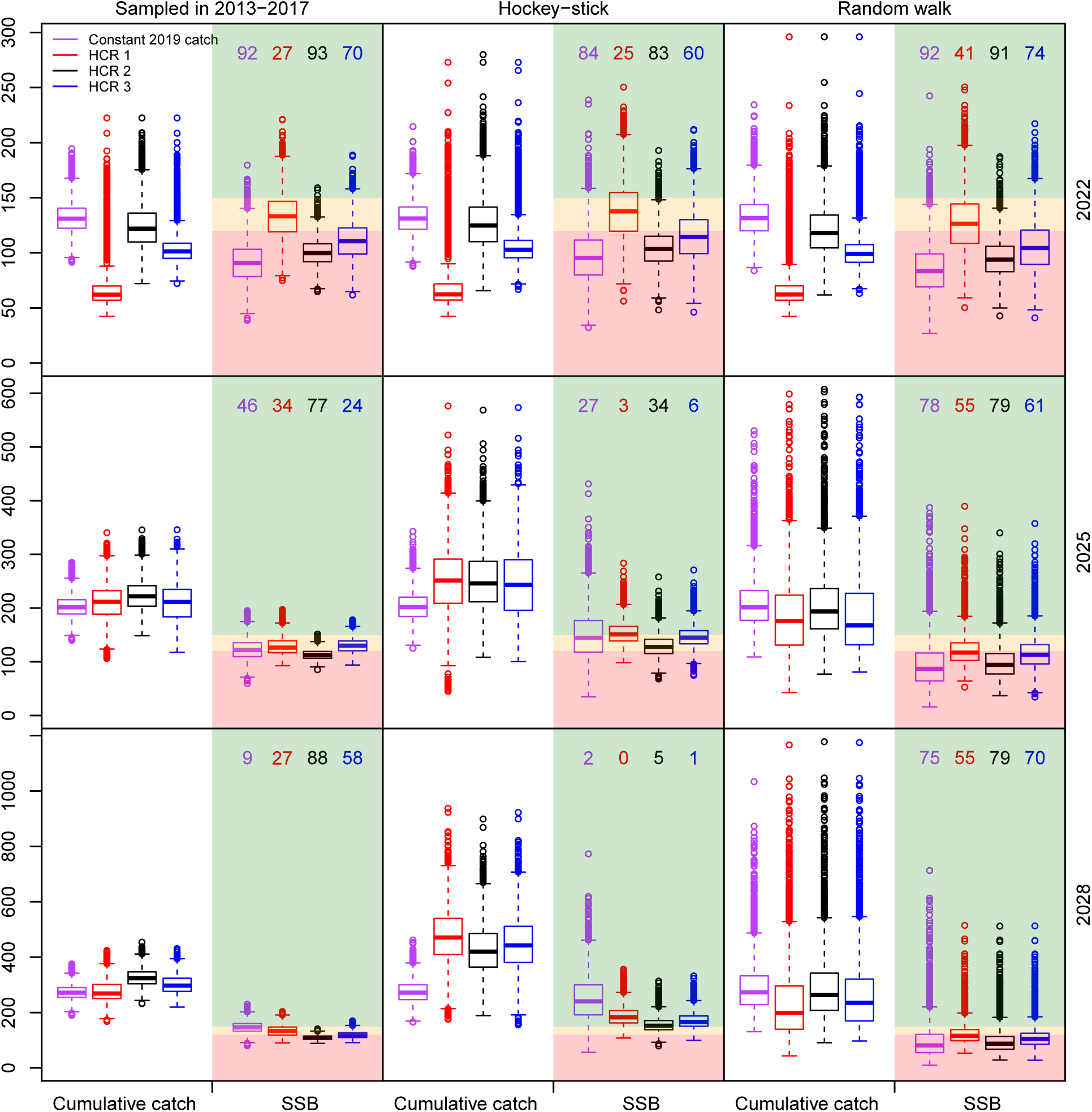
Boxplot of WBSS herring cumulative catch and SSB (thousand tonnes) from the 5000 replicates when *F_target_* = *F_MSY_* for different time frames (2022, 2025 and 2028) and for 4 chosen STs. The numbers are the probability of SSB falling below *B_lim_* in percent for the corresponding ST. The red zone is when *SSB < B_lim_*, the yellow zone is when *B_lim_ ≤ SSB ≤ MSY B_trigger_*, the green zon^2^e^5^is when *SSB > MSY B_trigger_* .

Comparing median cumulative catch with median SSB over time for cod (Figure 7) highlights that no matter the HCR considered and the assumption on recruitment and *F_target_*, the level of cumulative catch and SSB obtained at the end of the forecast period were similar despite different fishing behaviors assumed below *MSY B_trigger_* (cf. Figure 1). Assuming the 2019 catch constant in the forecast resulted in largely different results compared to the HCRs. Getting to a risk on the cod stock lower than 5% was possible to achieve in the short-term (2022) for HCRs 1-3 and for all recruitment assumptions except the random walk despite the fact that this latter assumption resulted in positively skewed SSB (Figure 8). Similar to what was observed for WBSS herring, HCR 1 showed larger span in cumulative catch and SSB than other STs, notably when recruitment was more variable (hockey-stick and random walk). The variations in the cumulative catch for all forecast years are given for both species in Figures A4.27 and A4.28.

**Figure 7:**
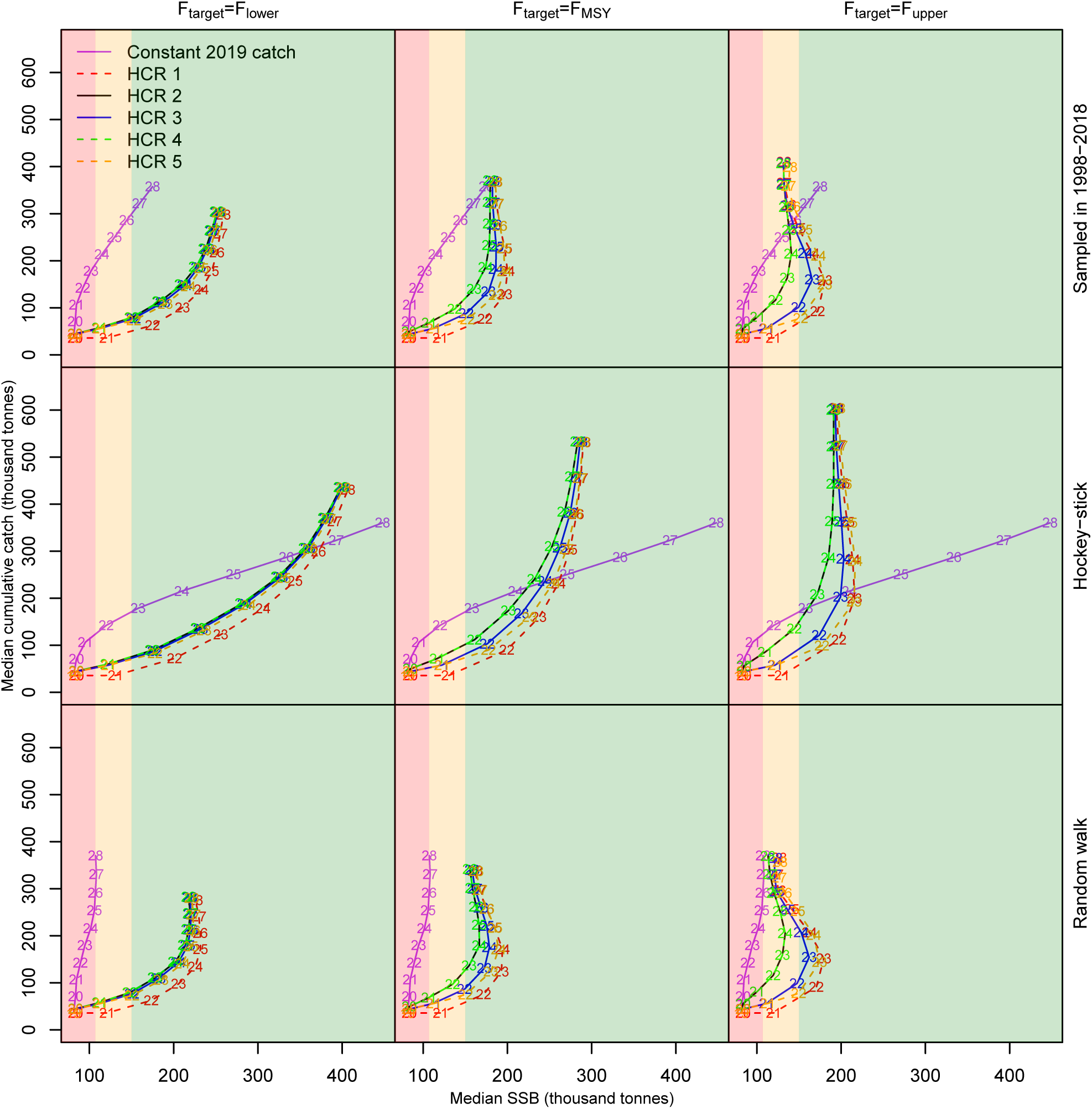
North Sea cod median cumulative catch against median SSB in the forecast period. The numbers correspond to the forecast years (2019-2028). The red zone is when *SSB < B_lim_*, the yellow zone is when *B_lim_ ≤ SSB ≤ MSY B_trigger_*, the green zone is when *SSB > MSY B_trigger_* .

**Figure 8:**
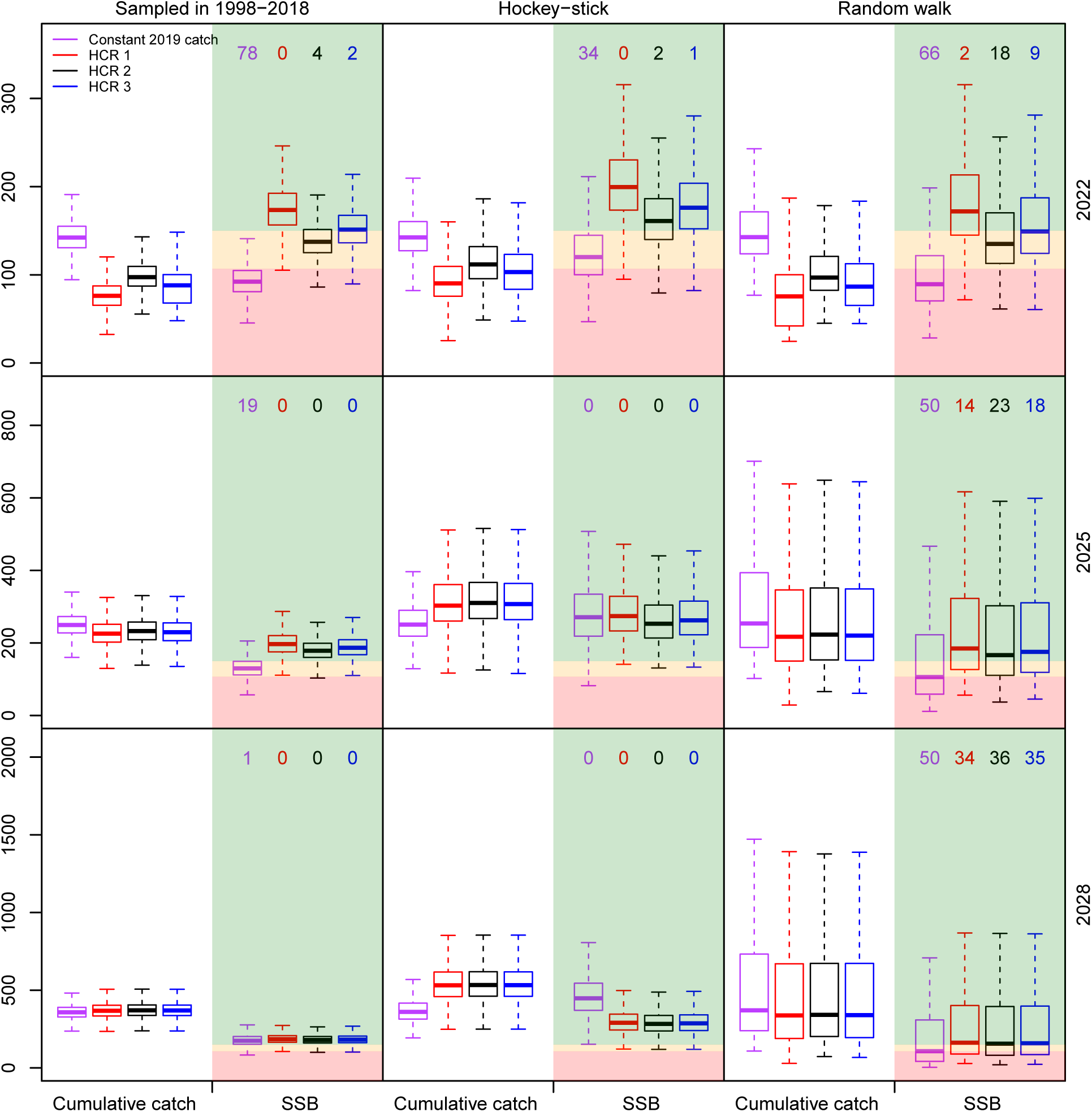
Boxplot of North Sea cod cumulative catch and SSB (thousand tonnes) from the 5000 replicates when *F_target_* = *F_MSY_* for different time frames (2022, 2025 and 2028) and for 4 chosen STs (constant 2019 catch and HCRs 1-3). The numbers are the probability of SSB falling below *B_lim_* in percent for the corresponding ST. The red zone is when *SSB < B_lim_*, the yellow zone is when *B_lim_ ≤ SSB ≤ MSY B_trigger_*, the green zone is when *SSB > MSY B_trigger_* . Please note that due to very large confidence intervals for the random walk recruitment assumption (cf. Figure A4.23), for convenience, the outliers were removed from the figure.

The results of the long-term forecasts are given in Appendix A5. These results highlight that, current *F_target_* reference points for WBSS herring do not allow rebuilding of the stock above *B_lim_* for the current low recruitment levels, even if *F_target_* = *F_lower_* (Figure A5.1). Rebuilding is however possible with relatively high probability when a stock-recruitment relationship is used in the forecast (Figure A5.2). For North Sea cod, rebuilding above *B_lim_* is possible for all *F_target_* values and recruitment scenarios, but rebuilding probability is improved when a stock-recruitment relationship is assumed, notably for *F_target_* = *F_upper_* (Figures A5.1 and A5.2). Assuming these two recruitment regimes in the estimation of *F_MSY_* resulted in *F_MSY_* consistent with the current value of 0.31 y*^−^*^1^ for cod but not for herring (Tables A5.1 and A5.2). For herring, either a large *F_MSY_* was estimated coupled with low *B_MSY_* and *MSY* values when recruitment was sampled from the assessment estimates, or a *F_MSY_* of *∼* 0.25 y*^−^*^1^ was estimated, associated with values of *B_MSY_* and *MSY* significantly larger than current stock and catch levels.

## 4. Discussion

This study exhibited large contrasts between the two considered stocks. On one hand, WBSS herring, which has been recommended a zero catch for two years, had difficulty rebuilding in the short-to medium-term despite consequent reduction in fishing mortality compared to current levels. On the other hand, North Sea cod benefited from two years of reduced fishing mortality implied by all HCRs at the beginning of the forecast period, and rebuilt quickly, even when recruitment was assumed to decrease in the future. For both species, all *F_target_* values were below the current 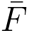 (in last year of assessment) and when *F_target_ ≤ F_MSY_*, below any historical 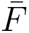. The difference between stocks’ recovery cannot be explained by a difference between current fishing pressure and *F_target_* and other reasons are discussed below.

WBSS herring recovery is compromised with the current reference points used in ICES since while rebuilding above *B_lim_* was possible for the two most pessimistic recruitment assumptions (sampled in 2013-2017 and random walk with negative drift), it was not stable until the end of the forecast period. This oscillating pattern around *B_lim_* is due to the fact that the *F_target_* and *B_lim_* values are inconsistent with the current recruitment regime and this was clearly illustrated by the reference points estimated in this study (Tables A5.1 and A5.2). Reference points for WBSS herring used in the HCRs were estimated using the entire historical SSB and recruitment time series (ICES, 2018c), which included an early period of large productivity, so assuming recruitment stays at low levels or decreases was more pessimistic. On the contrary, when recruitment was assumed to follow a hockey-stick stock-recruitment relationship, herring could rebuild in the medium-term. This is because the hockey-stick recruitment assumption is more in line with the assumptions taken when estimating the reference points used in the HCRs. Indeed, when a hockey-stick was assumed when estimating the MSY reference points in this study, SSB at *F_MSY_* is more than 4 times the SSB in 2018 while it is estimated to be lower than the current SSB by *∼* 35% when recruitment is assumed to stay at low levels (Tables A5.1 and A5.2). Rebuilding above *B_lim_* = 120000 t with current recruitment levels when the corresponding *B_MSY_* is 48400 t is therefore not realistic. For this stock, it would be interesting to investigate what would be the real *F_MSY_* and *B_MSY_* values when a stock recruitment relationship is assumed removing the SSB and recruitment pairs at high values, since the values estimating here corresponds to yield per-recruit estimates rather than MSY reference points. These inconsistencies between current recruitment regimes and reference point estimates explains why rebuilding for North Sea cod occurred more readily. Indeed the reference points for cod were estimated using SSB and recruitment pairs removing the early period of high productivity (ICES, 2015, 2017b), so the period used corresponds already to a relatively low recruitment levels and all recruitment options allowed the stock to recover in the forecast above *B_lim_* and sometimes *MSY B_trigger_* . This is validated by the reference point estimation conducted in this study, where for both recruitment assumptions, *F_MSY_* was estimated around the current *F_MSY_* value of 0.31 y*^−^*^1^ and led to equilibrium SSB values around 1.7-2.9 times the SSB estimated in 2018 (Tables A5.1 and A5.2). In addition, long-term simulations with constant 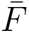 = *F_target_* also showed that the probability of cod rebuilding is mainly improved for *F_target_* = *F_upper_* when the stock-recruitment relationship is used, and assuming current recruitment levels did not compromise cod recovery for the other two *F_target_* values (Figures A5.1 and A5.2). Using favorable recruitment regimes for scientific advice could result in unrealistically optimistic predictions (Caddy and Agnew, 2004) but, we also demonstrate here, that using pessimistic recruitment regimes for scientific advice compared to regimes used for reference point estimation could result in an impossibility to reach biomass reference points. Reference points should therefore be in accordance with current recruitment regimes if they are to be used for practical management advice. This type of medium-term forecasts could therefore highlight possible problems with current reference points that could motivate proposing a benchmark or inter-benchmark to revise them. In practice, when changes in productivity may be difficult to identify or when high productivity estimates are necessary to fit a stock-recruitment relationship used to estimate *F_MSY_* ; simple long-term projections of the stock assuming current recruitment levels for constant 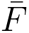 = *F_MSY_* could illustrate possible problems with reference points being inconsistent with current low stock size and recruitment regime (Appendix A5) and should be tested during reference point estimation.

In this study, recruitment was the largest source of uncertainty in future stock response to management as the results were clearly sensitive to the recruitment assumption. For both species, the difference in the recruitment assumption often induced more variability in SSB and cumulative catch than the difference in the HCRs (Figures 6 and 8). This means that the recovery of fish stocks is very dependent on future recruitment. Since recruitment may be difficult to predict, it is therefore very important to test different plausible recruitment assumptions when considering HCRs. Obtaining reliable stock productivity estimates should improve in the future with improvements in predicting oceanic systems in the medium-term (ICES, 2018b), but mean-while, it is relevant to compare optimistic recruitment assumptions with more pessimistic ones to have the full range of worst to best case scenarios with corresponding uncertainty. This could possibly highlight the stability of certain HCRs to recruitment regimes or could be used to present managers with the different stock trajectories obtained for the different recruitment assumptions. For instance, this study shows that for North Sea cod, the recruitment assumption may not limit the stock recovery since rebuilding above *B_lim_* was achieved under all the considered recruitment assumptions. For WBSS herring, however, recruitment led to more variable rebuilding outcomes, notably rebuilding above current targets is compromised if recruitment does not improve in the future. These simulations highlight therefore the necessity to maybe consider exceptional precautionary approaches for this stock if rebuilding is required in the medium-term.

Any management strategies study using HCRs (Kvamsdal et al., 2016; Punt et al., 2016), is limited by the uncertainty in the reference points used (Rice and Rivard, 2007). Reference points are uncertain due to modeling choice and errors but also due to lack of understanding of biological and environmental drivers of the variations in productivity. Also, reference points usually ignore external factors that can affect fish mortality such as environmental factors or multispecies interactions (Trijoulet et al., 2018, 2019, 2020). Reference points such as *F_MSY_* are obtained from long-term equilibrium estimation and are highly dependent on the stock-recruitment relationship used during the estimation process. The stock-recruitment relationship is in itself uncertain and possibly biased as it is usually fitted outside the assessment model without taking into account the variances and covariances in the SSB and recruitment pairs (Brooks and Deroba, 2015; Albertsen and Trijoulet, in press). Also, in ICES, the biomass reference point, *B_lim_*, is either obtained by fitting a hockey-stick relationship to the SSB and recruitment pairs or by a human choice following the ICES guidelines for semi-quantitative estimation of *B_lim_*. In the case of the two species considered in this study, the latter method was used (ICES, 2015, 2018c). In situations where the productivity regime may have changed using historical stock-recruit pairs or the lowest observed biomass to determine *B_lim_* may not be an appropriate way to define rebuilding targets. In other jurisdictions, *B_lim_* is often calculate as a fraction of *B_target_* (e.g. *B_MSY_*) rather than estimated semi-quantitatively (DFO, 2009; ICCAT, 2019). This would provide an objective estimate of *B_lim_* and could be envisaged in ICES, but may still result in a *B_lim_* value difficult to rebuild above, if *B_MSY_* is estimated assuming an optimistic stock recruitment relationship. Also, it would not allow the precautionary constraint in the estimation of MSY reference points (less than 5% chance to fall below *B_lim_*) since in that case *B_lim_* has to be estimated first.

The forecasts account for uncertainty in the current perception of the stock and current fishing pressure by simulating 5000 replicates of numbers at age and fishing mortality at age. However, uncertainty in observations (e.g. weight at age or natural mortality) and model parameters are not considered. The results also do not account for implementation error, which means that the 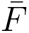 that was obtained from a HCR is the assumed realized 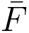 applied on the stock. Exploratory runs were conducted in this study to compare HCR outputs and to highlight trade-offs between fishing opportunities and stock conservation in the medium-term rather than to choose an HCR for real management purpose so the results need to be seen in this context. Comparing HCRs may be difficult to illustrate if implementation and observation errors significantly changes the outputs of the HCRs. How-ever, accounting for implementation error, observation errors and parameter uncertainty will be required if the simulations were used to evaluate robust rebuilding strategies, for instance, for a management plan.

Management decisions can suffer from the uncertainty in stock projections. Managers are often presented with median projection results where the uncertainty or variability in the results could be missing or not well presented. This could result in misunderstandings if the outputs of the forecast are differentt from what is assessed the following year (Kelly et al., 2006). This could also result in a lack of trust in fisheries scientists from managers and stakeholders. It is therefore important to present and discuss uncertainty with managers (Rosenberg and Brault, 1991; Peterman, 2004; Kvamsdal et al., 2016). It is important to compare the outputs and variability of different fishing scenarios or HCRs and quantify the subsequent risk on the stock (Deroba and Bence, 2008; Sethi, 2010). Here, we notably show that HCRs that allowed large variations in 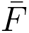 around *B_lim_* were usually less stable (e.g. HCR 1), notably in the short-term. This is due to the low size of the stocks in the first years of forecasts that resulted in large variations in 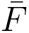 depending on where the simulated stock biomass fell compared to the biomass thresholds for each replicate. As the stocks approached *MSY B_trigger_*, less variations were observed among the HCRs as the shape of the HCRs mainly differed below *B_lim_* (Figure 1).

Managers and fishers have usually a preference for small variations in TACs (Kell et al., 2006; Kelly et al., 2006). A decrease in TACs could threaten fishers, livelihood, but an increase in fishing mortality could also mean higher costs such as for fuel or capacity, or difficult to increase current production. Some studies investigated management rules where the variability in catch was bounded so that the fishing mortality couldnt decrease or increase as rapidly as the variations in SSB (Kell et al., 2006; Kelly et al., 2006). Both studies concluded that, for stocks that are already low, reducing the variability in catch in the forecasts induced a longer time for stock rebuilding and a higher risk on the stock. This was also observed here, notably for North Sea cod where, in the short- (2022) and early medium-term (2025), keeping the 2019 catch constant resulted in smaller projected SSB and higher risk on the stock compared to the HCRs no matter the recruitment assumption (Figure 8). However, this is mainly due to the fact that all HCRs resulted in a decrease in catches compared to 2019 at the beginning of the forecast. The higher 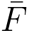 associated with the 2019 catch is therefore less precautionary. We show that applying the MSY approach (HCR 1), or any other HCR that induces a steep decrease in 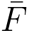 when SSB decreases below *B_lim_* (e.g. HCRs 3 and 5), resulted in large variations in 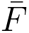 and in subsequent catch no matter the recruitment assumption. This happened in the transition period for cod but over the entire forecast period for WBSS herring. From a management point of view, the catch resulting from these HCRs are less likely to be applied in practice as a smoother change in 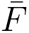 may be preferred by managers.

Stock recovery may necessitate prompt management actions (Caddy and Agnew, 2004; Shertzer and Prager, 2006). Shelton and Morgan (2014) evoke the problem of lag between fish stock assessment and TAC implementation as a reason for the possible lack of success in stock rebuilding when fishing mortality is not reduced fast enough so stock perception changes faster than managers’ response. This raises the question of when it is best to act. The case of WBSS herring can easily illustrate this problem. The perception of the stock changed in 2017 and resulted in a benchmark for the stock a year later (ICES, 2018c). Since then, ICES has recommended a zero catch advice which was never implemented and a rebuilding plan which was never followed up by a request from managers (ICES, 2019i, 2018a). This study shows that recovery for the stock is compromised and will be dependent on future recruitment. Ideally, even if managers react before the stock falls below its biomass threshold, significant changes in stock perception can still occur independently of management, notably after benchmarks where new data, models or configurations are often implemented. This motivates the importance of reacting promptly to changes in stocks, maybe already when the stock approaches *B_lim_*, instead of waiting for the stock to fall below it. The ICES harvest rule with decreasing 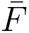 from *F_MSY_* to zero for SSB below *MSY B_trigger_* induces larger 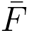 below *MSY B_trigger_* than other frameworks that assume that 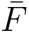 decreases from *F_target_* at the biomass trigger reference point to a reduced 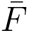 or zero at *B_lim_* and below it (NAFO, 2004; New Zealand Ministry of Fisheries, 2011; ICCAT, 2019). Also *F_target_* is *F_MSY_* in ICES, but other jurisdictions use a *F_target_* below *F_MSY_* as a precautionary approach to avoid falling below *B_MSY_* for reasons such as uncertainty in *B_MSY_* and difficulty to keep the realized 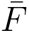 at *F_MSY_* (e.g. NAFO, 2004; NRC, 2014; ICCAT, 2019). Harvest rules with *F_target_ < F_MSY_* induce therefore lower 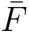 for any SSB value than the ICES harvest rule. Moreover, given the way *MSY B_trigger_* is estimated in ICES, its value is often close to *B_lim_*. This short distance and high 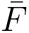 below *MSY B_trigger_* provides therefore less protection to the stock falling below *B_lim_*. According to NRC, 2014, HCRs that reduce fishing mortality promptly and smoothly when SSB is below *B_MSY_* can reduce the probability of the stocks falling below their limit reference point. Future research in ICES should look into HCRs where 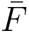 is reduced as soon as SSB is below a higher trigger reference point such as *B_MSY_*, or a fraction of it, rather than below a lower value such as *MSY B_trigger_* . Simulation evaluations using different trigger points could be used in the future to highlight the trade-offs between the level of risk of falling below *B_lim_*, catch levels and variability in catches.

The risk on WBSS herring, measured as the probability of falling below *B_lim_*, was only below 5% at the end of the forecast period when an optimistic recruitment (hockey-stick) was assumed (Figures 6, A4.6, A4.14 and A4.22). For the other two recruitment regimes, even if SSB could increase in the short-term, probability of falling below *B_lim_* was high and even low values of *F_target_* only allowed the probability to decrease to between 20 and 50%. When probability decreased, it often oscillated back to larger values after a few years, reducing the probability of stable recovery for the stock to the end of the forecast. With the current reference point estimates, it seems therefore difficult if not impossible, to reduce the risk on herring if recruitment does not improve in the future, whereas it is possible to have a risk below 5% for cod in the short-term as long as the recruitment does not decrease in the future (random walk with negative drift assumption, Figures 8, A4.10, A4.18 and A4.26). The risk threshold implied by the ICES MSY approach may therefore not be relevant for stocks that do not rebuild in short-term advice and a higher risk may be acceptable (Kell et al., 2006). For these stocks, as it is sometimes done in other jurisdictions (New Zealand Ministry of Fisheries, 2011; NRC, 2014; Australian Government, 2018; ICCAT, 2019), it may therefore be necessary for scientists and managers to agree on what an acceptable deviation from the 5% risk on the stock is in the short-to medium-term, while the low risk level could still be considered in the long-term. For stocks that are low and cannot rebuild in short-term forecasts used for advice (e.g. WBSS herring in this study), the time frame for the evaluation of management strategies has also to be realistic as recovery may be slower than expected and could necessitate looking at medium-term horizons rather than the common short-term advice time frames (Rosenberg et al., 2006; Gröger et al., 2007; Shelton and Morgan, 2014).

Previous studies showed that pelagic stocks can recover after fishery closure if conditions are favorable (Gullestad et al., 2018; Dickey-Collas et al., 2010). According to Murawski (2010), success of rebuilding plans in Canada and in the US usually follows a large decrease in fishing mortality at the beginning of the management years. In this study, for both cod and herring, assuming a HCR where 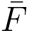 varies with SSB resulted in a decrease in 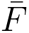 at the beginning of the forecast for all HCRs. It is therefore difficult but necessary to find a balance between HCRs that ensure both stock sustainability and maintaining fishers’ livelihood in the short-term. These considerations are acknowledged in other jurisdictions (e.g. DFO, 2009; New Zealand Ministry of Fisheries, 2011; ICCAT, 2019). In the US, catch agreed by managers cannot exceed the scientific catch advice given by the Scientific and Statistical Committees (U.S. Government, 2007; NRC (National Research Council), 2014), but this is not the case in other jurisdictions including in ICES. Currently scientific advice from ICES for depleted stocks without adopted management plans is solely given to ensure stock sustainability and this explains why there is often a discrepancy between what is recommended by scientists and what is applied by managers. In this study, we show that, if cod recruitment does not decrease in the future, the median cumulative catch in the medium-term can be similar and the risk on the stock low for all HCRs, demonstrating that a smooth change in 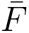 is a reasonable option for ensuring a sustainable fishery on the stock. For WBSS herring, HCRs 1 and 3 resulted in similar median cumulative catch and risk on the stock in the medium-term (Figure 6) but HCR 1 closed the fishery for 3 consecutive years whereas HCR 3 allowed for a low 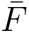 for 4 consecutive years. It is therefore possible to keep fishing in the short-term for the same trade-offs in the medium-term. A lower *F_target_* (*F_lower_*) also improved the probability of rebuilding for both species for often little concession in median cumulative catch. For depleted stocks, delivering advice on scenarios that ensure stock rebuilding in medium-term rather than in the short-term while maintaining fishing over time may therefore be more appropriate than using the MSY approach.

This type of medium-term forecasts would be relevant for stocks in need of rebuilding plans (Garcia et al., 2018). They could be used to guide development of rebuilding plan requests. For instance, to define realistic rebuilding objectives for specific stocks such as rebuilding targets, rebuilding time frame, rebuilding probability or HCRs. In ICES and many other jurisdictions, management strategy evaluations (MSE) are usually used to choose a fishing scenarios or HCR that will be used as advice within a management plan. However, MSEs suffer from complex modeling implementations and need of extensive computer power (Punt et al., 2016; ICES, 2019n). This usually results in outcomes not being available before months or years after a rebuilding plan evaluation is requested. When zero catch is advised for a stock and no management plan is already in place, medium-term forecasts could be used as a platform to illustrate the trade-offs between rebuilding time, rebuilding probability, stock trajectories and predicted catches to provide quick guidance to managers during scientific advice or quota allocation, notably when zero catch cannot be realistically implemented.

## Acknowledgments

This work was funded by a grant from the European Fisheries and Maritime Fund (33113-B-17-106, Ministry of Environment and Food in Denmark). The authors would like to thank the members of the ICES 2019 Herring Assessment Working Group for the Area South of 62*^◦^*N (HAWG) for showing interest in the current study, which led to interesting discussions and motivated writing this manuscript. We are also grateful to Andre E. Punt and David Miller for comments on an early version of this manuscript.

## Footnotes

1 Process held every 3-5 years where the data, the model and its configuration are agreed on, the stock is assessed, and reference points are estimated. Inter-benchmarks can be used for minor changes such as model configurations or data modifications. See ICES (2019a) for more detail.

2 In ICES, *MSY B_trigger_* is usually estimated as the 5*^th^* percentile of the spawning stock biomass (SSB) when fished at *F_MSY_*, but, when the variations in SSB do not allow for the 5*^th^* percentile to be estimated (e.g. the stock was not fished at *F_MSY_* for at least 5 years), it can also be set as the 95*^th^* percentile (*B_pa_*) of the biomass below which the stock is considered to have a reduced reproductive capacity (*B_lim_*.) (ICES, 2017a).

3 Spawning stock biomass limit reference point used in ICES under which the stock is believed to have impaired recruitment.

4 In mixed fisheries, stock that has the lowest quota and, due to the Landing Obligation, forces a fleet to stop fishing even if it still has quotas for other stocks.

5 *F_lower_* and *F_upper_* are the mean of the lowest and highest values respectively within the range of *F_MSY_* . The range is estimated to obtain no more than a 5% reduction in long-term yield compared to MSY. Similarly than for *F_MSY_*, the range of *F_MSY_* is restricted to the extra precautionary consideration of less than 5% chance of SSB falling below *B_lim_*. See for example Chapter 1, Article 2 of European Commission (2018a) for more detail.

## References

Albertsen, C.M., Trijoulet, V., in press. Model-based estimates of reference points in an age-based state-space stock assessment model. Fisheries Research .

Australian Government, 2018. Guidelines for the Implementation of the Commonwealth Fisheries Harvest Strategy Policy. Technical Report. Department of Agriculture and Water Resources.

Barrowman, N.J., Myers, R.A., 2000. Still more spawner-recruitment curves: the hockey stick and its generalizations. Canadian Journal of Fisheries and Aquatic Sciences 57, 665–676. doi:10.1139/f99-282.

Berg, C.W., Nielsen, A., 2016. Accounting for correlated observations in an age-based state-space stock assessment model. ICES Journal of Marine Science 73, 1788–1797. doi:10.1093/icesjms/fsw046.

Brooks, E.N., Deroba, J.J., 2015. When “data” are not data: the pitfalls of post hoc analyses that use stock assessment model output. Canadian Journal of Fisheries and Aquatic Sciences 72, 634–641. doi:10.1139/cjfas-2014-0231, arXiv:https://doi.org/10.1139/cjfas-2014-0231.

Caddy, J.F., Agnew, D.J., 2004. An overview of recent global experience with recovery plans for depleted marine resources and suggested guidelines for recovery planning. doi:10.1007/s11160-004-3770-2.

Cadrin, S.X., Pastoors, M.A., 2008. Precautionary harvest policies and the uncertainty paradox. Fisheries Research doi:10.1016/j.fishres.2008.06.004.

Deroba, J.J., Bence, J.R., 2008. A review of harvest policies: Understanding relative performance of control rules. Fisheries Research 94, 210–223. URL: http://www.sciencedirect.com/science/article/pii/S0165783608000192, doi:10.1016/j.fishres.2008.01.003.

DFO (Fisheries and Oceans Canada), 2009. A fishery decision-making framework incorporating the precautionary approach. URL: http://www.dfo-mpo.gc.ca/reports-rapports/regs/sff-cpd/precaution-eng.htm.

Dickey-Collas, M., Nash, R.D., Brunel, T., Van Damme, C.J., Marshall, C.T., Payne, M.R., Corten, A., Geffen, A.J., Peck, M.A., Hatfield, E.M., Hintzen, N.T., Enberg, K., Kell, L.T., Simmonds, E.J., 2010. Lessons learned from stock collapse and recovery of North Sea herring: A review. doi:10.1093/icesjms/fsq033.

EU, 2013. Regulation (EU) No 1380/2013 of the European Parliament and of the Council of 11 December 2013 on the Common Fisheries Policy, amending Council Regulations (EC) No 1954/2003 and (EC) No 1224/2009 and repealing Council Regulations (EC) No 2371/2002 and (EC) No 639/2004 and Council Decision 2004/585/EC. Technical Report. Official Journal of the European Union.

EU, 2017. Council Regulation (EU) 2017/1970 of 27 October 2017 fixing for 2018 the fishing opportunities for certain fish stocks and groups of fish stocks applicable in the Baltic Sea and amending Regulation (EU) 2017/127. Official Journal of the European Union URL: https://eur-lex.europa.eu/legal-content/EN/TXT/?uri=CELEX:32017R1970.

EU, 2018a. Council Regulation (EU) 2018/120 of 23 January 2018 fixing for 2018 the fishing opportunities for certain fish stocks and groups of fish stocks, applicable in Union waters and, for Union fishing vessels, in certain non-Union waters, and amending Regulation (EU) 2017/127. Official Journal of the European Union URL: https://eur-lex.europa.eu/legalcontent/en/TXT/?uri=CELEX:32018R0120.

EU, 2018b. Council Regulation (EU) 2018/1628 of 30 October 2018 fixing for 2019 the fishing opportunities for certain fish stocks and group of fish stocks applicable in the Baltic Sea and amending Regulation (EU) 2018/120 as regards certain fishing opportunities in. Official Journal of the European Union URL: https://eur-lex.europa.eu/legal-content/en/TXT/?uri=CELEX:32018R1628.

EU, 2019a. Council Regulation (EU) 2019/124 of 30 January 2019 fixing for 2019 the fishing opportunities for certain fish stocks and groups of fishing stocks, applicable in Union waters and, for Union fishing vessels, in certain non-Union waters. Official Journal of the European Union URL: https://eur-lex.europa.eu/legal-content/EN/TXT/?uri=uriserv:OJ.L{\_}.2019.029.01.0001.01.ENG.

EU, 2019b. COUNCIL REGULATION (EU) 2019/1838 of 30 October 2019 fixing for 2020 the fishing opportunities for certain fish stocks and groups of fish stocks applicable in the Baltic Sea and amending Regulation (EU) 2019/124 as regards certain fishing opportunities in other waters. Official Journal of the European Union, 14.

EU, Norway, 2019. Agreed record of conclusions of fisheries consultations between Norway and the European Union on the regulation of fisheries in Skagerrak and Kattegat for 2020. URL: https://ec.europa.eu/fisheries/sites/fisheries/files/docs/body/2020-norway-fisheries-consultations-skagerrak-kattegat{\_}en.pdf.

EU, Norway, 2020. Agreed record of fisheries consultations between the European Union and Norway for 2020. URL: https://ec.europa.eu/fisheries/sites/fisheries/files/docs/body/2020-norway-fisheries-consultations-north-sea{\_}en.pdf.

European Commission, 2018a. Proposal for a REGULATION OF THE EUROPEAN PARLIAMENT AND OF THE COUNCIL establishing a multiannual plan for fish stocks in the Western Waters and adjacent waters, and for fisheries exploiting those stocks, amending Regulation (EU) 2016/1139 establishing a multiannual plan for the Baltic Sea, and repealing Regulations (EC) No 811/2004, (EC) No 2166/2005, (EC) No 388/2006, (EC) 509/2007 and (EC) 1300/2008. URL: https://eur-lex.europa.eu/legal-content/en/TXT/?uri=CELEX:52018PC0149.

European Commission, 2018b. Proposal for a REGULATION OF THE EUROPEAN PARLIAMENT AND OF THE COUNCIL establishing a multiannual plan for fish stocks in the Western Waters and adjacent waters, and for fisheries exploiting those stocks, amending Regulation (EU) 2016/1139 establishing a multiannual plan for the Baltic Sea, and repealing Regulations (EC) No 811/2004, (EC) No 2166/2005, (EC) No 388/2006, (EC) 509/2007 and (EC) 1300/2008. URL: https://http://data.europa.eu/eli/reg/2018/973/oj.

FAO, 2018. The state of world fisheries and agriculture 2018 - meeting the sustainable development goals. Rome. Licence: CC BY-NC-SA 3.0 IGO. URL: http://www.fao.org/3/i9540en/i9540en.pdf.

Garcia, S.M., 1994. The Precautionary Principle: its implications in capture fisheries management. Ocean & Coastal Management 22, 99–125. URL: http://www.sciencedirect.com/science/article/pii/0964569194900140, doi:https://doi.org/10.1016/0964-5691(94) 90014-0.

Garcia, S.M., Ye, Y., Rice, J., Charles, A., 2018. Rebuilding of marine fisheries Part 1: Global review. 630/1, Food and Agriculture Organization of the United Nations. URL: http://www.fao.org/3/ca0161en/CA0161EN.pdf.

Gröger, J.P., Rountree, R.A., Missong, M., Rätz, H.J., 2007. A stock rebuilding algorithm featuring risk assessment and an optimization strategy of single or multispecies fisheries ICES Journal of Marine Science 64, 1101–1115. doi:10.1093/icesjms/fsm085.

Gullestad, P., Howell, D., Stenevik, E.K.and Sandberg, P., Bakke, G., 2018. Management and rebuilding of herring and cod in the Northeast Atlantic, in: Rebuilding of marine fisheries Part 2: Case studies.. FAO Fisheries and Aquaculture Technical paper, Rome, pp. 12–37.

Hilborn, R., Ovando, D., 2014. Reflections on the success of traditional fisheries management. ICES Journal of Marine Science 71, 1040–1046. doi:10.1093/icesjms/fsu034.

ICCAT (International Commission for the Conservation of Atlantic Tunas), 2019. Compendium of the Management Recommendations and Resolutions adopted by ICCAT for the Conservation of Atlantic Tunas and Tuna-Like Species. Technical Report.

ICES, 2015. Report of the Benchmark Workshop on North Sea Stocks (WKNSEA). Technical Report. 2-6 February 2015, Copenhagen, Denmark. ICES CM 2015/ACOM:32.

ICES, 2017a. ICES fisheries management reference points for category 1 and 2 stocks. Technical Report. ICES Advice Technical Guidelines, ICES Advice 2017, Book 12. doi:10.17895/ices.pub.3036.

ICES, 2017b. Report of the Working Group on Assessment of Demersal Stocks in the North Sea and Skagerrak (2017). Technical Report. 26 April-5 May 2017, ICES HQ. ICES CM 2017/ACOM:21.

ICES, 2018a. Herring (Clupea harengus) in subdivisions 20–24, spring spawners (Skagerrak, Kattegat, and western Baltic). Technical Report. ICES Advice on fishing opportunities, catch, and effort Baltic Sea and Greater North Sea Ecoregions. Report of the ICES Advisory Committee 2018. ICES Advice 2018. doi:10.17895/ices.pub.4390.

ICES, 2018b. Interim Report of the Working Group on Seasonal to Decadal Prediction of Marine Ecosystems (WGS2D). Technical Report. 27–31 August 2018, ICES Headquarters, Copenhagen, Denmark. ICES CM 2018/EPDSG:22.

ICES, 2018c. Report of the benchmark workshop on pelagic stocks (WKPELA 2018). Technical Report. 12–16 February 2018, ICES HQ, Copenhagen, Denmark. ICES CM 2018/ACOM:32.

ICES, 2019a. Advice basis. Technical Report. Report of the ICES Advisory Committee. ICES Advice 2019. doi:10.17895/ices.advice.5757.

ICES, 2019b. Baltic Sea Ecoregion – Fisheries overview. Technical Report. ICES Advice 2019, ICES Fisheries Overviews, Baltic Sea Ecoregion. URL: http://www.ices.dk/sites/pub/Publication%20Reports/Advice/2019/2019/BalticSeaEcoregion_FisheriesOverviews.pdf.

ICES, 2019c. Barents Sea Ecoregion – Fisheries overview. Technical Report. ICES Advice 2019, ICES Fisheries Overviews, Barents Sea Ecoregion. URL: http://www.ices.dk/sites/pub/Publication%20Reports/Advice/2019/2019/FisheriesOverview_BarentsSea_2019.pdf, doi:10.17895/ices.advice.5705.

ICES, 2019d. Bay of Biscay and Iberian Coast ecoregion – Fisheries overview, including mixed-fisheries considerations. Technical Report. ICES Advice 2019, ICES Fisheries Overviews, Bay of Biscay and Iberian Coast ecoregion. URL: http://www.ices.dk/sites/pub/Publication%20Reports/Advice/2019/2019/FisheriesOverviews_BoBIberian_2019.pdf, doi:10.17895/ices.advice.5709.

ICES, 2019e. Celtic Seas ecoregion – Fisheries overview, including mixed-fisheries considerations. Technical Report. ICES Advice 2019, ICES Fisheries Overviews, Celtic Seas ecoregion. URL: http://www.ices.dk/sites/pub/Publication%20Reports/Advice/2019/2019/FisheriesOverviews_CelticSeas_2019.pdf, doi:10.17895/ices.advice.5708.

ICES, 2019f. Cod (Gadus morhua) in Subarea 4, Division 7.d, and Subdivision 20 (North Sea, eastern English Channel, Skagerrak). Technical Report. ICES Advice on fishing opportunities, catch, and effort Greater North Sea Ecoregions. Report of the ICES Advisory Committee 2019. ICES Advice 2019.

ICES, 2019g. Greater North Sea Ecoregion - Fisheries overview, including mixed-fisheries considerations. Technical Report. ICES Advice 2019, ICES Fisheries Overviews Great North Sea Ecoregion. doi:10.17895/ices.advice.5710.

ICES, 2019h. Greater North Sea Ecoregion – Fisheries overview, including mixed-fisheries considerations. Technical Report. ICES Advice 2019, ICES Fisheries Overviews, Great North Sea Ecoregion. URL: http://www.ices.dk/sites/pub/Publication%20Reports/Advice/2019/2019/FisheriesOverview_GreaterNorthSea_2019.pdf, doi:10.17895/ices.advice.5710.

ICES, 2019i. Herring (Clupea harengus) in subdivisions 20–24, spring spawners (Skagerrak, Kattegat, and western Baltic). Technical Report. ICES Advice on fishing opportunities, catch, and effort Baltic Sea and Greater North Sea Ecoregions. Report of the ICES Advisory Committee 2019. ICES Advice 2019.

ICES, 2019j. Icelandic Waters ecoregion – Fisheries overview. Technical Report. ICES Advice 2019, ICES Fisheries Overviews, Icelandic Waters ecoregion. URL: http://www.ices.dk/sites/pub/Publication%20Reports/Advice/2019/2019/FisheriesOverview_IcelandicWaters_2019.pdf, doi:10.17895/ices.advice.5706.

ICES, 2019k. Norwegian Sea ecoregion – Fisheries overview. Technical Report. ICES Advice 2019, ICES Fisheries Overviews, Norwegian Sea ecoregion. URL: http://www.ices.dk/sites/pub/Publication%20Reports/Advice/2019/2019/FisheriesOverviews_Norwegian%20Sea_2019.pdf, doi:10.17895/ices.advice.5706.

ICES, 2019l. Report of the Herring Assessment Working Group for the Area South of 62°N (HAWG). Technical Report. 23–31 January 2019 and 13-21 March 2019. ICES HQ, Copenhagen, Denmark.

ICES, 2019m. Working Group on the Assessment of Demersal Stocks in the North Sea and Skagerrak (WGNSSK). Technical Report. ICES Scientific Reports. 1:7. doi:10.17895/ices.pub.5402.

ICES, 2019n. WORKSHOP ON NORTH SEA STOCKS MANAGEMENT STRATEGY EVALUATION (WKNSMSE). Technical Report 12. ICES Scientific Reports. URL: http://www.ices.dk/sites/pub/PublicationReports/ExpertGroupReport/acom/2019/WKNSMSE/ICESWKNSMSEReport2019.pdf, doi:10.17895/ices.pub.5090.

Kell, L.T., Pilling, G.M., Kirkwood, G.P., Pastoors, M.A., Mesnil, B., Korsbrekke, K., Abaunza, P., Aps, R., Biseau, A., Kunzlik, P., Needle, C.L., Roel, B.A., Ulrich, C., 2006. An evaluation of multi-annual management strategies for ICES roundfish stocks. ICES Journal of Marine Science 63, 12–24. doi:10.1016/j.icesjms.2005.09.003.

Kelly, C.J., Codling, E.A., Rogan, E., 2006. The Irish Sea cod recovery plan: some lessons learned. ICES Journal of Marine Science 63, 600–610. doi:10.1016/j.icesjms.2005.12.001.

Kvamsdal, S.F., Eide, A., Ekerhovd, N.A., Enberg, K., Gudmundsdottir, A., Hoel, A.H., Mills, K.E., Mueter, F.J., Ravn-Jonsen, L., Sandal, L.K., Stiansen, J.E., Vestergaard, N., 2016. Harvest control rules in modern fisherries managementharvest control rules in modern fisheries management. Elementa: Science of the Anthropocene 4, 1–22. doi:10.12952/journal.elementa.000114.

Murawski, S.A., 2010. Rebuilding depleted fish stocks: The good, the bad, and, mostly, the ugly. ICES Journal of Marine Science doi:10.1093/ icesjms/fsq125.

NAFO (Northwest Atlantic Fisheries Organization), 2004. NAFO Precautionary Approach Framework, NAFO/FC Doc. 04/18, Serial N5069. Technical Report. URL: https://archive.nafo.int/open/fc/2004/fcdoc04-18.pdf.

New Zealand Ministry of Fisheries, 2011. Operational guidelines for New Zealand’s harvest strategy standard. Technical Report.

Nielsen, A., Berg, C.W., 2014. Estimation of time-varying selectivity in stock assessments using state-space models. Fisheries Research 158, 96–101. URL: http://www.sciencedirect.com/science/article/pii/S0165783614000228, doi:10.1016/j.fishres.2014.01.014.

NRC (National Research Council), 2014. Evaluating the Effectiveness of Fish Stock Rebuilding Plans in the United States. The National Academies Press, Washington, DC. URL: https://www.nap.edu/catalog/18488/evaluating-the-effectiveness-of-fish-stock-rebuilding-plans-in-the-united-state doi:10.17226/18488.

Peterman, R.M., 2004. Possible solutions to some challenges facing fisheries scientists and managers. ICES Journal of Marine Science 61, 1331–1343. doi:10.1016/j.icesjms.2004.08.017.

Punt, A.E., Butterworth, D.S., de Moor, C.L., De Oliveira, J.A.A., Haddon, M., 2016. Management strategy evaluation: best practices. Fish and Fisheries 17, 303–334. doi:10.1111/faf.12104.

Rice, J.C., Rivard, D., 2007. The dual role of indicators in optimal fisheries management strategies. ICES Journal of Marine Science 64, 775–778. doi:10.1093/icesjms/fsm033.

Richards, L.J., Maguire, J.J., 1998. Recent international agreements and the precautionary approach: new directions for fisheries management science. Canadian Journal of Fisheries and Aquatic Sciences 55, 1545–1552. doi:10.1139/f98-043.

Rosenberg, A.A., Brault, S., 1991. Stock Rebuilding Strategies Over Different Time Scales. NAFO Sci. Coun. Studies 16, 171–181.

Rosenberg, A.A., Swasey, J.H., Bowman, M., 2006. Rebuilding US fisheries progress and problems. Frontiers in Ecology and the Environment 4, 303– 308. doi:10.1890/1540-9295(2006)4[303:RUFPAP]2.0.CO;2.

Sethi, S.A., 2010. Risk management for fisheries Fish and Fisheries 11, 341–365. URL: https://onlinelibrary.wiley.com/doi/abs/10.1111/j.1467-2979.2010.00363.x, doi:10.1111/j.1467-2979.2010.00363.x.

Shelton, P.A., Morgan, M.J., 2014. Impact of maximum sustainable yield-based fisheries management frameworks on rebuilding North Atlantic cod stocks. Journal of Northwest Atlantic Fishery Science 46, 15–25. doi:10.2960/j.v46.m697.

Shertzer, K.W., Prager, M.H., 2006. Delay in fisheries management: diminished yield, longer rebuilding, and increased probability of stock collapse1. ICES Journal of Marine Science 64, 149–159. doi:10.1093/icesjms/fsl005.

Trijoulet, V., Fay, G., Curti, K., Smith, B., Miller, T.J., 2019. Performance of multispecies population models: insights on the influence of diet data. ICES Journal of marine Science doi:10.1093/icesjms/fsz053.

Trijoulet, V., Fay, G., Miller, T.J., 2020. Performance of a state-space multi-species model: What are the consequences of ignoring predation and process errors in stock assessments? Journal of Applied Ecology 57, 121–135. doi:10.1111/1365-2664.13515.

Trijoulet, V., Holmes, S.J., Cook, R.M., 2018. Grey seal predation mortality on three depleted stocks in the West of Scotland: What are the implications for stock assessments? Canadian Journal of Fisheries and Aquatic Sciences 75, 723–732. doi:10.1139/cjfas-2016-0521.

UN (United Nations), 1995. Agreement for the implementation of the provisions of the United Nations convention on the law of the sea of 10 December 1982 relating to the conservation and management of straddling fish stocks and highly migratory fis stocks. URL: https://www.un.org/Depts/los/convention{\_}agreements/convention{\_}overview{\_}fish{\_}stocks.htm.

U.S. Government, 2007. Magnuson-Stevens Fishery Conservation and Management Act. Technical Report. URL: https://www.fisheries.noaa.gov/resource/document/magnuson-stevens-fishery-conservation-and-management-act.

Wetzel, C.R., Punt, A.E., editor: Emory Anderson, H., 2016. The impact of alternative rebuilding strategies to rebuild overfished stocks. ICES Journal of Marine Science 73, 2190–2207. URL: https://doi.org/10.1093/icesjms/fsw073, doi:10.1093/icesjms/fsw073.

