## Supplementary material for "Can we fish on stocks that need rebuilding? Illustrating the trade-offs between stock conservation and fisheries considerations"

#### A1 Recruitment assumptions in the forecast

The two assumptions of recruitment sampled from specific assessment estimates and recruitment following a random walk with negative drift are illustrated in Figures A1.1 and A1.2 for 5000 replicates. For both assumptions, recruitment is not affected by a change in stock size and is therefore similar for all MSs tested.

The fit of the hockey-stick stock-recruitment relationship is given in Figure A1.3 for both species. The fitted curve and corresponding uncertainty were used in the forecast to simulated recruitment as a function of spawning stock biomass (SBB).

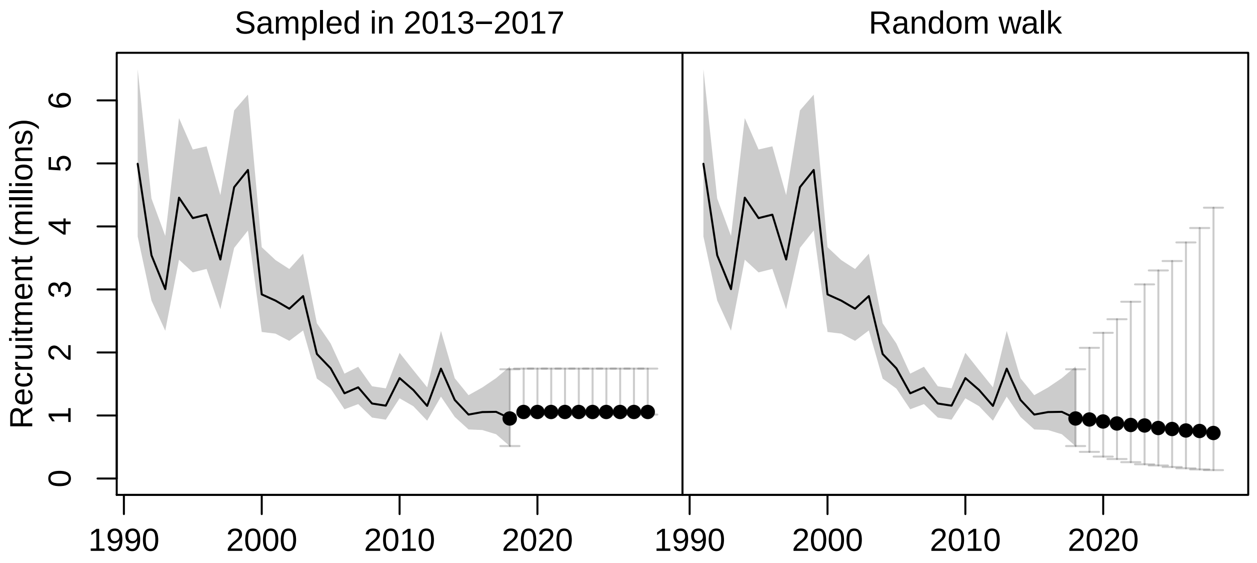


Figure A1.1: Recruitment assumptions for WBSS herring. The points are the medians and the segments the 95% confidence intervals.

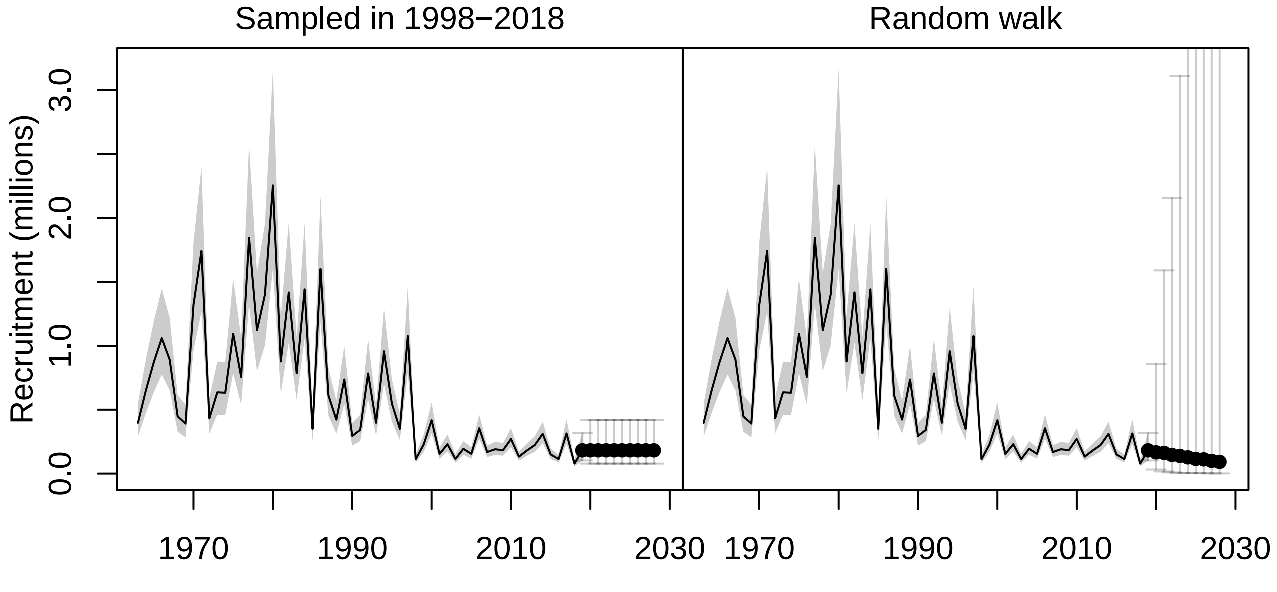


Figure A1.2: Recruitment assumptions for North Sea cod. The points are the medians and the segments the 95% confidence intervals. The upper limit of the confidence interval for the random walk assumption is not shown as it reaches quickly unrealistic levels.

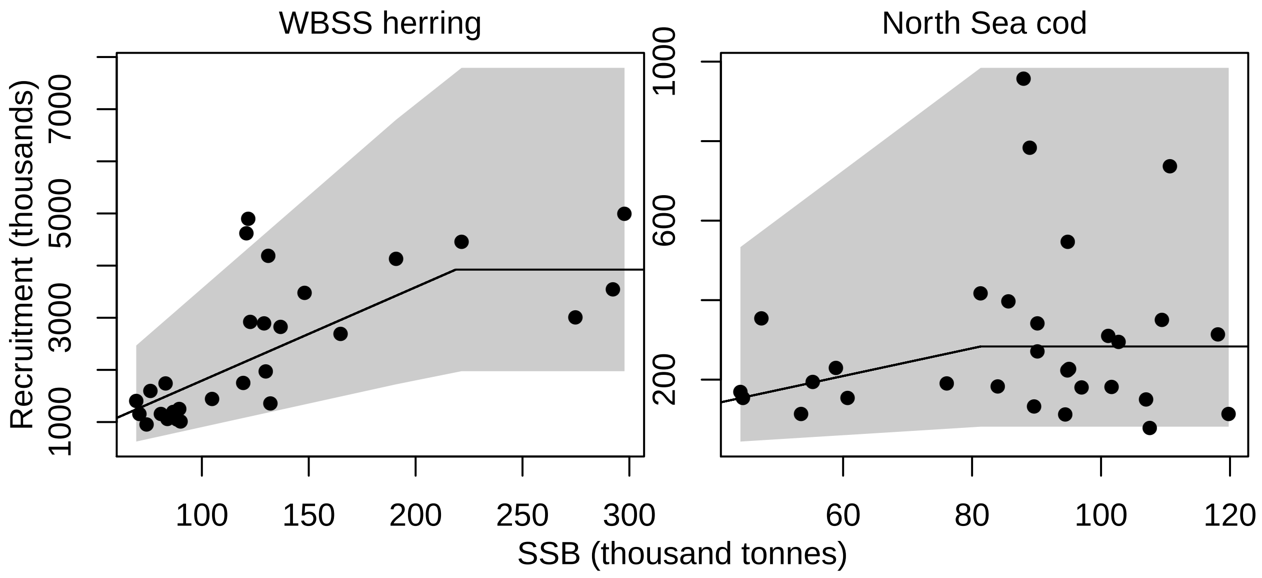


Figure A1.3: Fit of the hockey-stick stock-recruitment relationship for WBSS herring and North Sea cod. The points are the observations (estimates from the assessment models), the line is the fitted curve and the shaded area is the 95% confidence interval.

#### A2 Intermediate year assumptions in the forecast

Similarly to what is used in forecasts used for advice for WBSS herring and North Sea cod, catch constraints were given in the intermediate year (2019). We used the same assumptions in 2019 as those used in the 2019 advice forecasts (Table A2.1, see ICES (2019b,c) for more details).

Table A2.1: Catch constraint (in tonnes) used in the forecast for the intermediate year (2019). The assessment model for North Sea cod is not disaggregated into several commercial fishing fleet.

| Fleet | WBSS herring | North Sea cod |
|---|---|---|
| A | 1545 | |
| C | 12352 | |
| D | 469 | |
| F | 9001 | |
| All fleets | 23367 | 35358 |

#### A3 Method used for MSY reference point estimation

For both species, long-term forecasts were run for the assumption of recruitment sampled from the assessment estimates and following an hockey-stick relationship. The 5000 forecast replicates were run assuming the same assumption than the medium-term forecasts until the steady-state was reached (30 to 50 years) for different values of fishing mortality with a step of 0.001. The landings in the last year of the forecast was extracted and averaged across replicates. $F_{MSY}$ was then identified as the fishing mortality than maximizes the mean landings. Note than for WBSS herring discards are assumed negligible so landings and catch are the same but this is not the case for North Sea cod. When recruitment is randomly sampled from the assessment estimates and is therefore flat on average, $F_{MSY}$ corresponds to $F$ at maximum yield-per-recruit, $F_{max}$. The forecast for which $F = F_{MSY}$ was used to calculate median SSB at MSY $B_{MSY}$ and its variability (95% confidence interval in the stochastic forecast). The same was done for median landings at MSY and $F_{MSY}$.

#### A4 Results for all simulation trials (STs)

The following figures summarize the results for all STs for WBSS herring and North Sea cod presented in diverse manners. In addition to what is presented in the paper, these highlight also different ways of presenting the diagnostics of this type of studies. For each species and recruitment assumptions, SSB, $\bar{F}$ and total catch forecast trajectories are given. Median total catch against the risk on the stocks is illustrated. Cumulative catch over time with confidence intervals are also given.

#### A4.1 Base cases for WBSS herring and North Sea cod

For both base cases ($F = 0$ and catch constant at the 2019 level), the WBSS herring SSB in the forecasts increased over time for all recruitment assumptions but assuming a random walk with negative drift resulted in a lower median SSB than the two other recruitment assumptions (Figure A4.1). As expected, the uncertainty around the SSB estimates in the forecast increased rapidly when recruitment was assumed to follow a random walk, resulting in a span of SSB values much wider than for the other two assumptions, and skewed towards high SSB values. Given the current perception of the stock, the zero catch option allowed the median SSB to rebuild above $B_{lim}$ by 2022 and above $MSYB_{trigger}$ by 2023 for all recruitment assumptions. By the end of the forecast period, SSB reached levels similar to what was observed in the 1990s.

Keeping the catch constant at the 2019 level did not allow stock rebuilding in the short-term. Assuming recruitment stayed at levels similar to the last five year average (recruitment sampled in 2013-2017) only allowed the stock to recover above $B_{lim}$ on average by 2025, while median rebuilding above $B_{lim}$ starts by 2024 when a hockey-stick is assumed and was not possible under the random walk with negative drift assumption.

North Sea cod recovers above $B_{lim}$ by 2021 following fishing closure for all recruitment assumptions and gets above historical average after only two more years (Figure A4.2). Median SSB is dratiscally increased for both base cases when the recruitment assumption is more optimistic (hockey-stick) than staying at low regime level (sampled in 1998-2018) or decreasing further (random walk with negative drift).

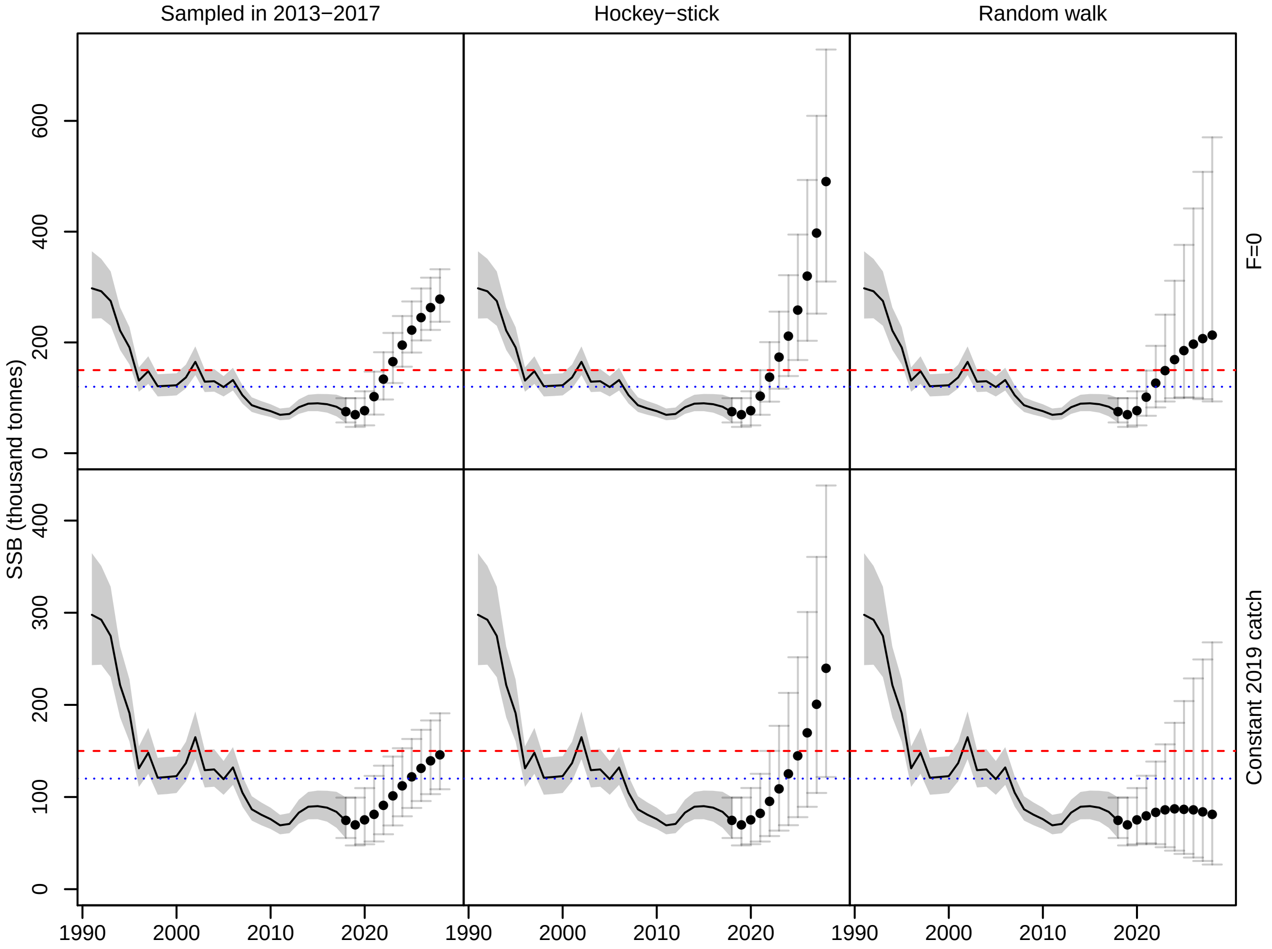


Figure A4.1: WBSS herring spawning stock biomass (SSB) estimated in the base cases for the three recruitment assumptions. The points are the median SSB and the segments the 95% confidence intervals estimated from the 5000 replicates. The red dashed line is $MSYB_{trigger}$ and the blue dotted line is $B_{lim}$.

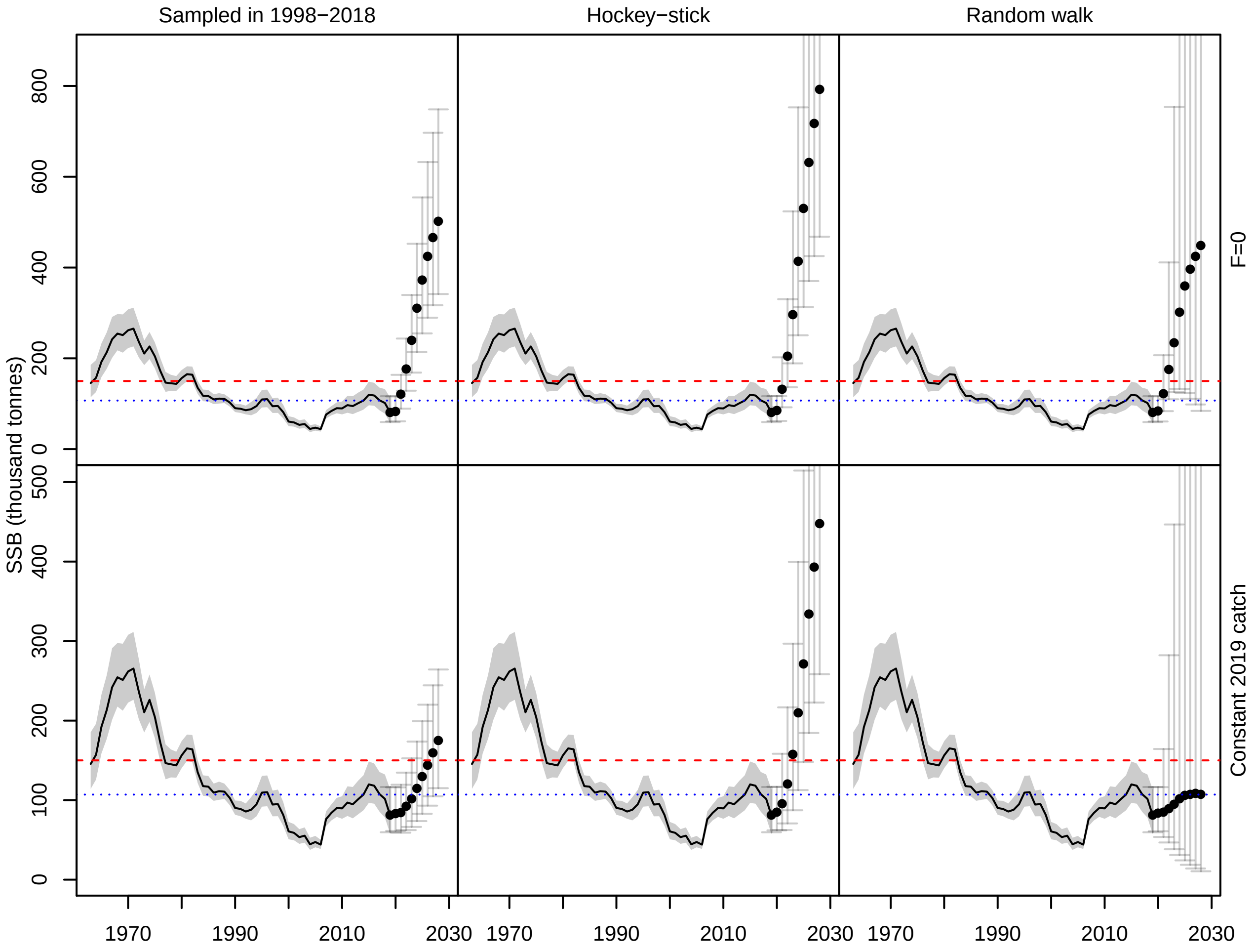


Figure A4.2: North Sea cod spawning stock biomass (SSB) estimated in the base cases for the three recruitment assumptions. The points are the median SSB and the segments the 95% confidence intervals estimated from the 5000 replicates. The red dashed line is $MSYB_{trigger}$ and the blue dotted line is $B_{lim}$. For plotting convenience the upper limits of the confidence intervals are not shown in the hockey-stick and random walk cases.

#### A4.2 STs with HCR

##### A4.2.1 Recruitment sampled from the assessment estimates

**WBSS herring** When recruitment is randomly sampled in 2013-2017 for each replicate, none of the forecast options enabled rebuilding of the stock above $B_{lim}$ by 2021 (Figure A4.3), which is the time frame usually given for ICES advice (ICES, 2019a). Rebuilding was possible from 2022 with the current $F_{target}$ value ($F_{MSY}$) for the HCR 1, and from 2023 for HCRs 3 and 5. However, under the HCRs 1 and 3, median SSB declined again below $B_{lim}$ after a few years while the median SSB stayed above $B_{lim}$ for HCR 5. Reducing $F_{target}$ to $F_{lower}$ resulted for all HCRs in rebuilding the stock to a stable level between $B_{lim}$ and $MSYB_{trigger}$ until the last year of the forecast period (2028). Increasing $F_{target}$ to $F_{upper}$ decreased the probability of stock rebuilding since no HCR resulted in $SSB \geq B_{lim}$ at the end of the forecast, except for the HCR 1 where $F = 0$ when $SSB < B_{lim}$. However, the latter option induced SSB oscillations between values lower than $B_{lim}$ and values around $MSYB_{trigger}$. For all options, there is little probability of further decline of the stock since they all enabled an increase in median SSB compared to the 2018 estimate. None of the HCRs showed a stable recovery of the stock above $MSYB_{trigger}$ and the upper limit of the 95% confidence interval around SSB rarely went above $MSYB_{trigger}$.

The change in $\bar{F}_{3-6}$ is presented in Figure A4.4. Large jumps in $F$ were observed for HCRs 1, 3 and 5 that oscillated between lowest $\bar{F}_{3-6}$ values (0 for HCR 1, 0.1 for HCRs 3 and 5). This induced large changes in total catch, notably for HCRs 1 and 3 (Figure A4.5).

Under the assumption of recruitment randomly sampled in 2013-2017, probability of WBSS herring SSB falling below $B_{lim}$ is high ($> 60\%$) in the first 3 forecast years where management was applied (2020-2022) for all HCRs except HCR 1 for which the probability was around 20% in 2022 but conditional on zero catch for 2020-2022 (Figure A4.6). For HCR 1, catches then increased but decreased again at the end of the forecast period for large values of $F_{target}$ ($F_{MSY}$ and $F_{upper}$). The probability of falling below $B_{lim}$ in the constant 2019 catch option did not decline to less than 50% until 2025. For HCRs 2 and 4 the results were very similar with very low probability of stock rebuilding ($< 20\%$) when $F_{target} = F_{MSY}$ or $F_{upper}$ with no improvements compared to the constant 2019 catch option. Probability of rebuilding with higher yield increased when $F_{target} = F_{lower}$. HCRs 3 and 5 presented similar patterns where the probability of rebuilding and yield increased over time but then decreased after 2024-2026. Median catches were slightly larger for HCR 3. Overall, HCRs 3 and 5 were the HCRs with the lowest probability of decline that still allowed reasonable herring catches. Lower values of $F_{target}$ enabled lower probability of falling below $B_{lim}$ without compromising on the level of catch.

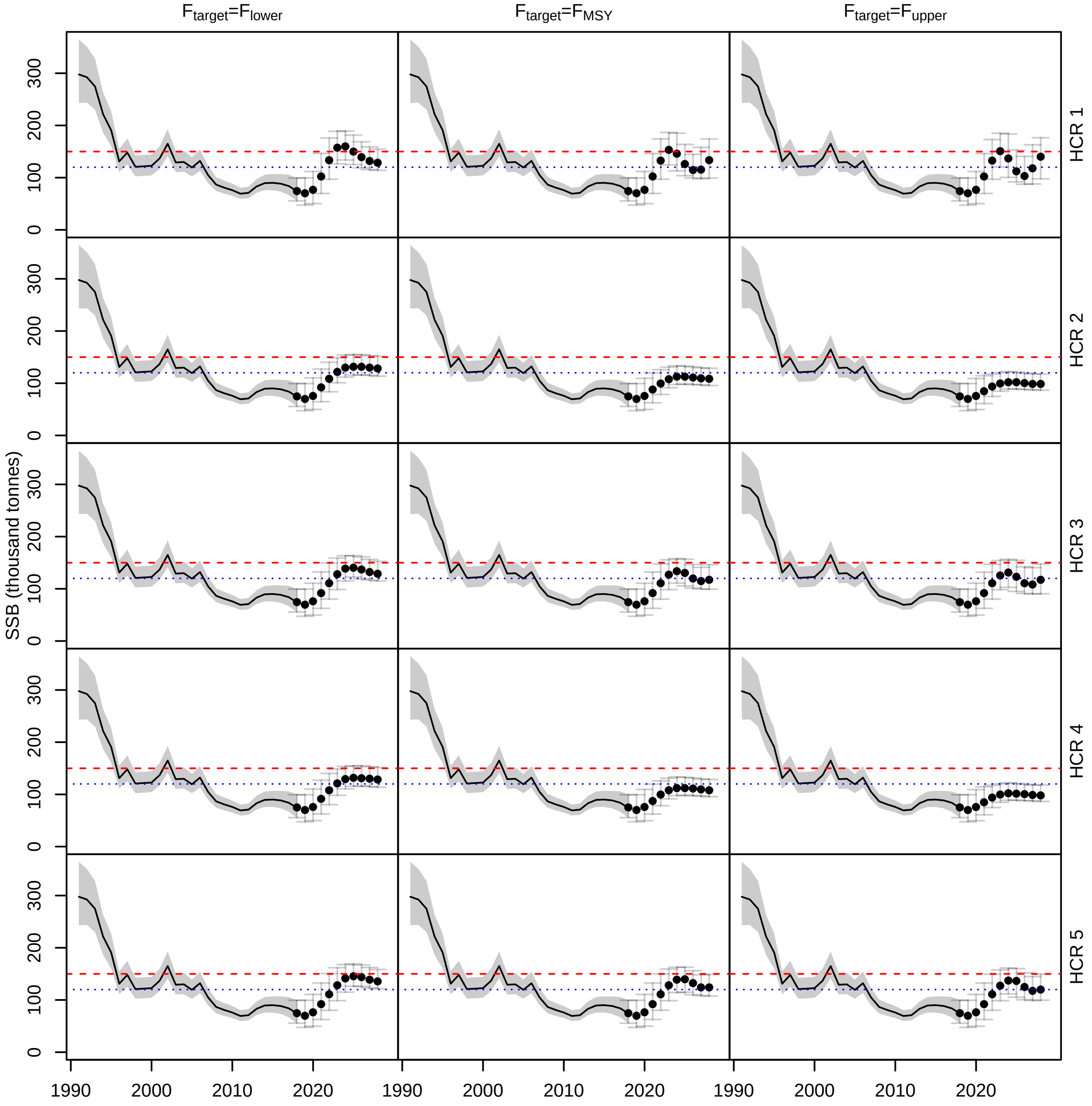


Figure A4.3: WBSS herring spawning stock biomass (SSB) estimated for all HCRs when recruitment is randomly sampled in 2013-2017. The points are the median SSB and the segments the 95% confidence intervals estimated from the 5000 replicates. The red dashed line is $MSY B_{trigger}$ and the blue dotted line is $B_{lim}$.

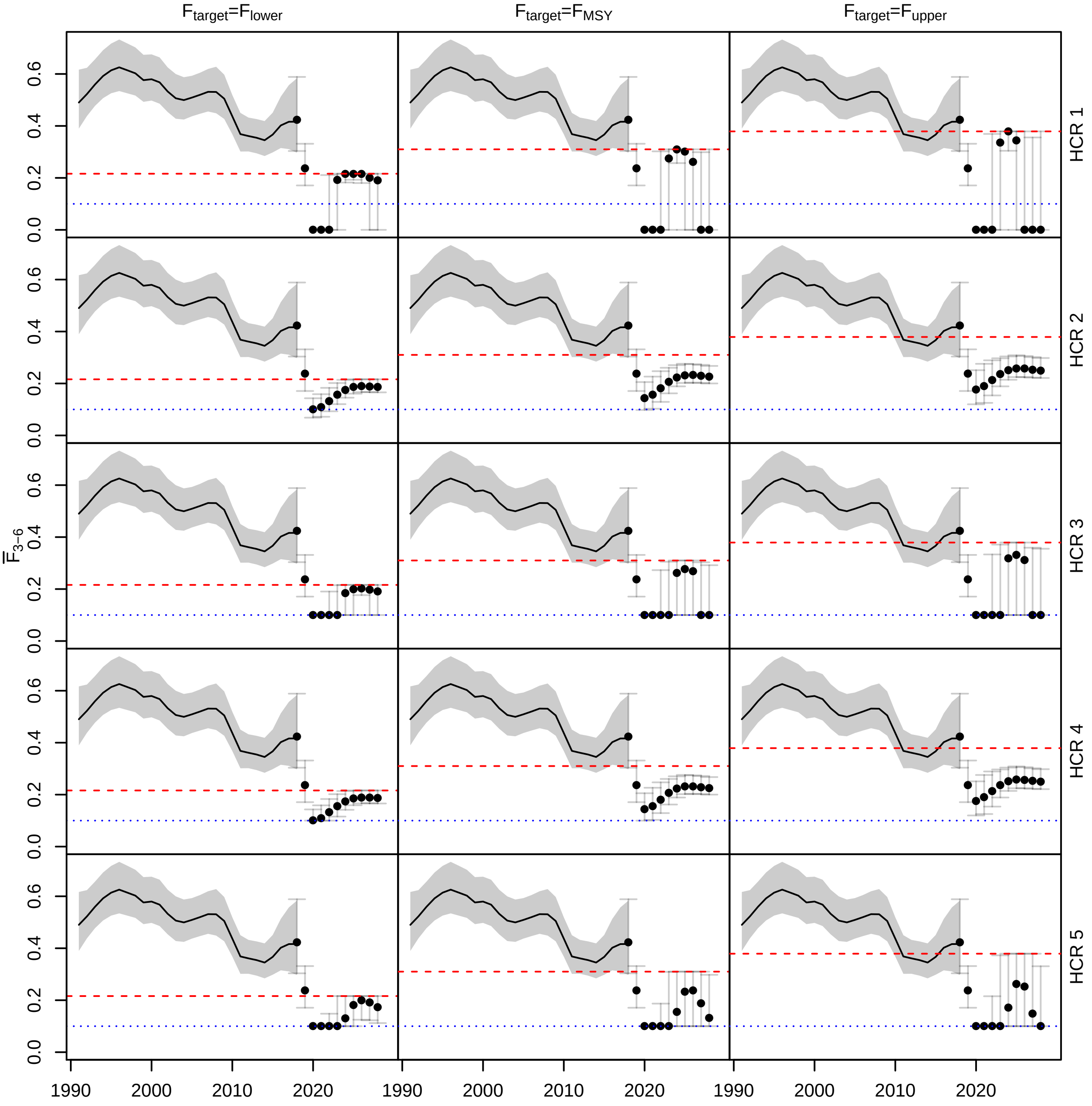


Figure A4.4: Average fishing mortality on WBSS herring for ages 3-6 ($\bar{F}_{3-6}$) estimated for all HCRs when recruitment is randomly sampled in 2013-2017. The points are the median $\bar{F}_{3-6}$ and the segments the 95% confidence intervals estimated from the 5000 replicates. The red dashed line is $F_{target}$ and the blue dotted line is $F = 0.1$.

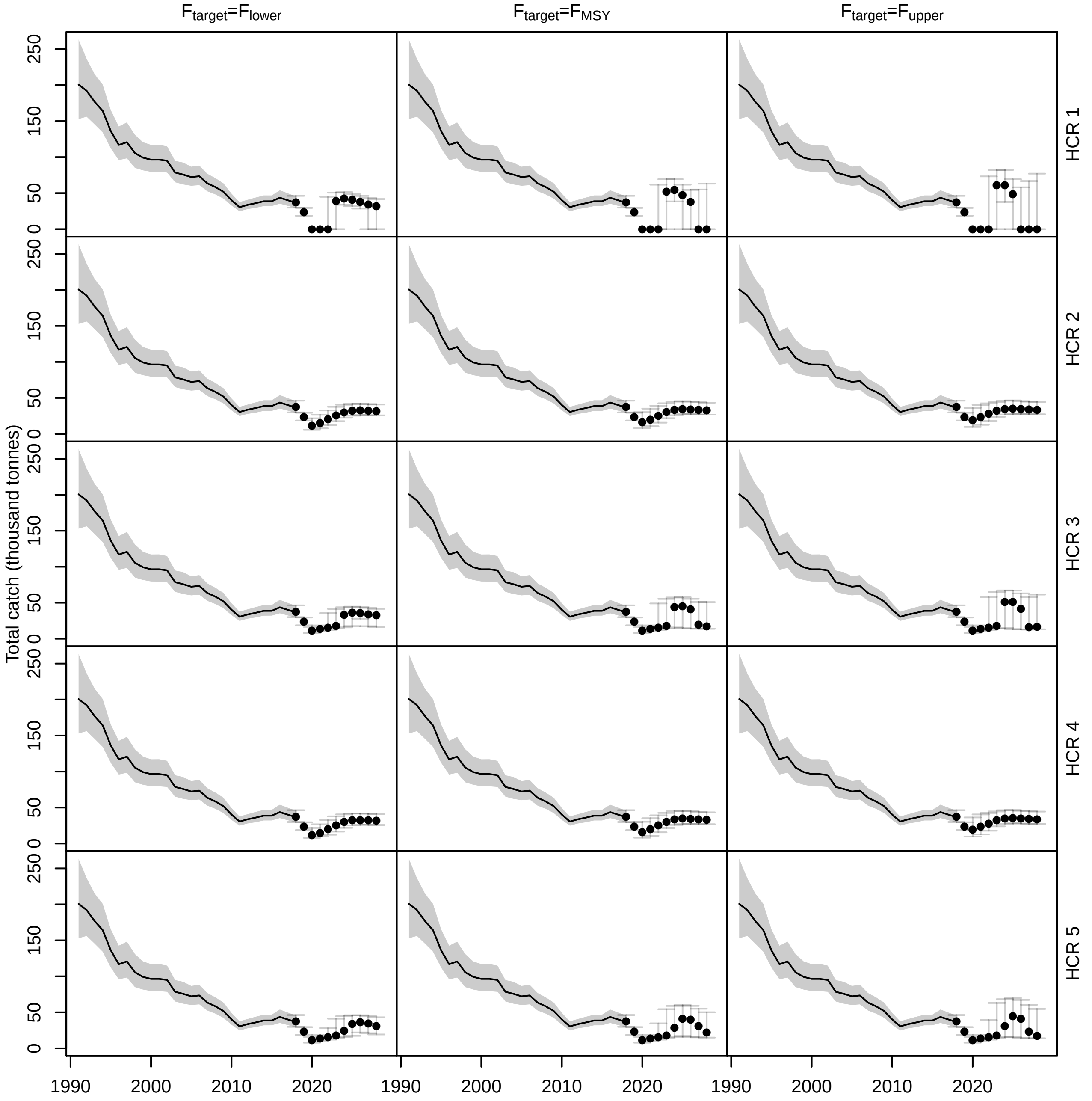


Figure A4.5: Total catch of WBSS herring estimated for all HCRs when recruitment is randomly sampled in 2013-2017. The points are the median catch and the segments the 95% confidence intervals estimated from the 5000 replicates.

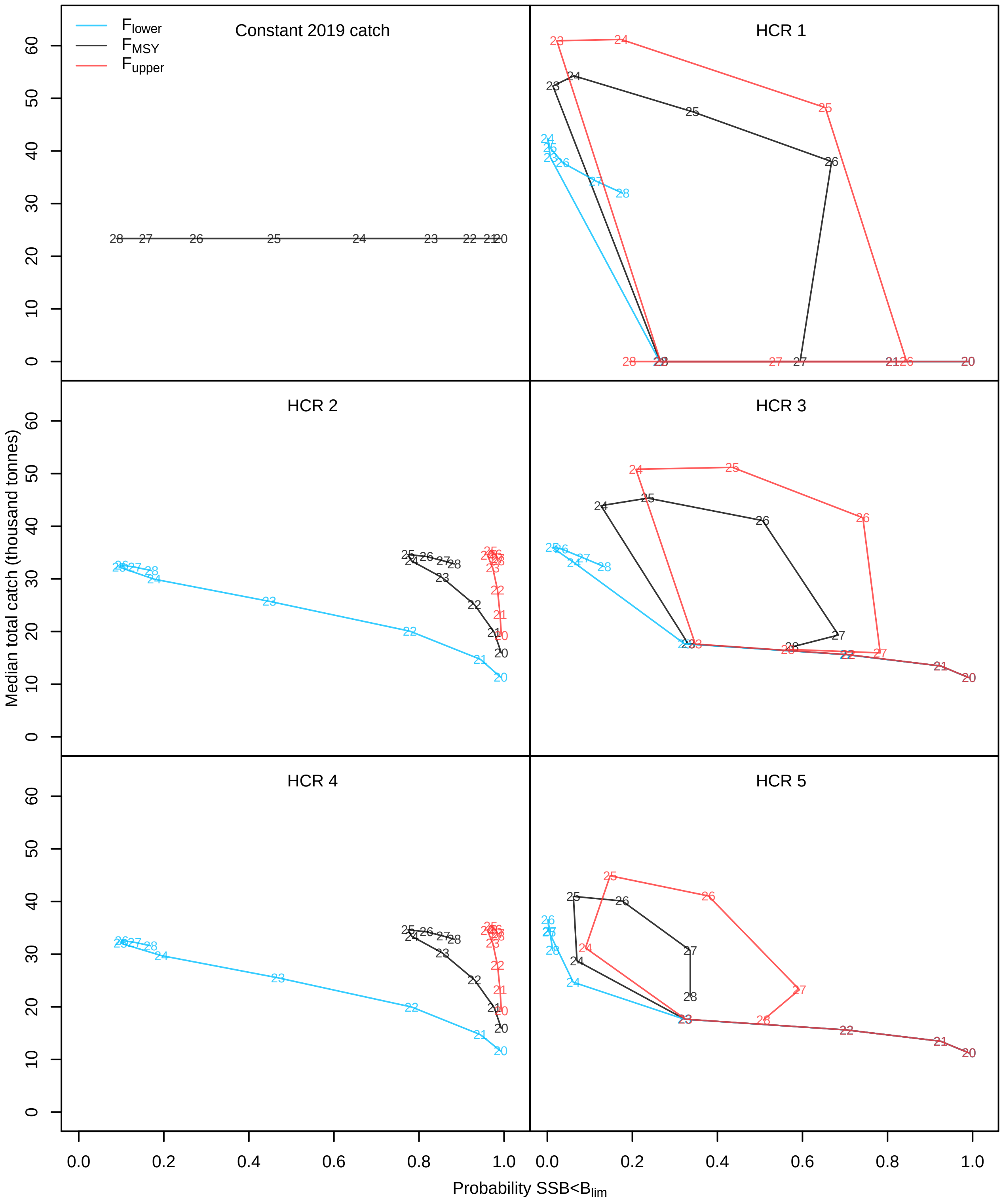


Figure A4.6: WBSS herring median total catch against probability of SSB falling below $B_{lim}$ when recruitment is randomly sampled in 2013-2017. The numbers correspond to the forecast years (2020-2028). Each color represents a $F_{target}$ option. Only one color is showed for the constant 2019 catch base case because the results did not depend on the $F_{target}$.

**North Sea cod** Compared to WBSS herring, cod recovered well for all HCRs and even for all $F_{target}$ values (Figure A4.7). However, this was possible with $\bar{F}_{2-4}$ values staying below any historical $\bar{F}_{2-4}$ level (Figure A4.8) and resulted in a total catch that stayed around levels observed in the past 20 years (Figure A4.9).

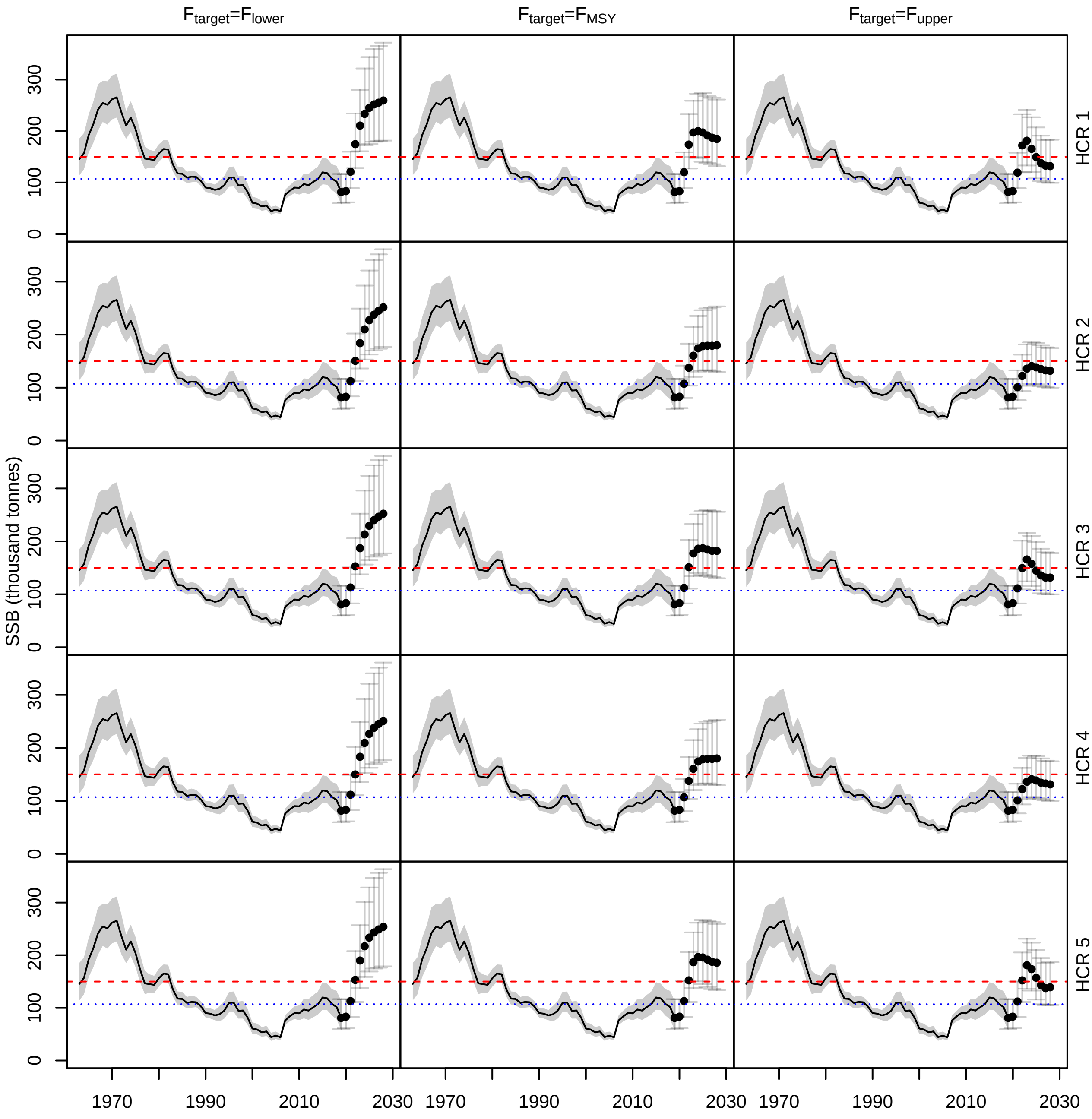


Figure A4.7: North Sea cod spawning stock biomass (SSB) estimated for all HCRs when recruitment is randomly sampled in 1998-2018. The points are the median SSB and the segments the 95% confidence intervals estimated from the 5000 replicates. The red dashed line is $MSYB_{trigger}$ and the blue dotted line is $B_{lim}$.

The risk on North Sea cod decreased rapidly from 2022 for all STs (Figure A4.10). However, despite higher median total catch, having a higher $F_{target}$ increased the risk of SSB falling below $B_{lim}$, notably at the end of the forecast period.

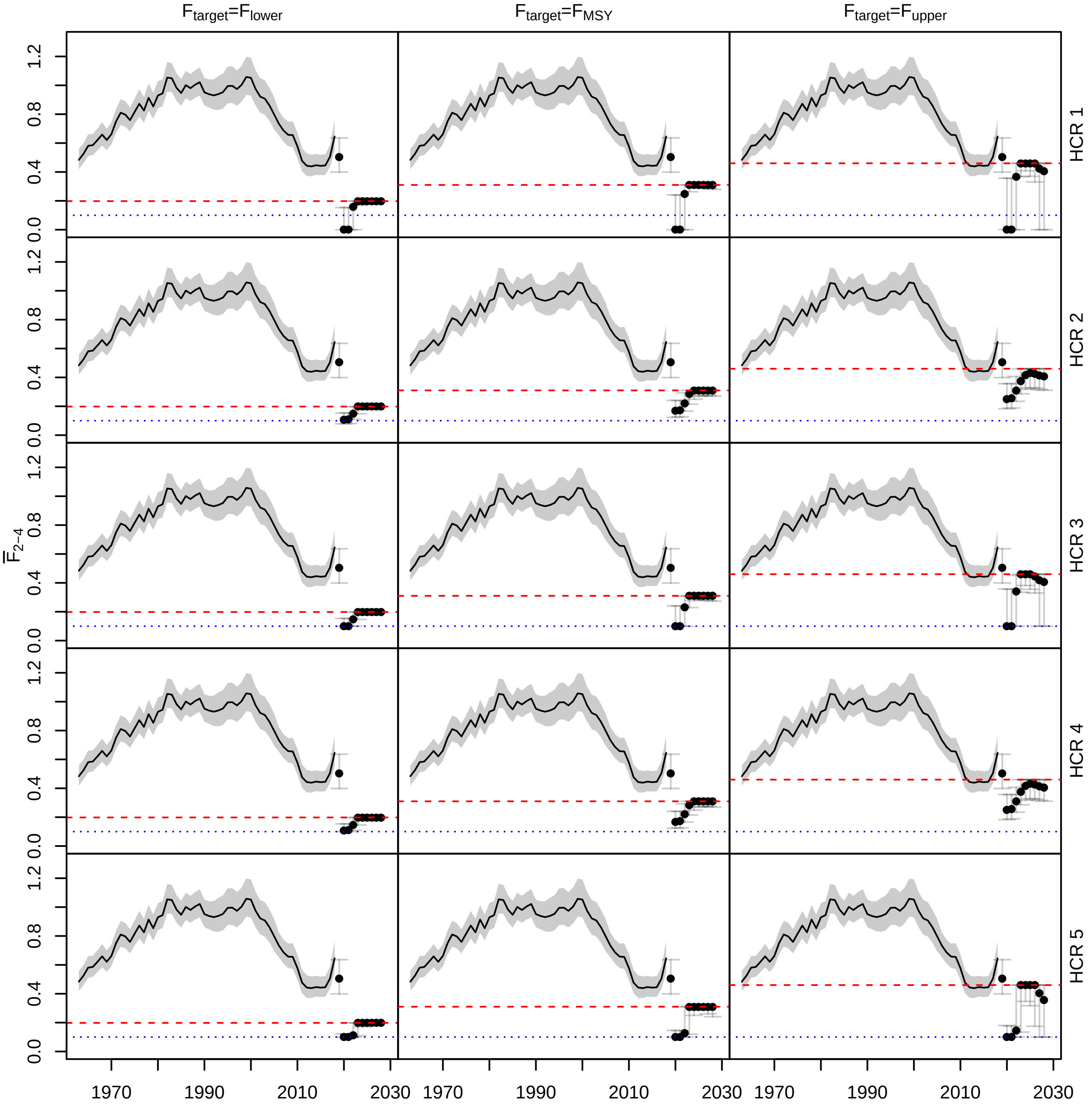


Figure A4.8: Average fishing mortality on North Sea cod for ages 3-6 ($\bar{F}_{2-4}$) estimated for all HCRs when recruitment is randomly sampled in 1998-2018. The points are the median $\bar{F}_{2-4}$ and the segments the 95% confidence intervals estimated from the 5000 replicates. The red dashed line is $F_{target}$ and the blue dotted line is $F = 0.1$.

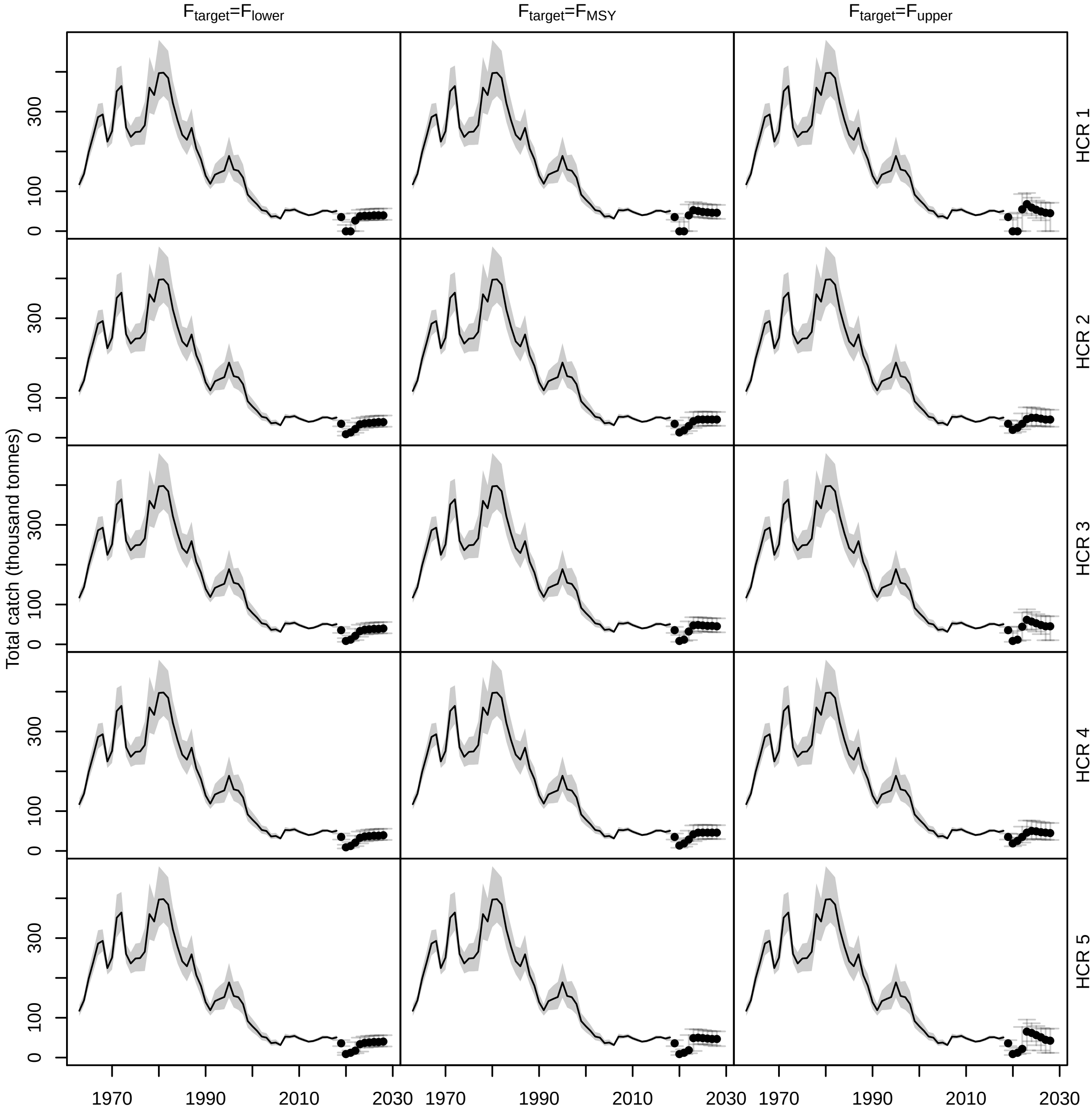


Figure A4.9: Total catch of North Sea cod estimated for all HCRs when recruitment is randomly sampled in 1998-2018. The points are the median catch and the segments the 95% confidence intervals estimated from the 5000 replicates.

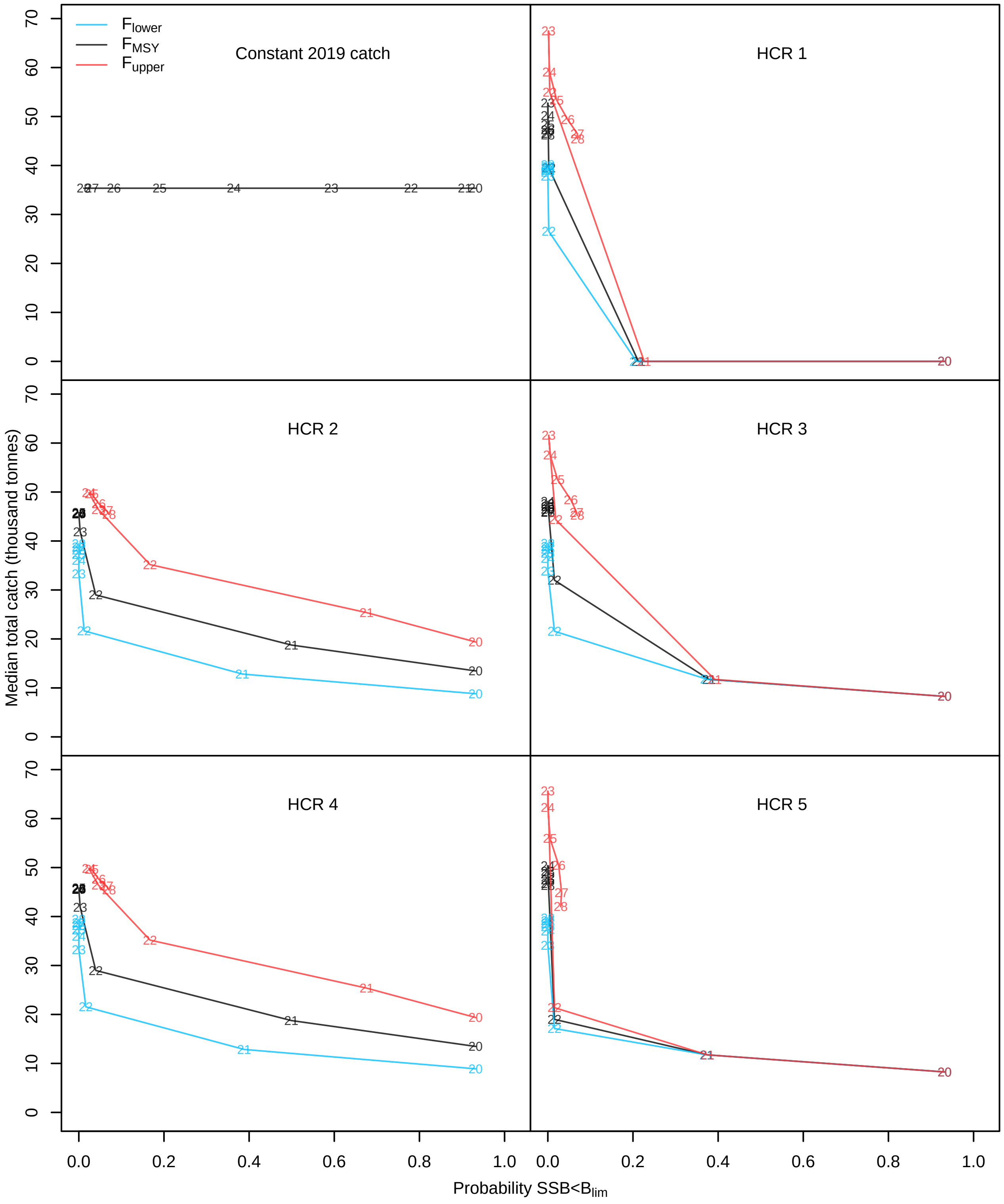


Figure A4.10: North Sea cod median total catch against probability of SSB falling below $B_{lim}$ when recruitment is randomly sampled in 1998-2018. The numbers correspond to the forecast years (2020-2028). Each color represents a $F_{target}$ option. Only one color is showed for the constant 2019 catch base case because the results did not depend on the $F_{target}$.

##### A4.2.2 Recruitment follows a hockey-stick stock-recruitment relationship

**WBSS herring** Assuming a stock-recruitment relationship resulted in an optimistic recovery where herring rebuilt above $B_{lim}$ in the early medium-term ($\sim$ 5 years) and above $MSY B_{trigger}$ by the end of the forecast period for all HCRs (Figure A4.11). Median $\bar{F}_{3-6}$ increased by the end of the forecast period for all HCRs but is reduced to 0 for 3 years for HCR 1 and 0.1 for 4 years for HCRs 3 and 5 (Figure A4.12). These changes in $F$ for these 3 HCRs resulted in marked changes in total catch (Figure A4.13). Despite an optimistic recovery with this recruitment assumption, the risk on the stock is still high even by the end of the forecast period, for most HCRs when $F_{target} \geq F_{MSY}$ (Figure A4.14).

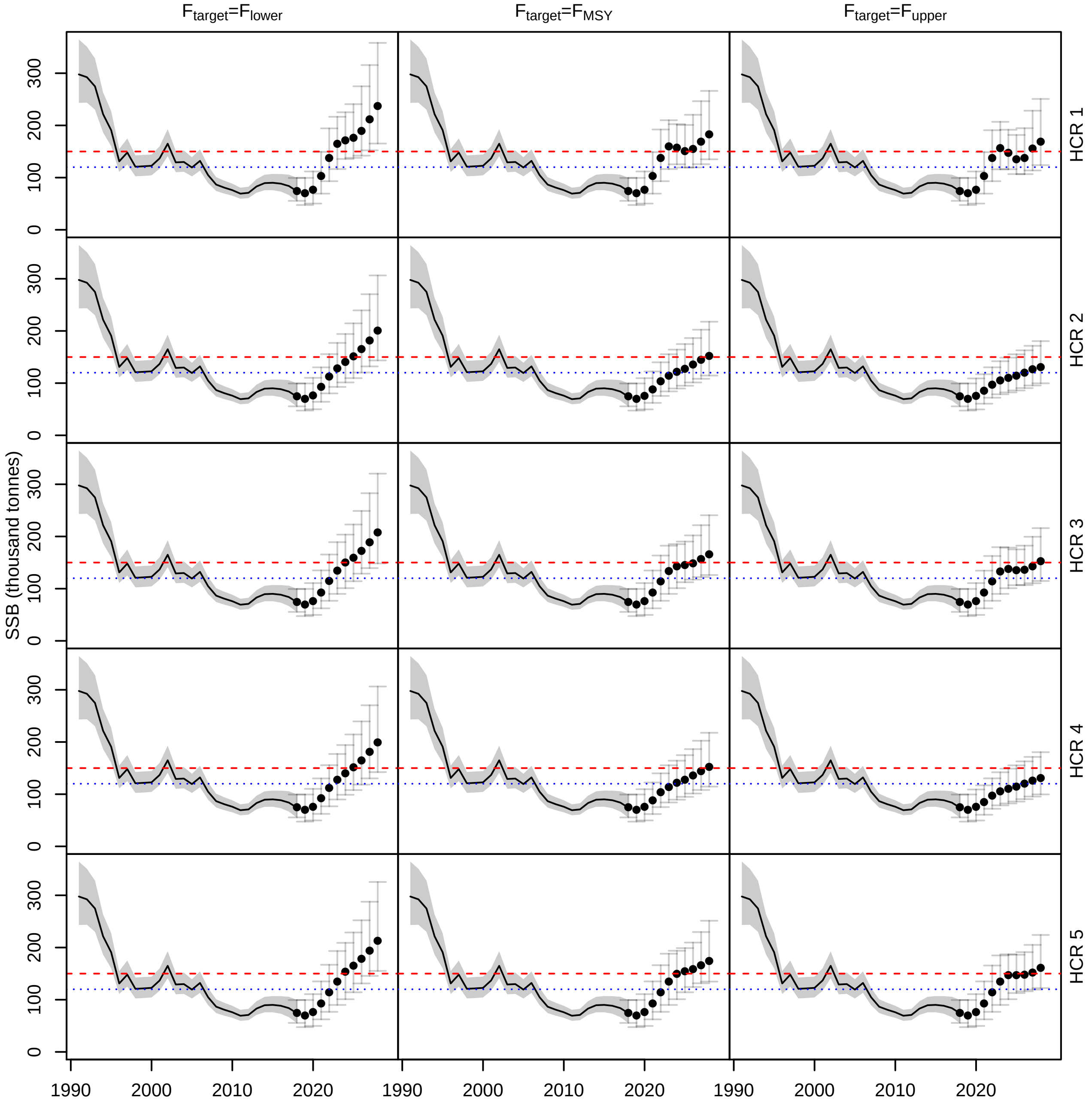


Figure A4.11: WBSS herring spawning stock biomass (SSB) estimated for all HCRs when recruitment follows a hockey-stick stock-recruitment relationship. The points are the median SSB and the segments the 95% confidence intervals estimated from the 5000 replicates. The red dashed line is $MSY B_{trigger}$ and the blue dotted line is $B_{lim}$.

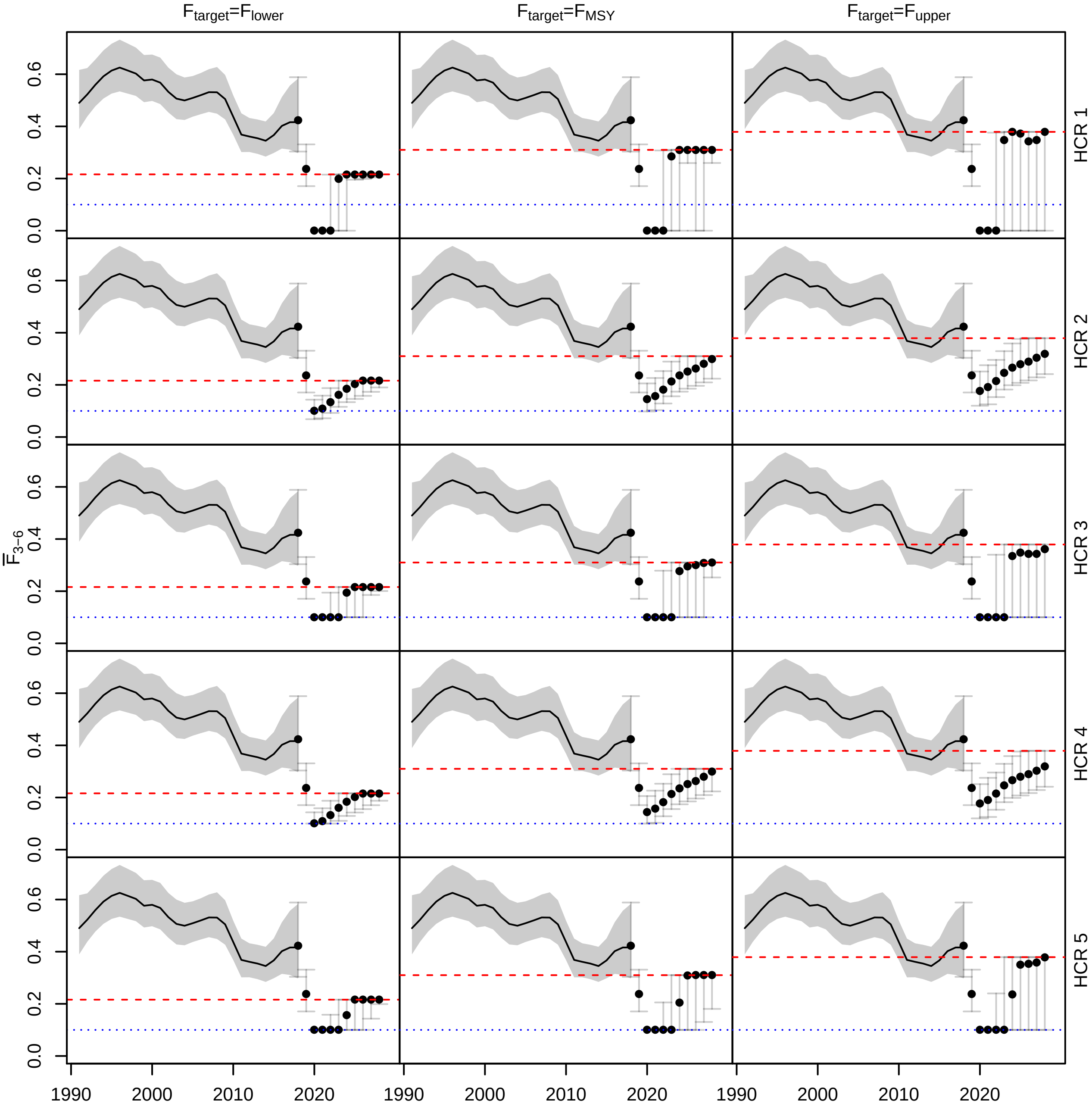


Figure A4.12: Average fishing mortality on WBSS herring for ages 3-6 ($\bar{F}_{3-6}$) estimated for all HCRs when recruitment follows a hockey-stick stock-recruitment relationship. The points are the median $\bar{F}_{3-6}$ and the segments the 95% confidence intervals estimated from the 5000 replicates. The red dashed line is $F_{target}$ and the blue dotted line is $F = 0.1$.

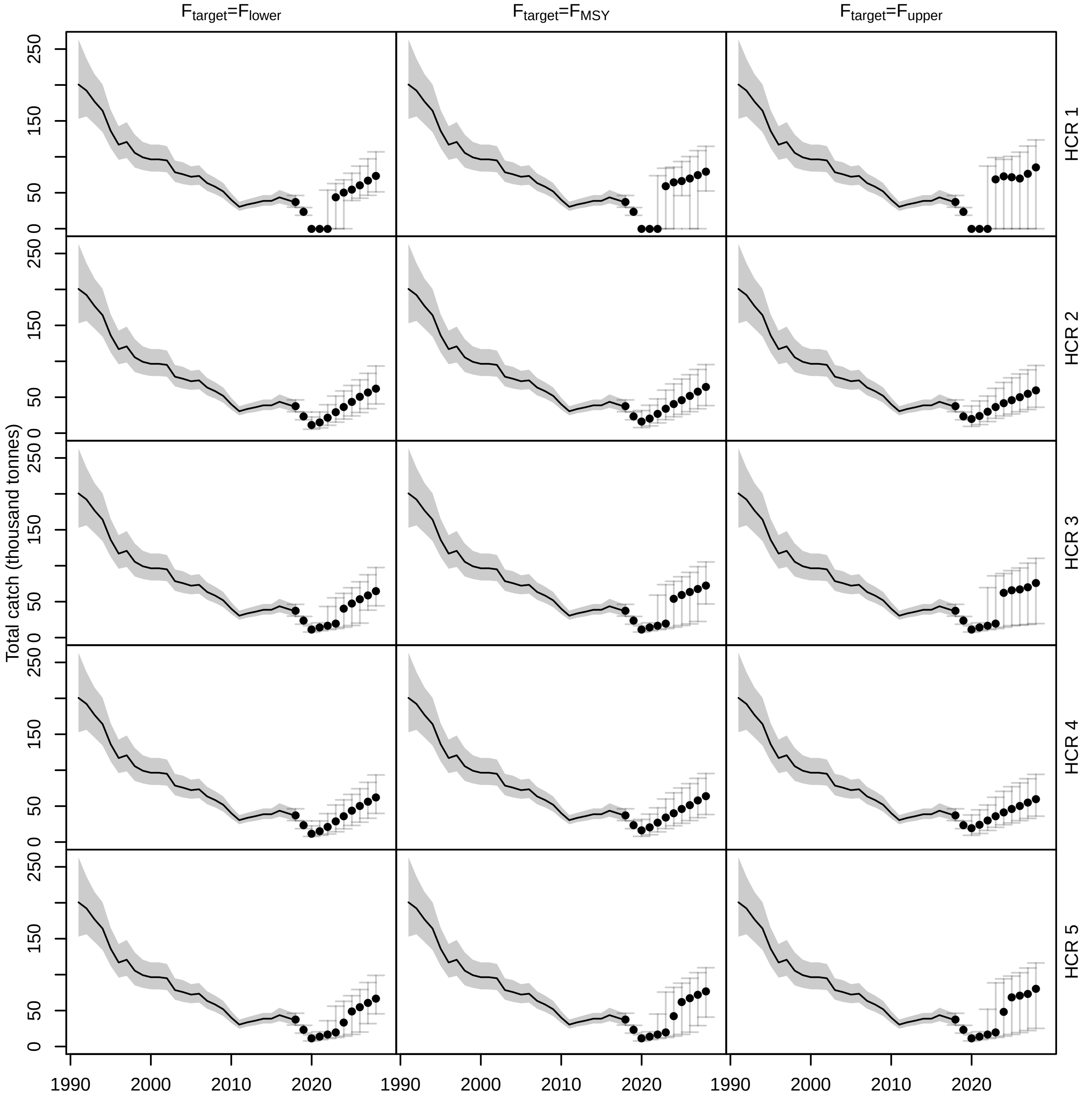


Figure A4.13: Total catch of WBSS herring estimated for all HCRs when recruitment follows a hockey-stick stock-recruitment relationship. The points are the median catch and the segments the 95% confidence intervals estimated from the 5000 replicates.

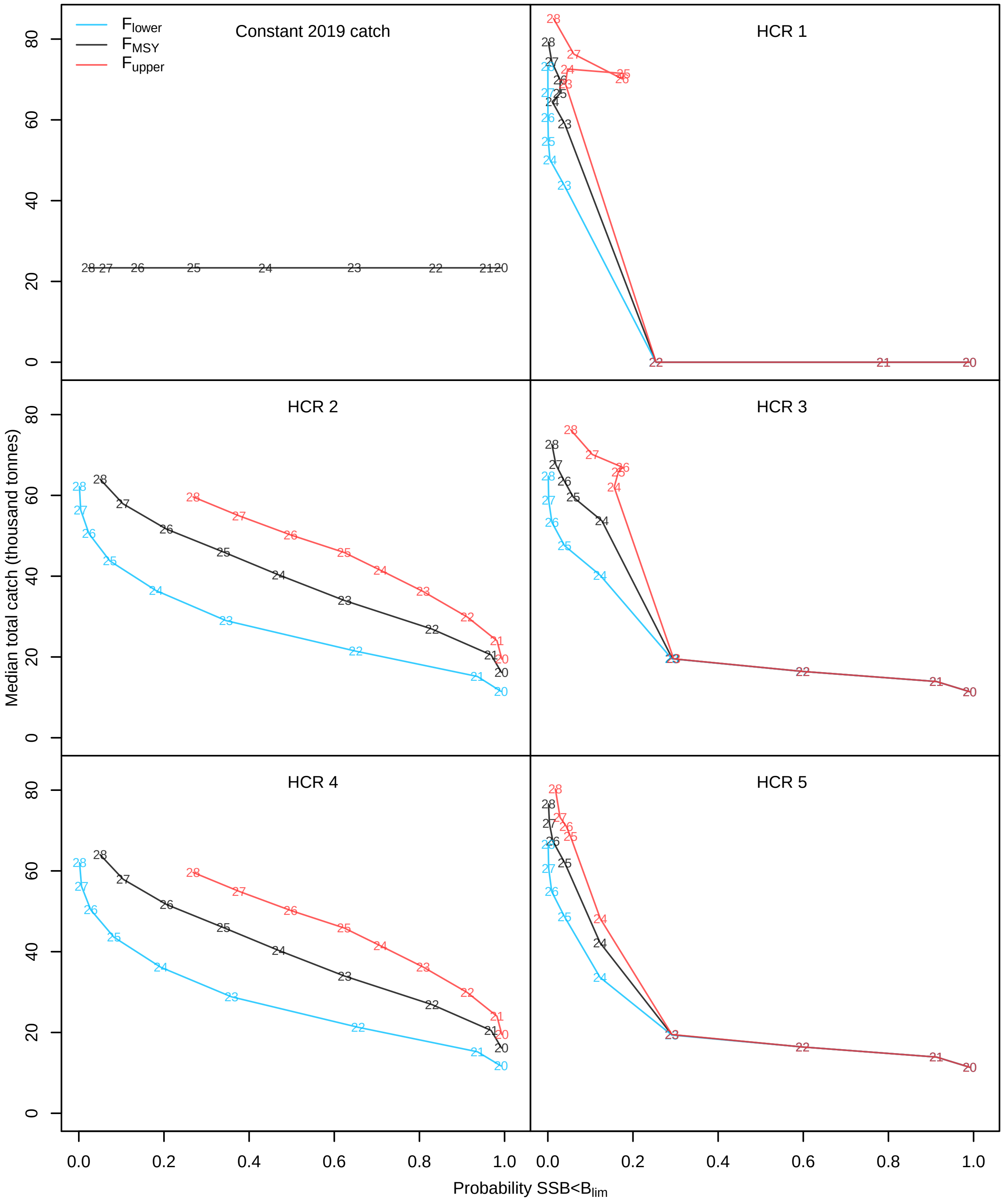


Figure A4.14: WBSS herring median total catch against probability of SSB falling below $B_{lim}$ when recruitment follows a hockey-stick stock-recruitment relationship. The numbers correspond to the forecast years (2020-2028). Each color represents a $F_{target}$ option. Only one color is showed for the constant 2019 catch base case because the results did not depend on the $F_{target}$.

**North Sea cod** North Sea cod SSB increased rapidly above historical levels when recruitment follows a hockey-stick relationship for all HCRs (Figure A4.15). Recovery was improved for low value of $F_{target}$. Median $\bar{F}_{2-4}$ got to its maximum in few years after the start of the forecast for all HCRs and $F_{target}$ levels (Figure A4.16). Given that all $F_{target}$ levels are below any historical $F$, catch increased compared to the end of the assessment estimates but stayed around levels observed in the past 30 years (Figure A4.17). The risk on the cod stock is rapidly reduced from 2022 and until the end of the forecast period for all STs and $F_{target}$ levels (Figure A4.18).

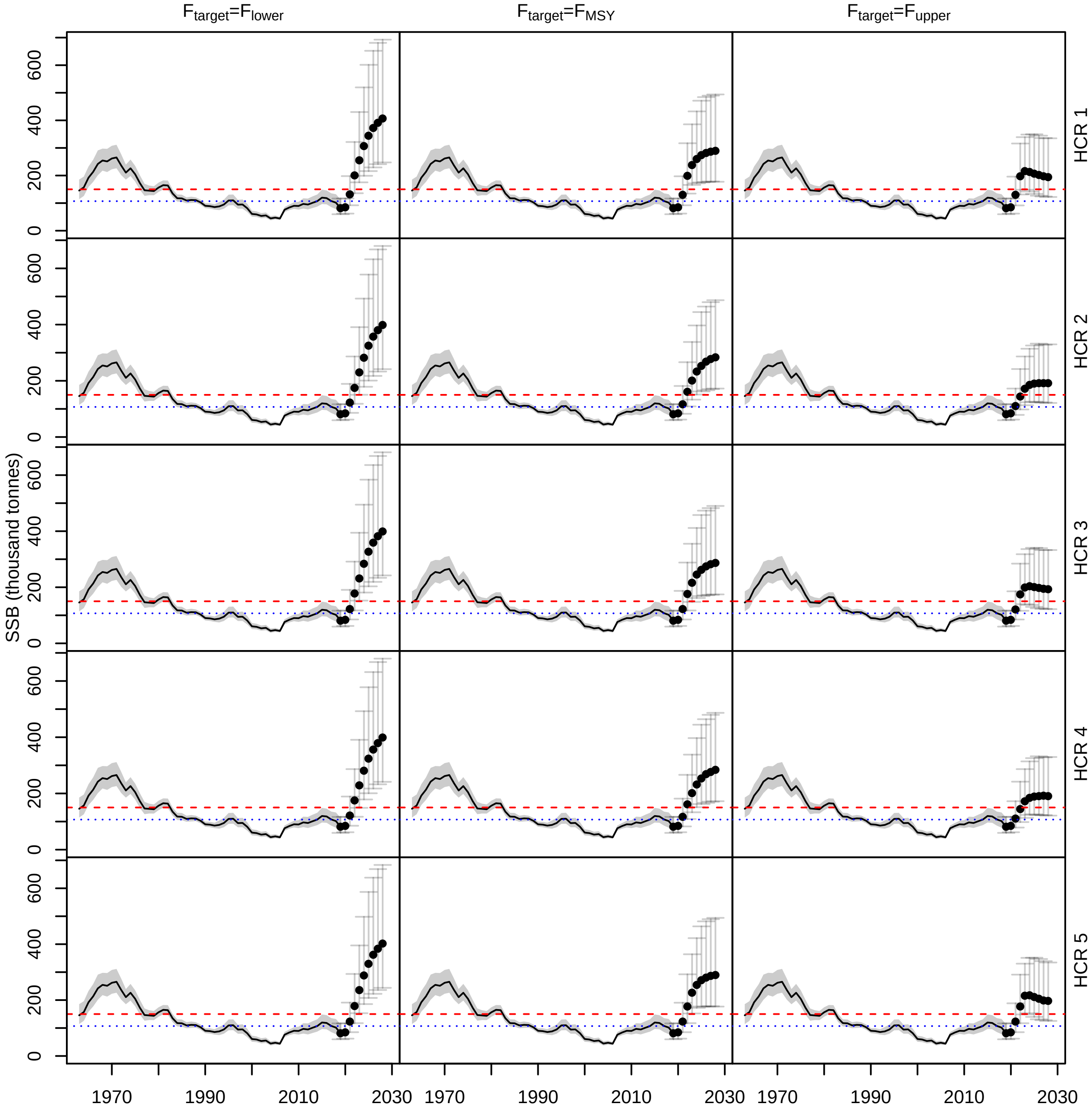


Figure A4.15: North Sea cod spawning stock biomass (SSB) estimated for all HCRs when recruitment follows a hockey-stick stock-recruitment relationship. The points are the median SSB and the segments the 95% confidence intervals estimated from the 5000 replicates. The red dashed line is $MSY B_{trigger}$ and the blue dotted line is $B_{lim}$.

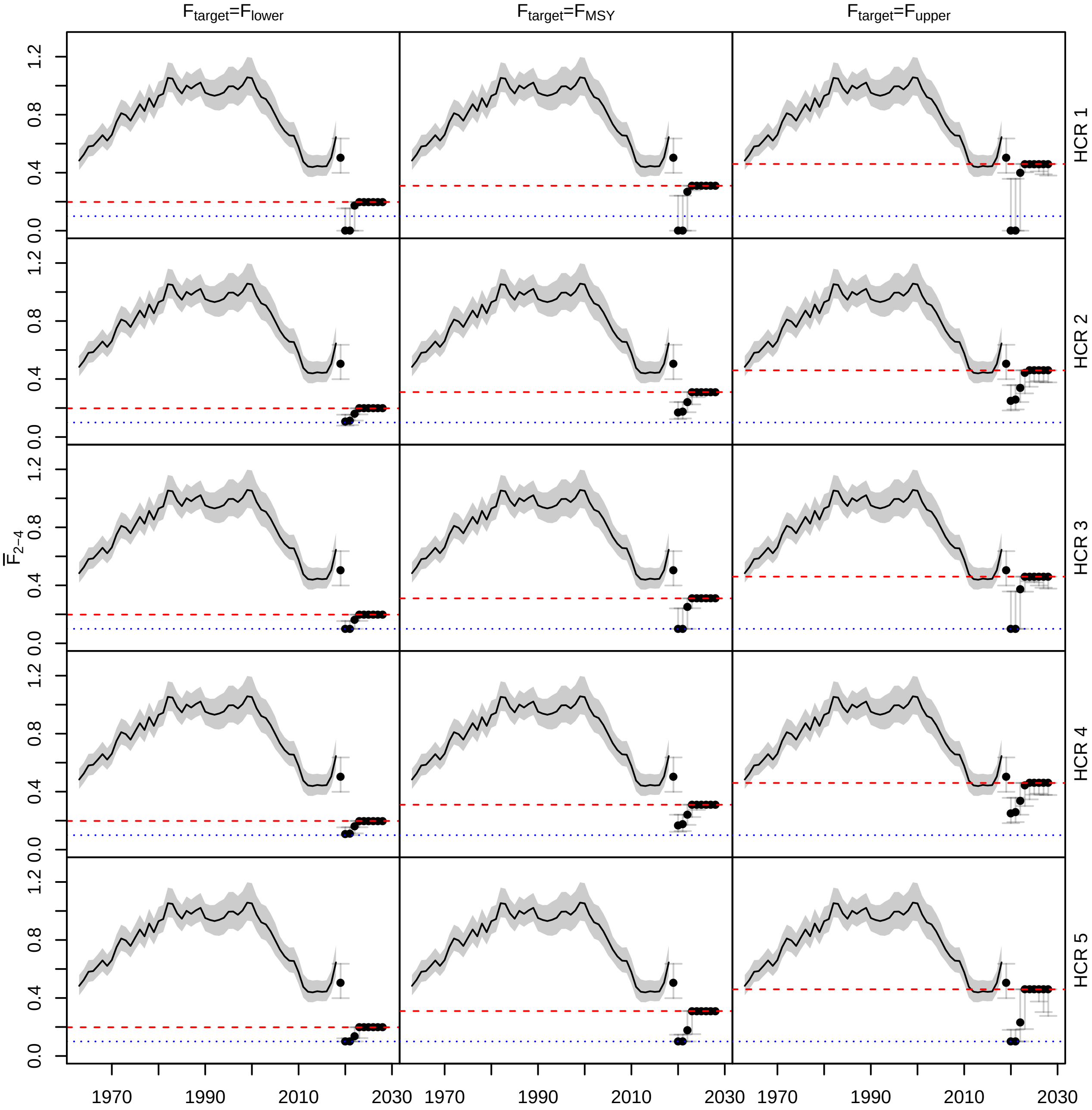


Figure A4.16: Average fishing mortality on North Sea cod for ages 2-4 ($\bar{F}_{2-4}$) estimated for all HCRs when recruitment follows a hockey-stick stock-recruitment relationship. The points are the median $\bar{F}_{2-4}$ and the segments the 95% confidence intervals estimated from the 5000 replicates. The red dashed line is $F_{target}$ and the blue dotted line is $F = 0.1$.

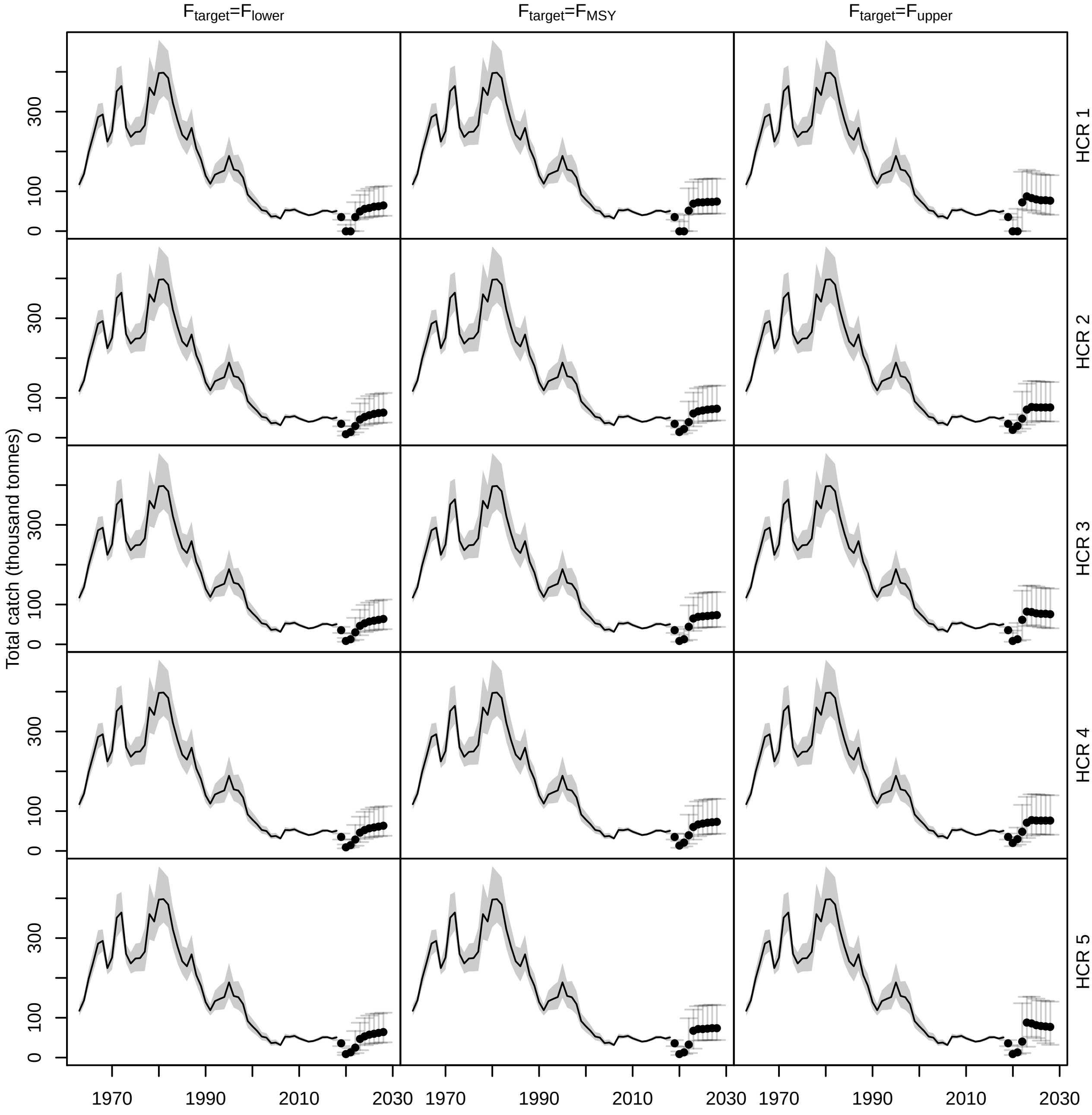


Figure A4.17: Total catch of North Sea cod estimated for all HCRs when recruitment follows a hockey-stick stock-recruitment relationship. The points are the median catch and the segments the 95% confidence intervals estimated from the 5000 replicates.

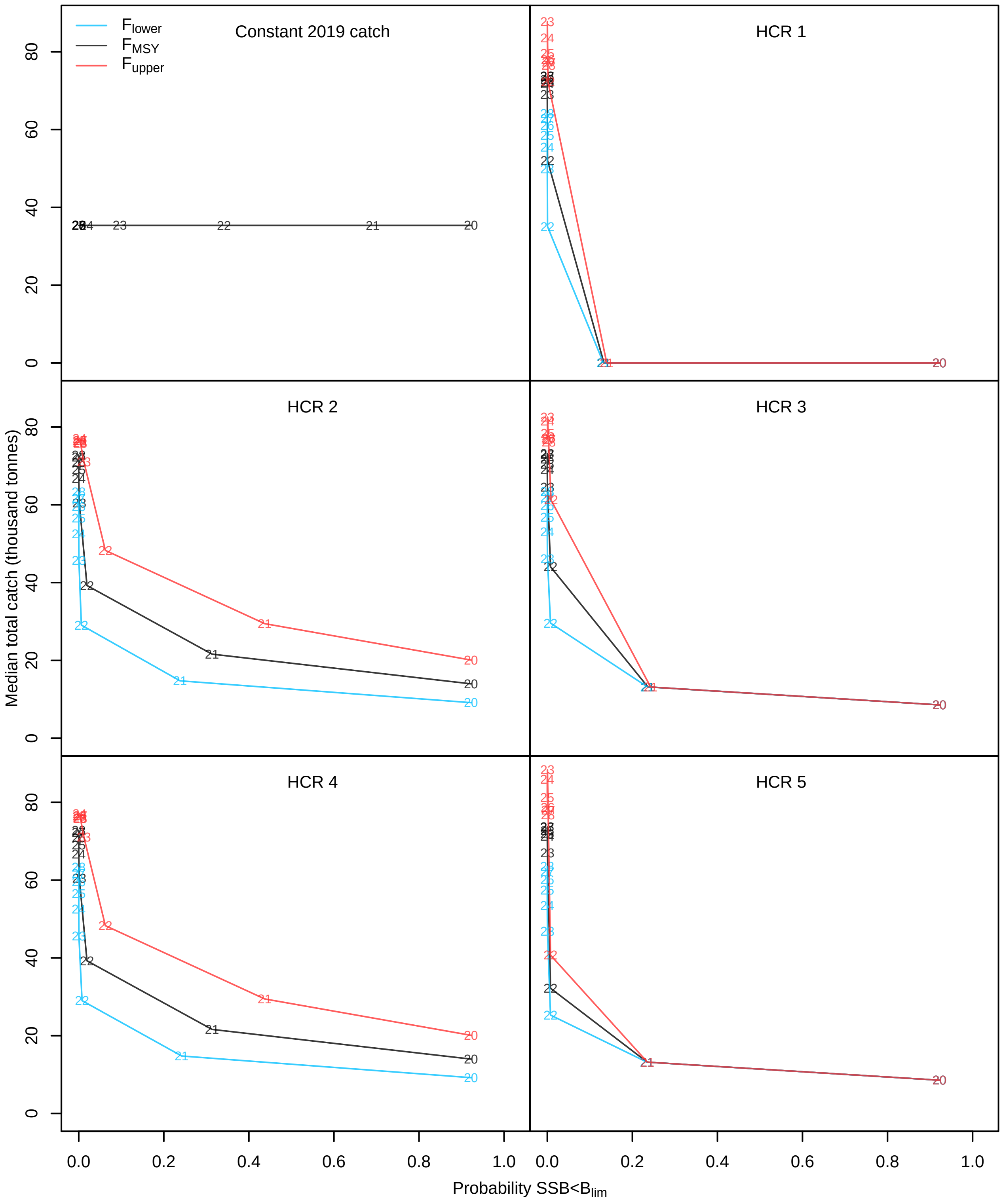


Figure A4.18: North Sea cod median total catch against probability of SSB falling below $B_{lim}$ when recruitment follows a hockey-stick stock-recruitment relationship. The numbers correspond to the forecast years (2020-2028). Each color represents a $F_{target}$ option. Only one color is showed for the constant 2019 catch base case because the results did not depend on the $F_{target}$.

##### A4.2.3 Recruitment as a random walk with negative drift

**WBSS herring** Assuming recruitment followed a random walk with negative drift worsened stock perception since none of the HCRs enabled median recovery of the stock above $B_{lim}$ for consecutive years until 2028 (Figure A4.19). As expected, uncertainty around SSB increased over time, however, similarly to what was observed in Figure A4.3, all options predicted an overall increase of SSB compared to the 2018 estimate. None of the options resulted in a median SSB above $MSYB_{trigger}$, but the upper limit of the 95% confidence interval around SSB went above $MSYB_{trigger}$, after only few years of management, for all HCR and $F_{target}$ options.

$\bar{F}_{3-6}$ had a larger probability to stay at its lowest level for HCRs 3 and 5 when a random walk with negative drift was assumed in the forecasts (Figure A4.20) while it kept varying for HCR 1. Median total catches were overall lower in the random walk scenario compared to the two others recruitment options but the 95% confidence intervals were wider (Figure A4.21).

Assuming a random walk with negative drift on recruitment in the forecasts showed very small probability of rebuilding for all HCRs except HCR 1, with probabilities of SSB falling below $B_{lim}$ above 50% no matter the $F_{target}$ value (Figure A4.22). HCRs 3 and 5, and HCRs 2 and 4 for $F_{target} = F_{lower}$ showed higher probability of rebuilding than the constant 2019 catch option but resulted in lower total catches. HCR 1 presented the same spiral pattern as in Figure A4.6, where after few years with zero catch, catches and probability of rebuilding increased but then decreased again by the end of the forecast period. Catches were lower compared to the results with recruitment randomly sampled in 2013-2017 (Figure A4.6).

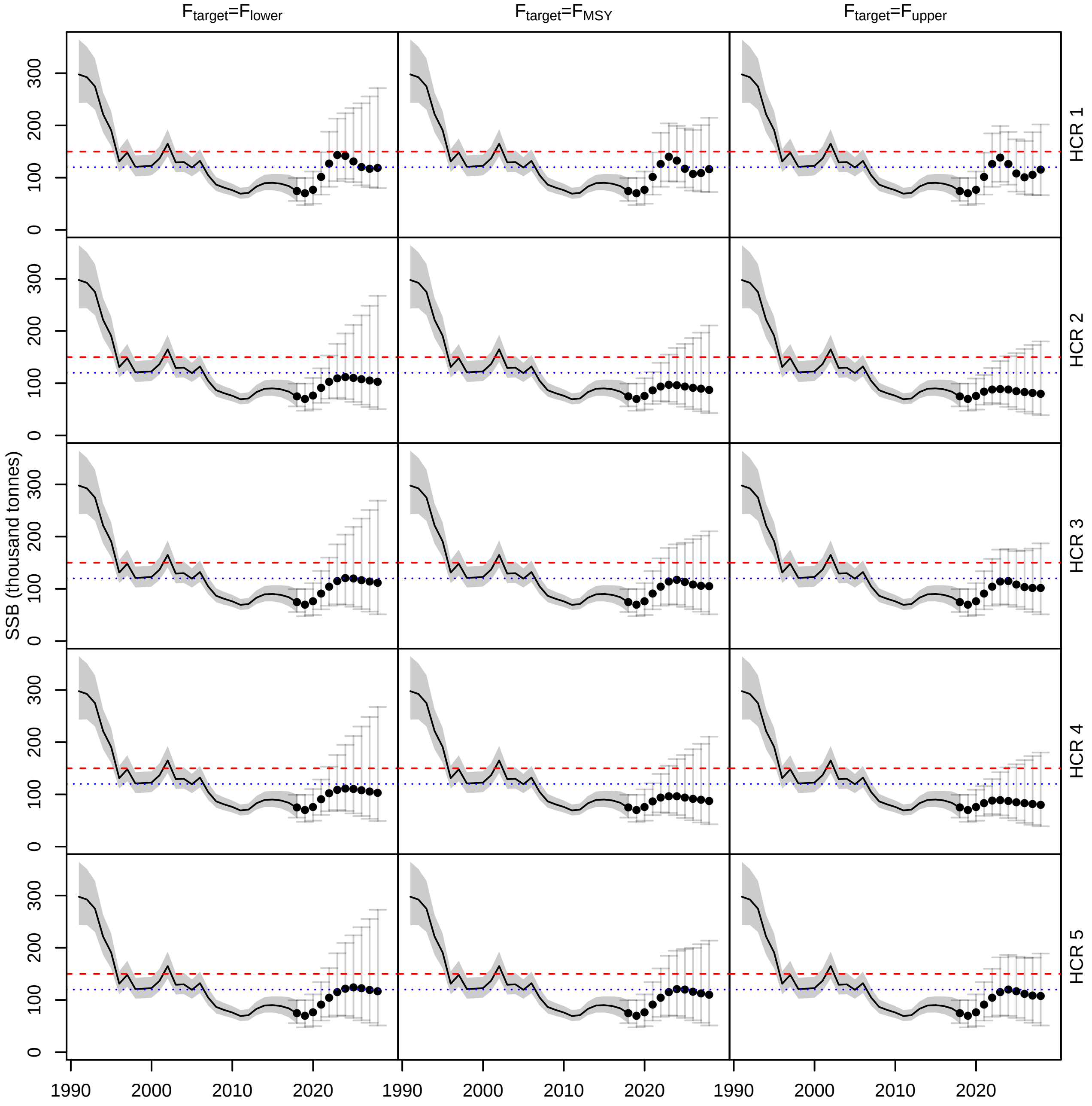


Figure A4.19: WBSS herring spawning stock biomass (SSB) estimated for all HCRs when recruitment is a random walk with negative drift. The points are the median SSB and the segments the 95% confidence intervals estimated from the 5000 replicates. The red dashed line is $MSYB_{trigger}$ and the blue dotted line is $B_{lim}$.

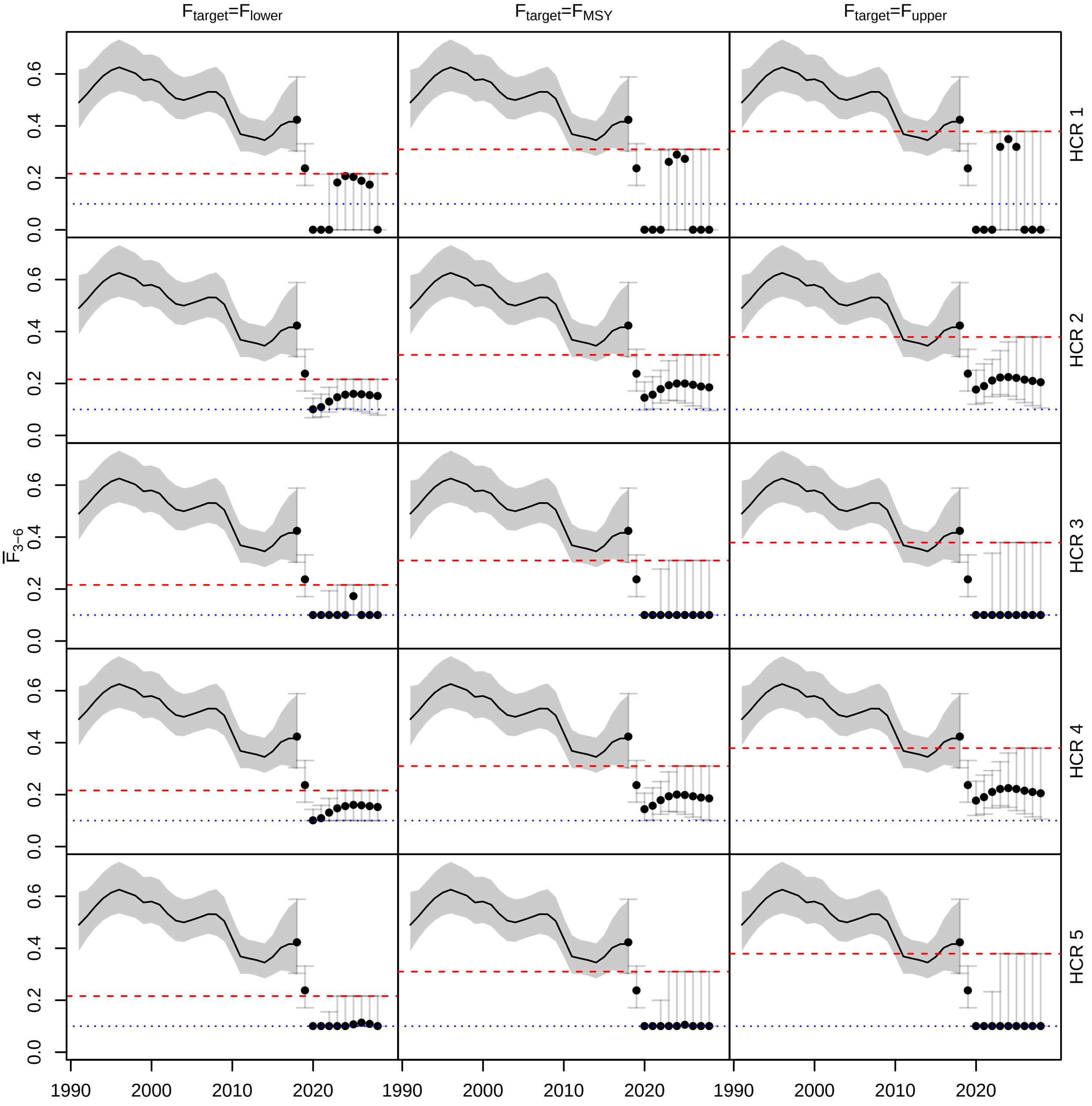


Figure A4.20: Average fishing mortality on WBSS herring for ages 3-6 ($\bar{F}_{3-6}$) estimated for all HCRs when recruitment is a random walk with negative drift. The points are the median $\bar{F}_{3-6}$ and the segments the 95% confidence intervals estimated from the 5000 replicates. The red dashed line is $F_{target}$ and the blue dotted line is $F = 0.1$.

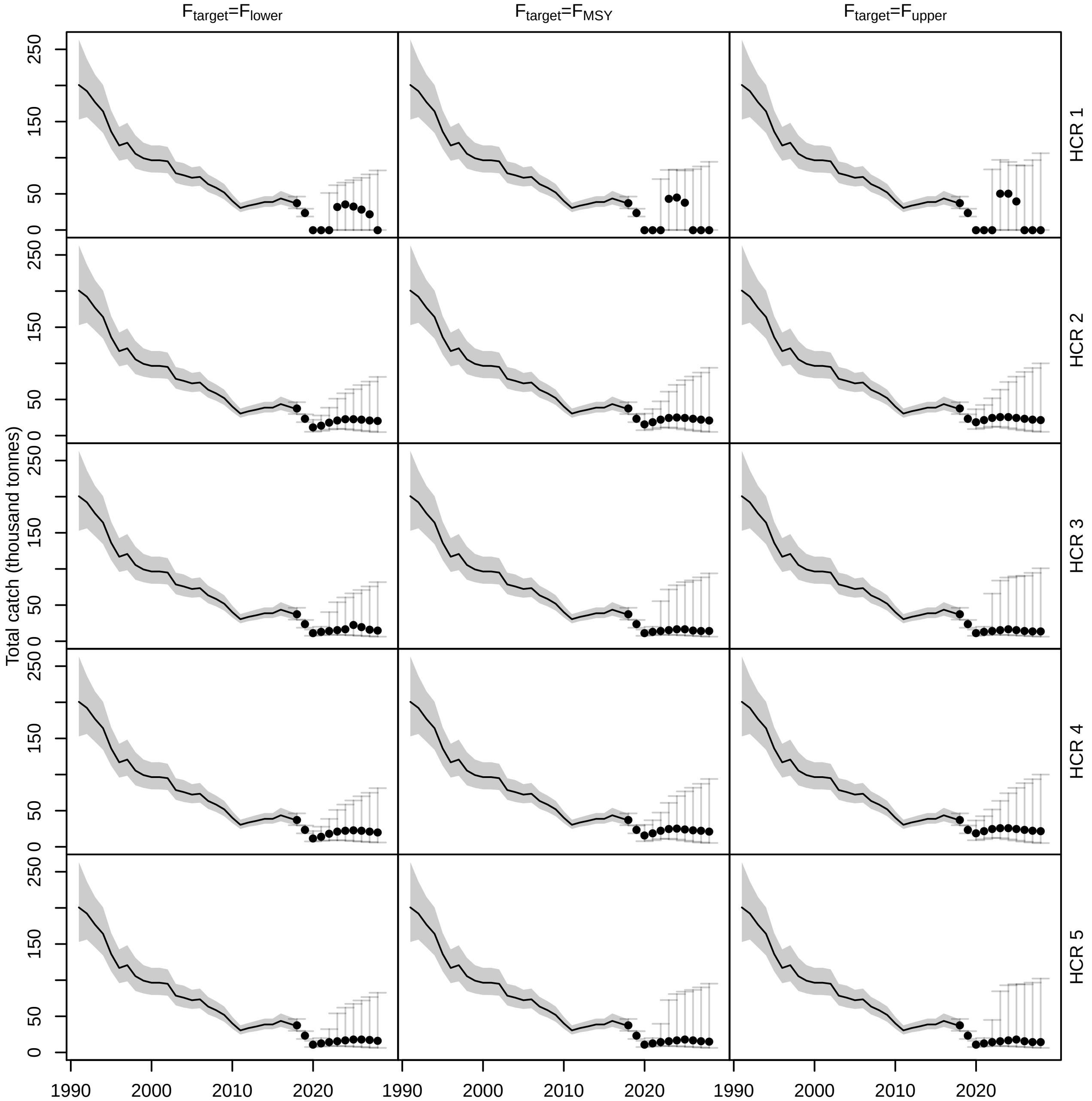


Figure A4.21: Total catch of WBSS herring estimated for all HCRs when recruitment is a random walk with negative drift. The points are the median catch and the segments the 95% confidence intervals estimated from the 5000 replicates.

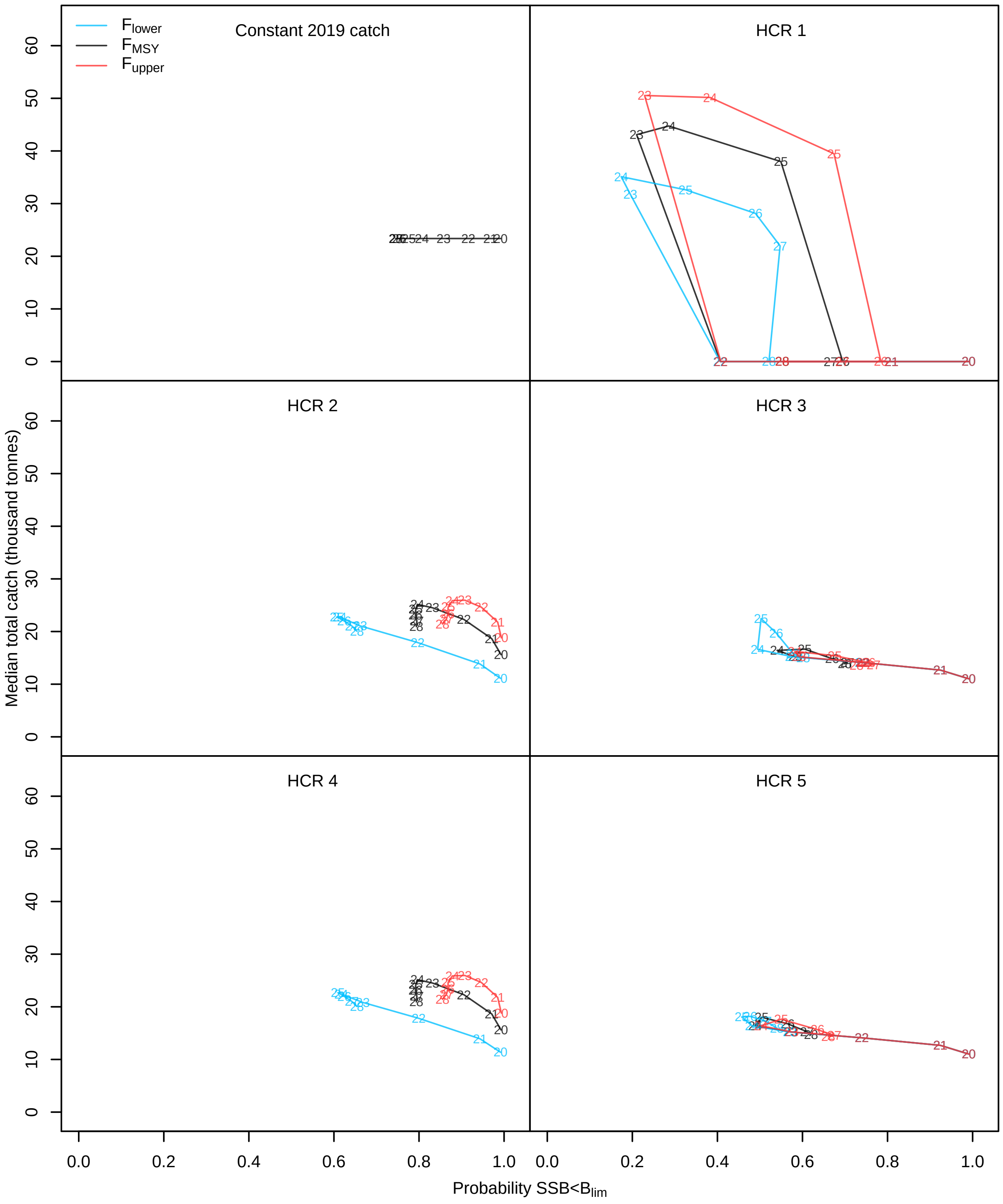


Figure A4.22: WBSS herring median total catch against probability of SSB falling below $B_{lim}$ when recruitment is a random walk with negative drift. The numbers correspond to the forecast years (2020-2028). Each color represents a $F_{target}$ option. Only one color is shown for the constant 2019 catch base case because the results did not depend on the $F_{target}$.

**North Sea cod** Overall, assuming a random walk with negative drift in recruitment still resulted in a recovery above $B_{lim}$ for cod (Figure A4.7) with median SSB being above $MSYB_{trigger}$ for all HCRs when $F_{target} \leq F_{MSY}$ (Figure A4.23).

Compared to the other two recruitment assumptions, the random walk with negative drift in recruitment increased the probability of getting the lower limit of the confidence interval around $\bar{F}_{2-4}$ at low values (Figure A4.24). This was also visible for total catch but, due to variance of the random walk increasing overtime, the upper limit of the confidence interval went above historical catch levels by the end of the forecast period for all HCRs (Figure A4.25).

Risk on the cod stock is clearly higher when recruitment is assumed to decrease in the future (Figure A4.26) compared to the other two recruitment assumptions (Figures A4.10 and A4.18). For all $F_{target}$ levels and STs, risk to fall below $B_{lim}$ was above 25% by the end of the forecast period.

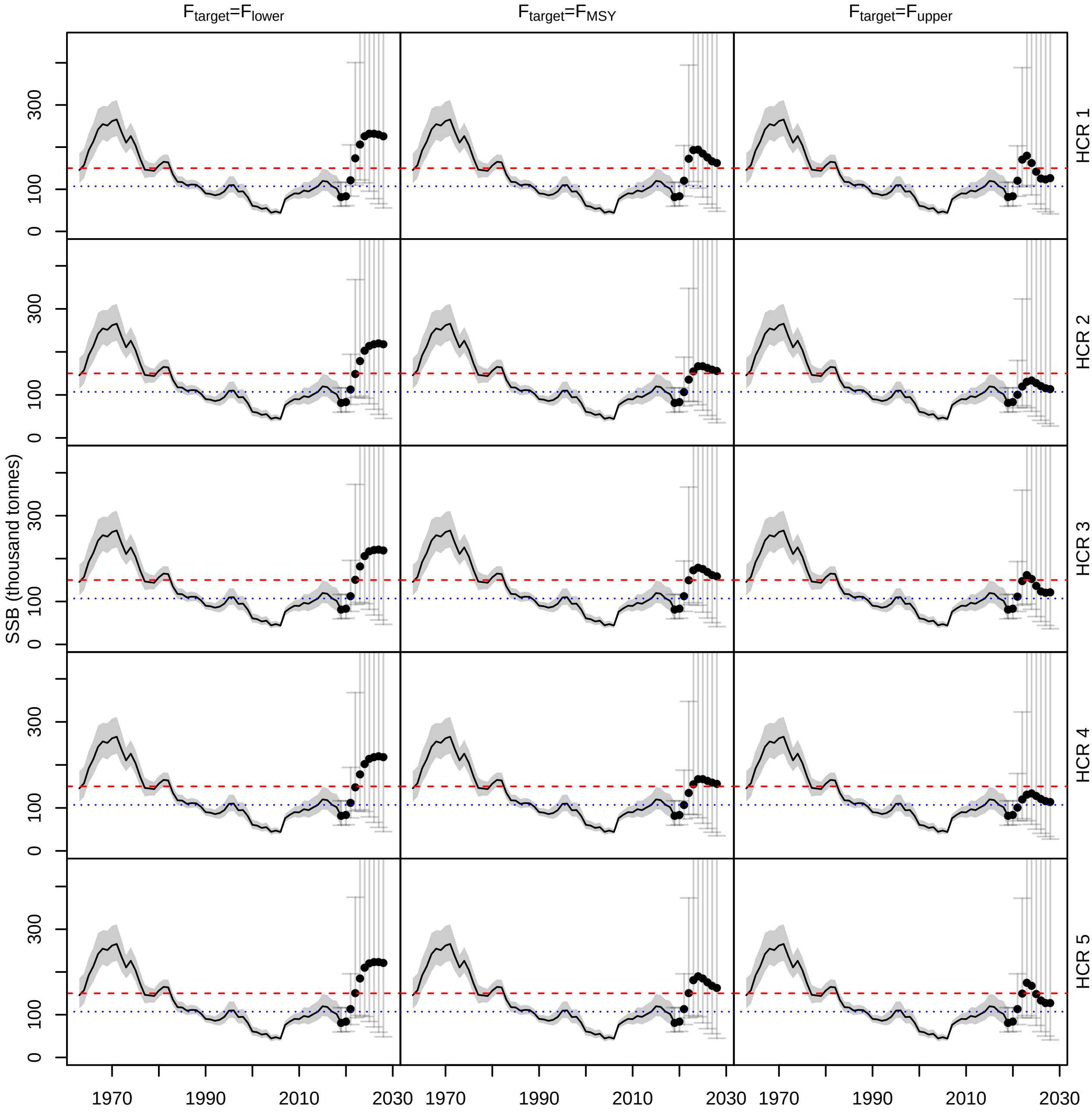


Figure A4.23: North Sea cod spawning stock biomass (SSB) estimated for all HCRs when recruitment is a random walk with negative drift. The points are the median SSB and the segments the 95% confidence intervals estimated from the 5000 replicates. The red dashed line is $MSYB_{trigger}$ and the blue dotted line is $B_{lim}$. For plotting convenience the upper limits of the confidence intervals are not shown.

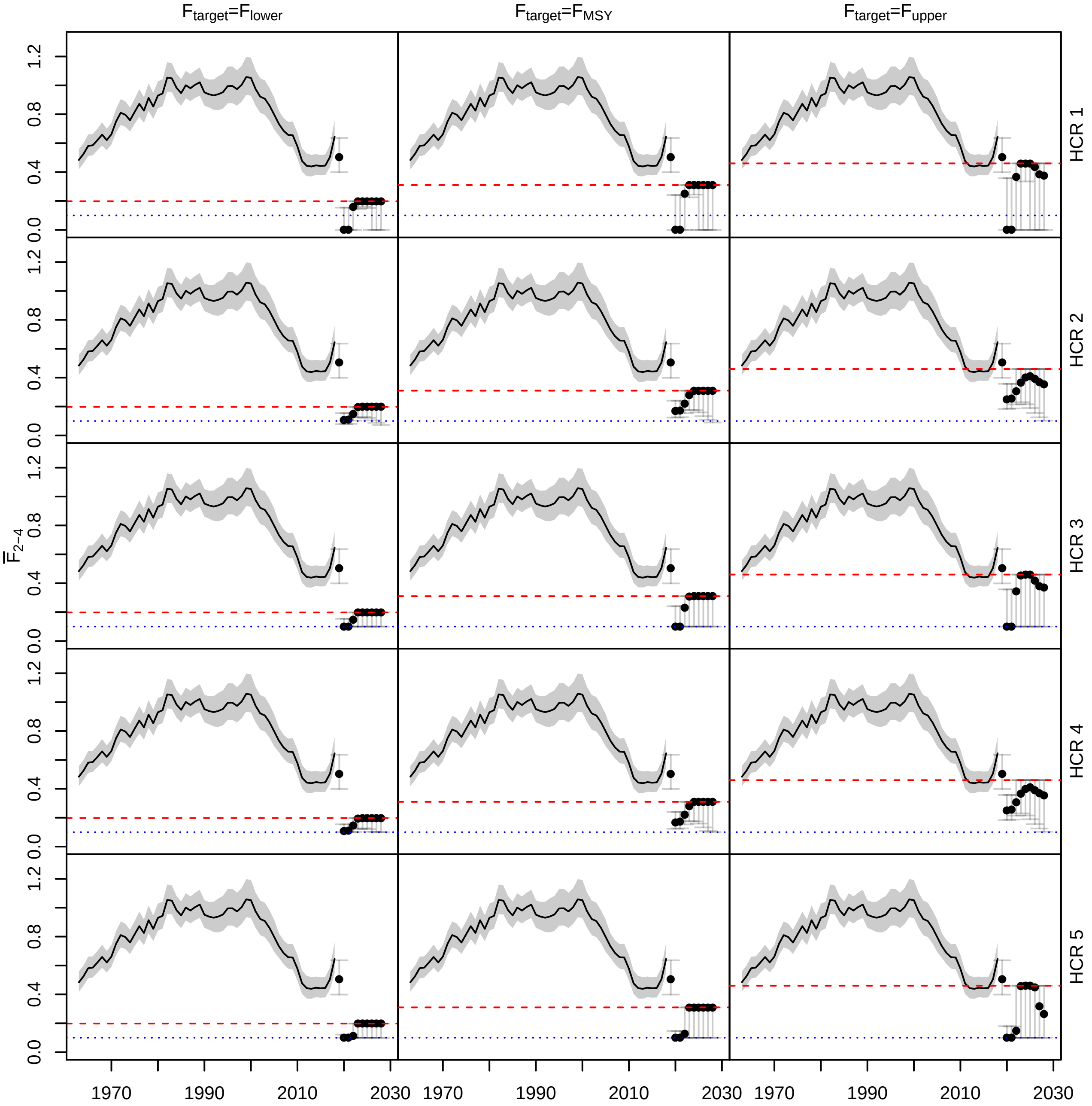


Figure A4.24: Average fishing mortality on North Sea cod for ages 2-4 ($\bar{F}_{2-4}$) estimated for all HCRs when recruitment is a random walk with negative drift. The points are the median $\bar{F}_{2-4}$ and the segments the 95% confidence intervals estimated from the 5000 replicates. The red dashed line is $F_{target}$ and the blue dotted line is $F = 0.1$.

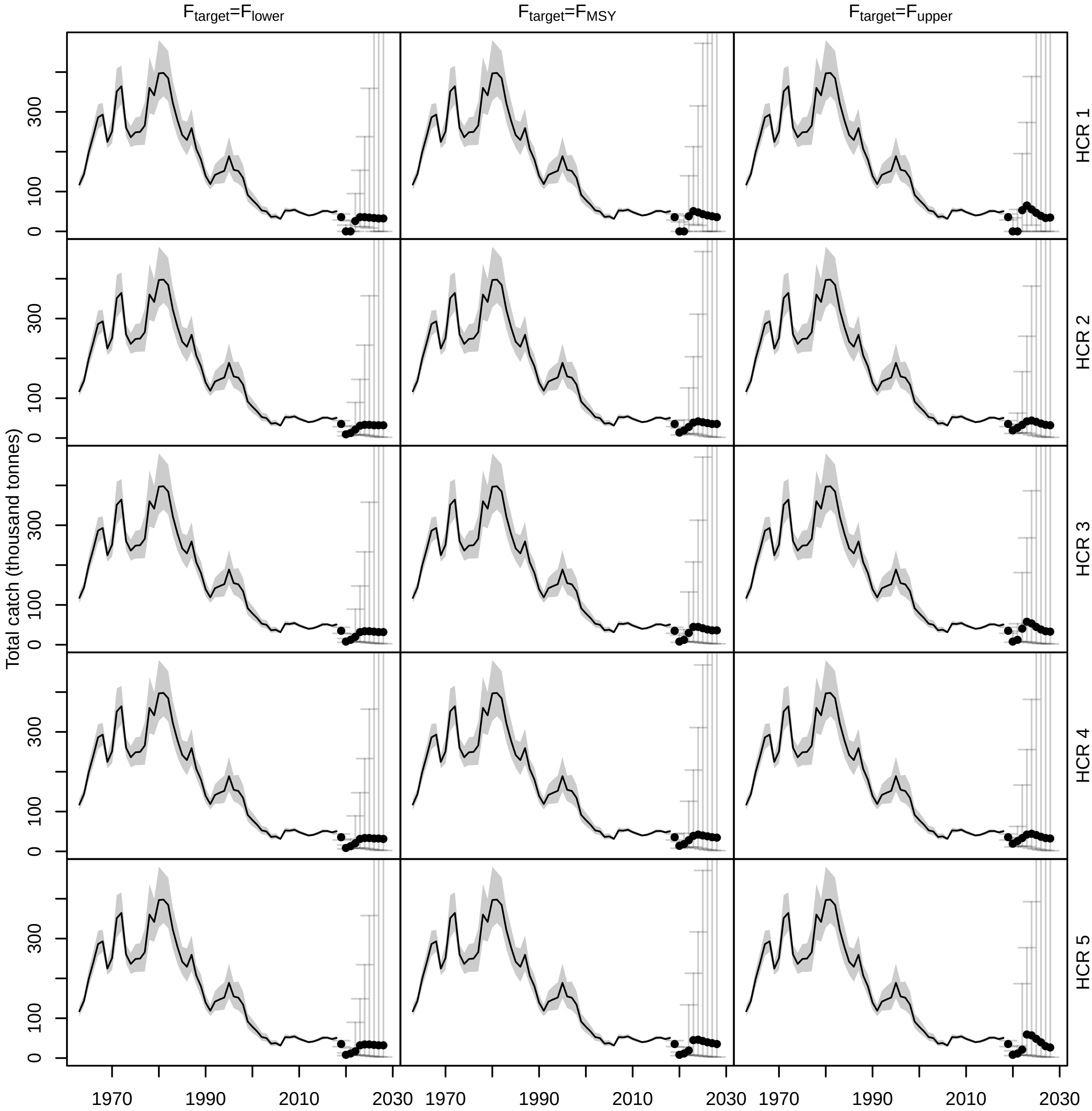


Figure A4.25: Total catch of North Sea cod estimated for all HCRs when recruitment is a random walk with negative drift. The points are the median catch and the segments the 95% confidence intervals estimated from the 5000 replicates. For plotting convenience the upper limits of the confidence intervals are not shown.

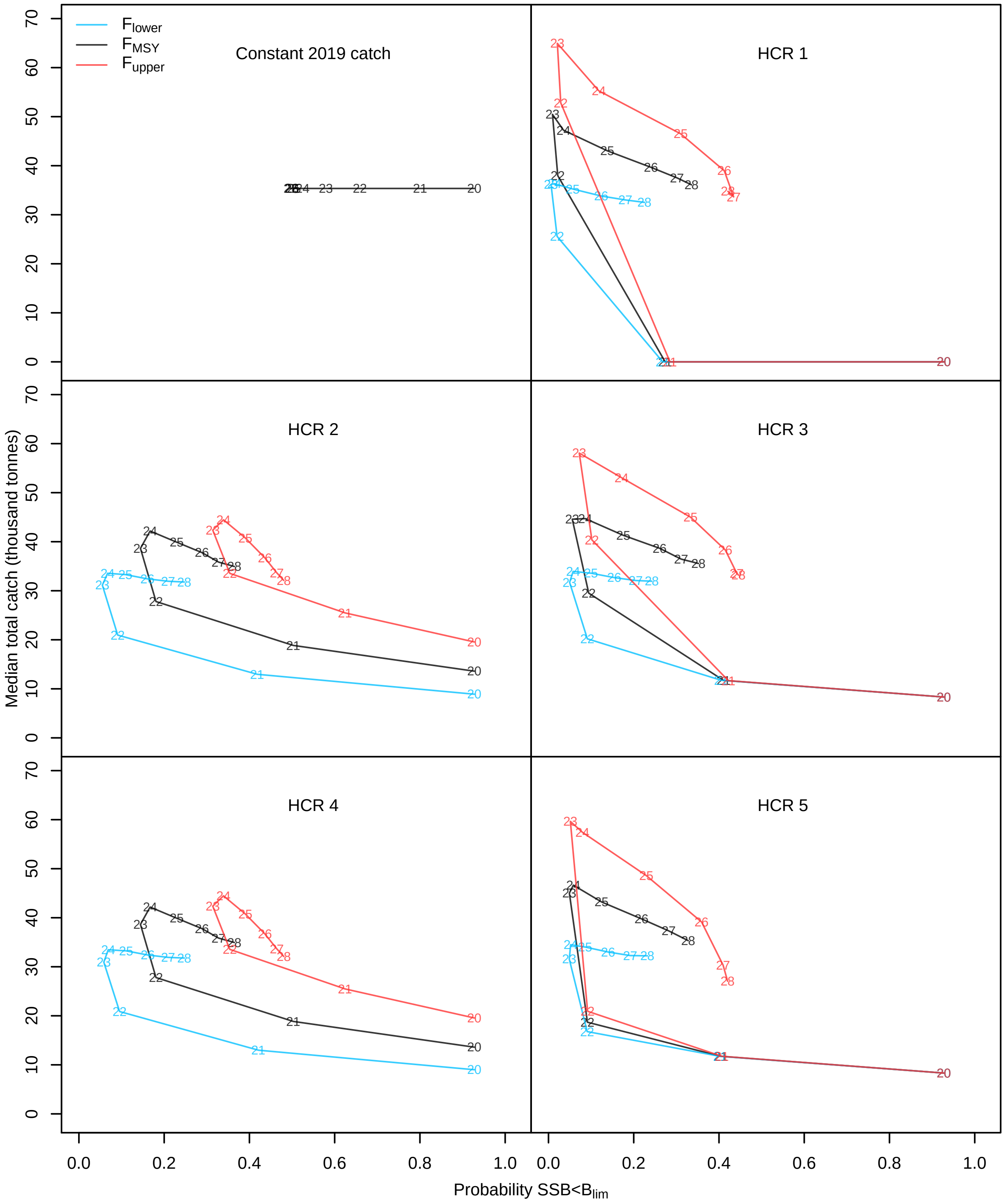


Figure A4.26: North Sea cod median total catch against probability of SSB falling below $B_{lim}$ when recruitment is a random walk with negative drift. The numbers correspond to the forecast years (2020-2028). Each color represents a $F_{target}$ option. Only one color is showed for the constant 2019 catch base case because the results did not depend on the $F_{target}$.

#### A4.3 Cumulative catch with confidence intervals for all forecast years

To supplement Figures 6 and 8, Figures A4.27 and A4.28 show the median and 95% confidence interval around the cumulative catch for all forecast years (2019-2018). For all species and $F_{target}$ levels, the constant 2019 catch ST was the one showing the smallest confidence interval and HCR 1 was the one showing the widest. This could be explained by the fact that when the catch is constant uncertainty in cumulative catch in the forecast is only given by the current perception of the stock and process errors. For the HCRs uncertainty also depends on the path for each replicates (how $F$ changes in the forecast as a function of SSB). HCR 1 had the widest confidence interval before large variations in $\bar{F}$ happened between replicates, see for example the confidence intervals in Figure A4.12.

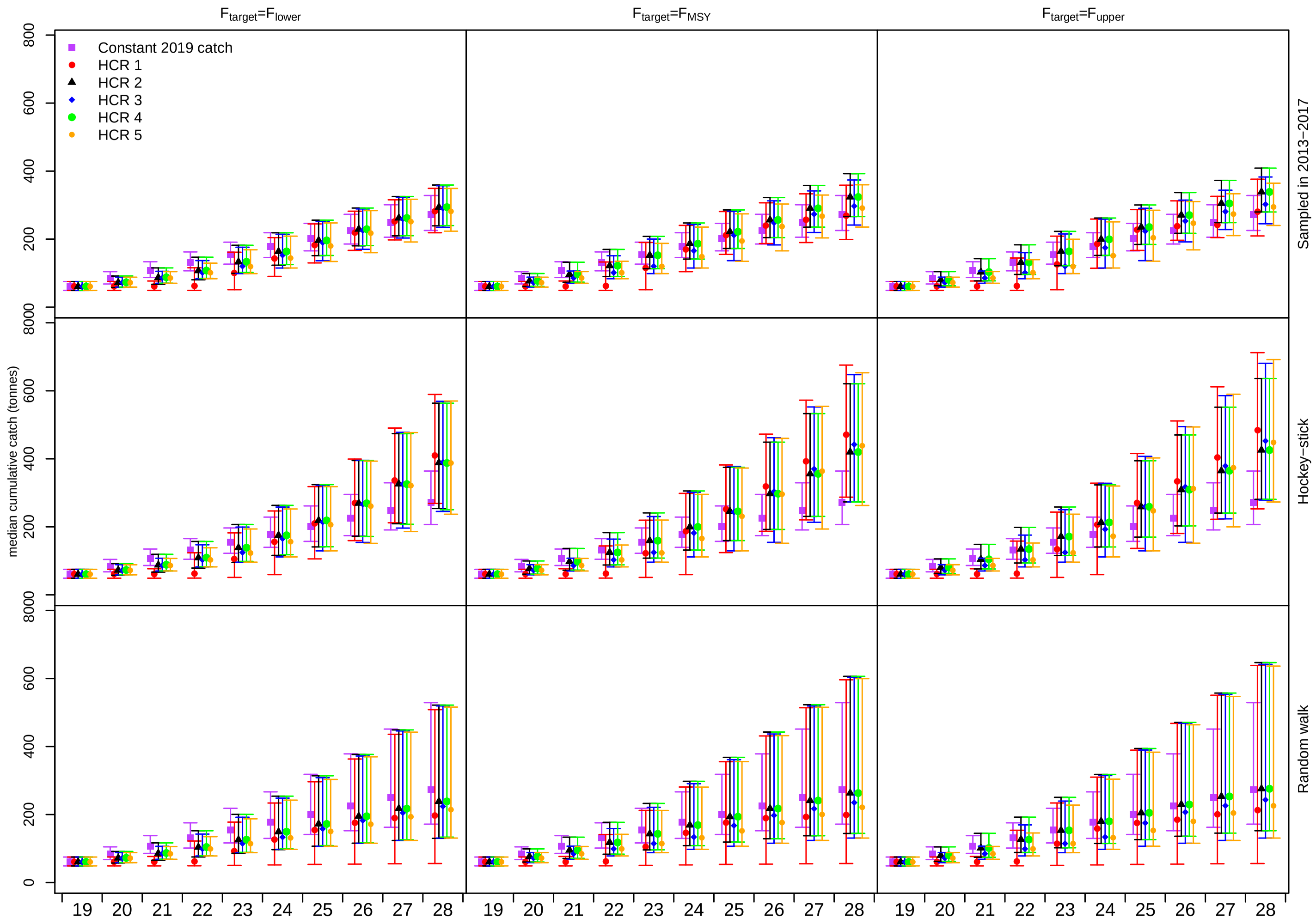


Figure A4.27: WBSS herring cumulative catch in the forecast period (2019-2028). The symbols correspond to the median cumulative catch and the segments are the 95% confidence intervals estimated from the 5000 replicates.

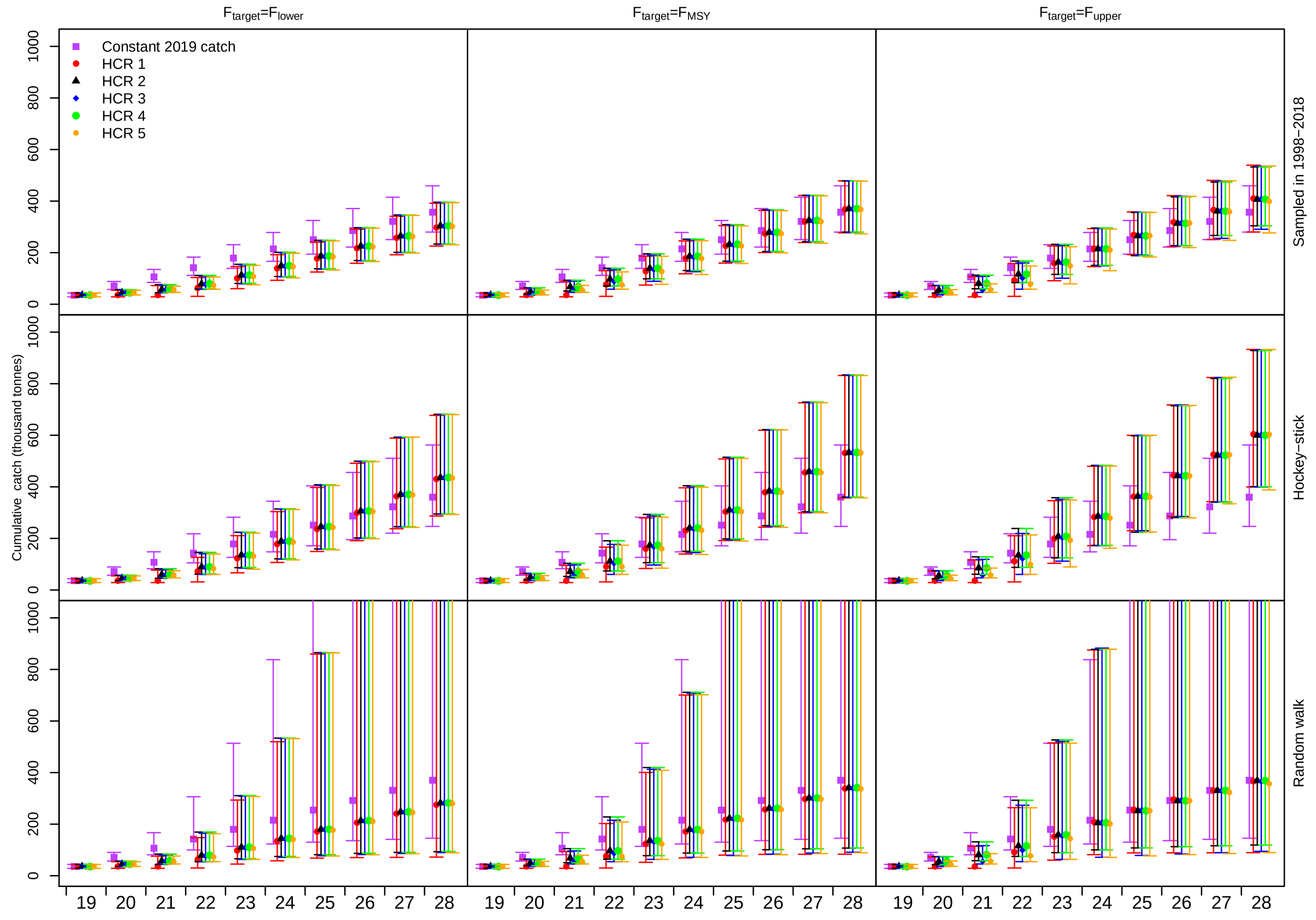


Figure A4.28: North Sea cod cumulative catch in the forecast period (2019-2028). The symbols correspond to the median cumulative catch and the segments are the 95% confidence intervals estimated from the 5000 replicates. For plotting convenience the upper limits of the confidence intervals are not shown in the random walk case. They reach a maximum of 4424 thousand tonnes for the constant catch scenario and $F_{target} = F_{lower}$ and $F_{MSY}$ and of 4696 thousand tonnes for the HCR 1 and $F_{target} = F_{upper}$.

### A5 Results of the long-term analysis and MSY reference point estimation

Given the current recruitment regime for WBSS herring, fishing at $F_{MSY}$ did not result in a median $SSB \geq B_{lim}$ in the long-term and the probability of falling below $B_{lim}$ was 98.1% (Figure A5.1). Even fishing at $F_{lower}$ only resulted in a median SSB slightly lower than $B_{lim}$ and a probability of rebuilding of 37.2% depending on the recruitment assumption. For North Sea cod, rebuilding above $B_{lim}$ is only compromised when $F_{target} = F_{upper}$, with a probability of rebuilding around 60% while it was more than 99% for the other two scenarios.

When recruitment is assumed to follow a stock-recruitment relationship, the probability of rebuilding for WBSS herring is improved (Figure A5.2). Fishing at $F_{MSY}$ allows a rebuilding probability of 88%. This is still below the 95% probability constraint used in ICES to estimate the reference points. This difference may be due to the fact that the stock-recruitment pairs used in this study include the most recent pairs and the stock perception has decreased since the reference points were estimated. For North Sea cod, assuming the stock-recruitment relationship mainly improves rebuilding probability for $F_{target} = F_{upper}$ that becomes now more than 95% and therefore consistent with ICES precautionary constraint.

This simple exercise highlights how the recruitment assumption highly influences the results of the projections.

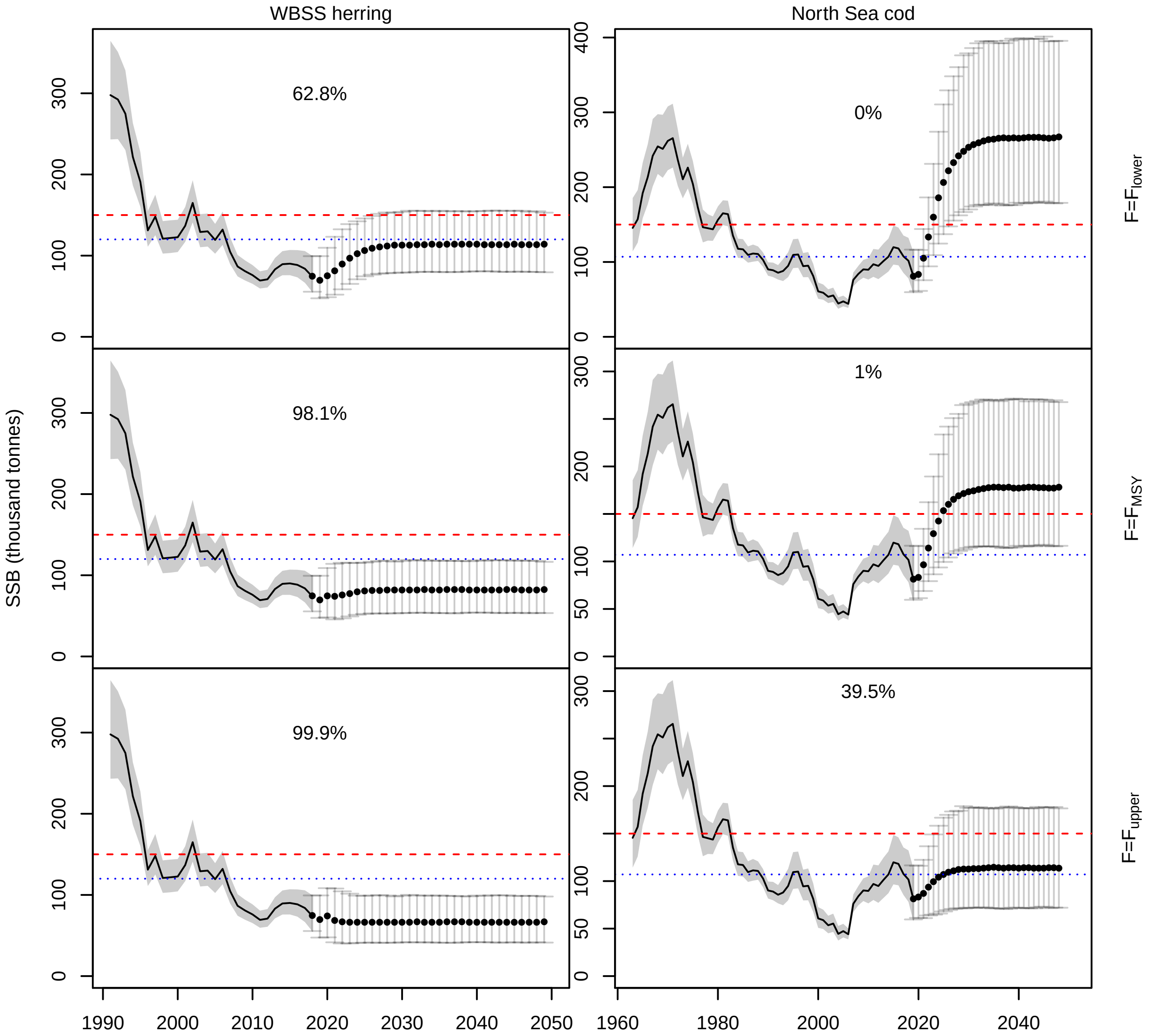


Figure A5.1: SSB estimated for long-term forecasts where $\bar{F}$ is fixed at the three different $F_{target}$ values assuming recruitment is sampled from the assessment estimates. The forecast was run for 30 years for both species. The points are the median SSB and the segments the 95% confidence intervals estimated from the 5000 replicates. The red dashed line is $MSYB_{trigger}$ and the blue dotted line is $B_{lim}$. Probability of SSB falling below $B_{lim}$ is given as percentage for each forecast.

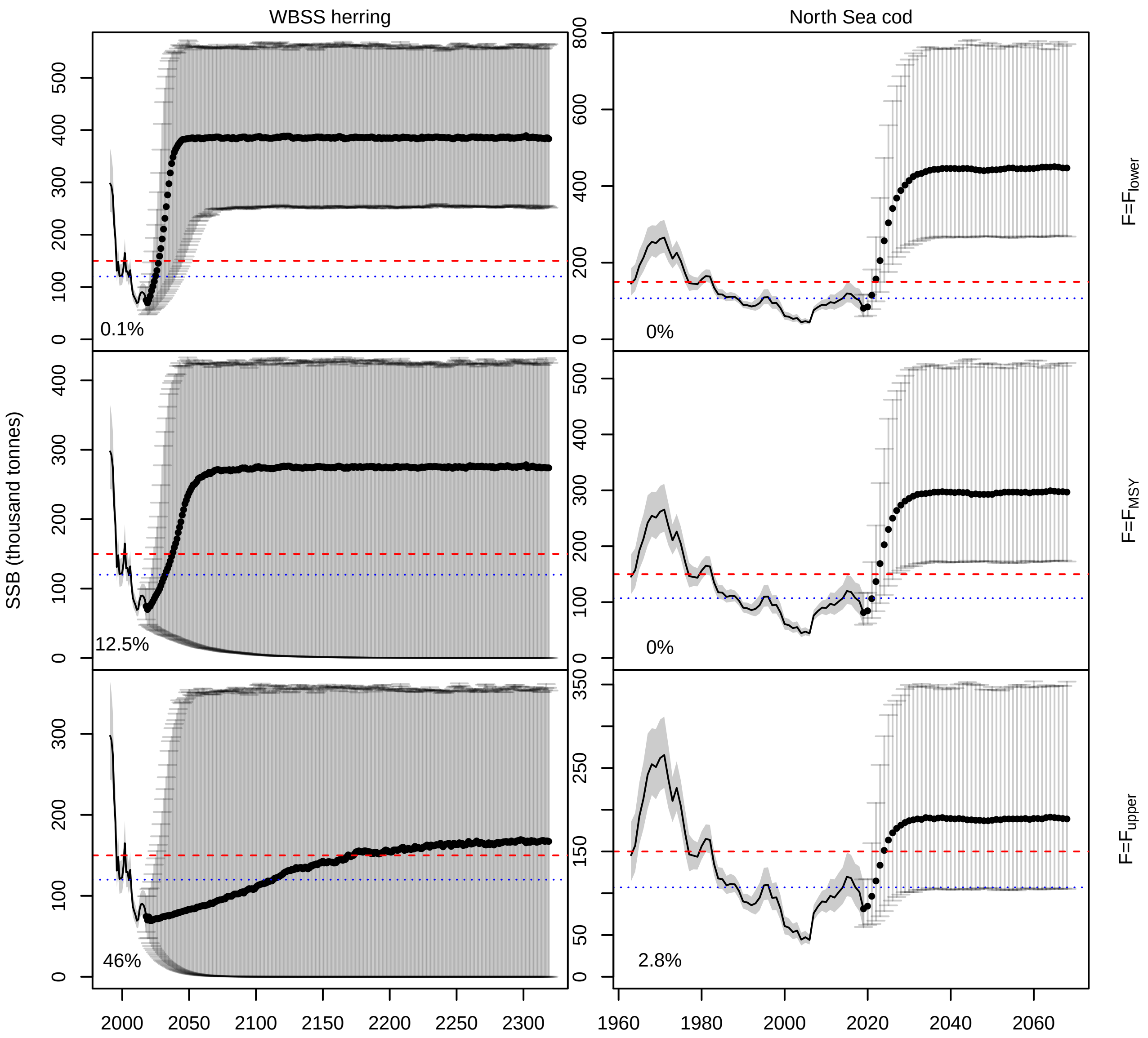


Figure A5.2: SSB estimated for long-term forecasts where $\bar{F}$ is fixed at the three different $F_{target}$ values assuming recruitment followsa hockey-stick relationship. The forecast was run for 300 years for herring (steady-state was slow to reach for $F_{target} = F_{upper}$) and 50 years for cod. The points are the median SSB and the segments the 95% confidence intervals estimated from the 5000 replicates. The red dashed line is $MSYB_{trigger}$ and the blue dotted line is $B_{lim}$. Probability of SSB falling below $B_{lim}$ is given as percentage for each forecast.

The yield curves obtained to estimate MSY reference points for the different recruitment assumptions are given in Figure A5.3. The results from the MSY reference point estimation for the assumption of recruitment randomly sampled from the assessment estimates are given in Table A5.1 and in Table A5.2 for the hockey-stick recruitment assumption. Assuming recruitment is sampled from the assessment estimates, resulted in a $F_{MSY}$ value for herring that is higher than the current $F_{MSY}$ (0.31) but with a corresponding $B_{MSY}$ largely below $B_{lim}$ (120000 t). Assuming recruitment follows a hockey-stick resulted in a smaller $F_{MSY}$ compared to the first option but in a large estimate for $B_{MSY}$, significantly larger than the current stock status. Results for cod were very consistent with the current value of 0.31 for $F_{MSY}$. Both assumptions resulted in similar $F_{MSY}$ but assuming a hockey-stick resulted in higher values for $B_{MSY}$ and $MSY$ compared to a flat recruitment.

Table A5.1: MSY reference point median estimates and 95% confidence interval (obtained from the forecast runs where $F = F_{MSY}$), for the assumption of recruitment randomly sampled from the assessment estimates.

| Stocks | $F_{MSY}$ ($y^{-1}$) | $B_{MSY}$ (t) | $MSY$ (t) |
|---|---|---|---|
| Herring | 0.496 (0.357-0.693) | 48400 (27692-76840) | 35834 (30232-44640) |
| Cod | 0.311 (0.246-0.392) | 177318 (115369-271548) | 37499 (25821-54331) |

Table A5.2: MSY reference point median estimates and 95% confidence interval (obtained from the forecast runs where $F = F_{MSY}$), for the assumption of recruitment following a hockey-stick stock-recruitment relationship.

| Stocks | $F_{MSY}$ ($y^{-1}$) | $B_{MSY}$ (t) | $MSY$ (t) |
|---|---|---|---|
| Herring | 0.248 (0.179-0.346) | 342763 (187285-505675) | 113146 (80656-151693) |
| Cod | 0.310 (0.245-0.391) | 297048 (172653-527979) | 61722 (37051-110763) |

The reason for $F_{MSY}$ for herring being lower than 0.31 when recruitment followed a hockey-stick is not very clear. This difference could be due to the fact that since ICES (2018), the perception of the stock has decreased so the SSB and recruitment pairs have changed and selectivity and other inputs to the forecast may also differ. The estimation of reference points in this study is mainly used to put the medium-term forecast results into context, and should not be considered as a cautious estimation of new reference points.

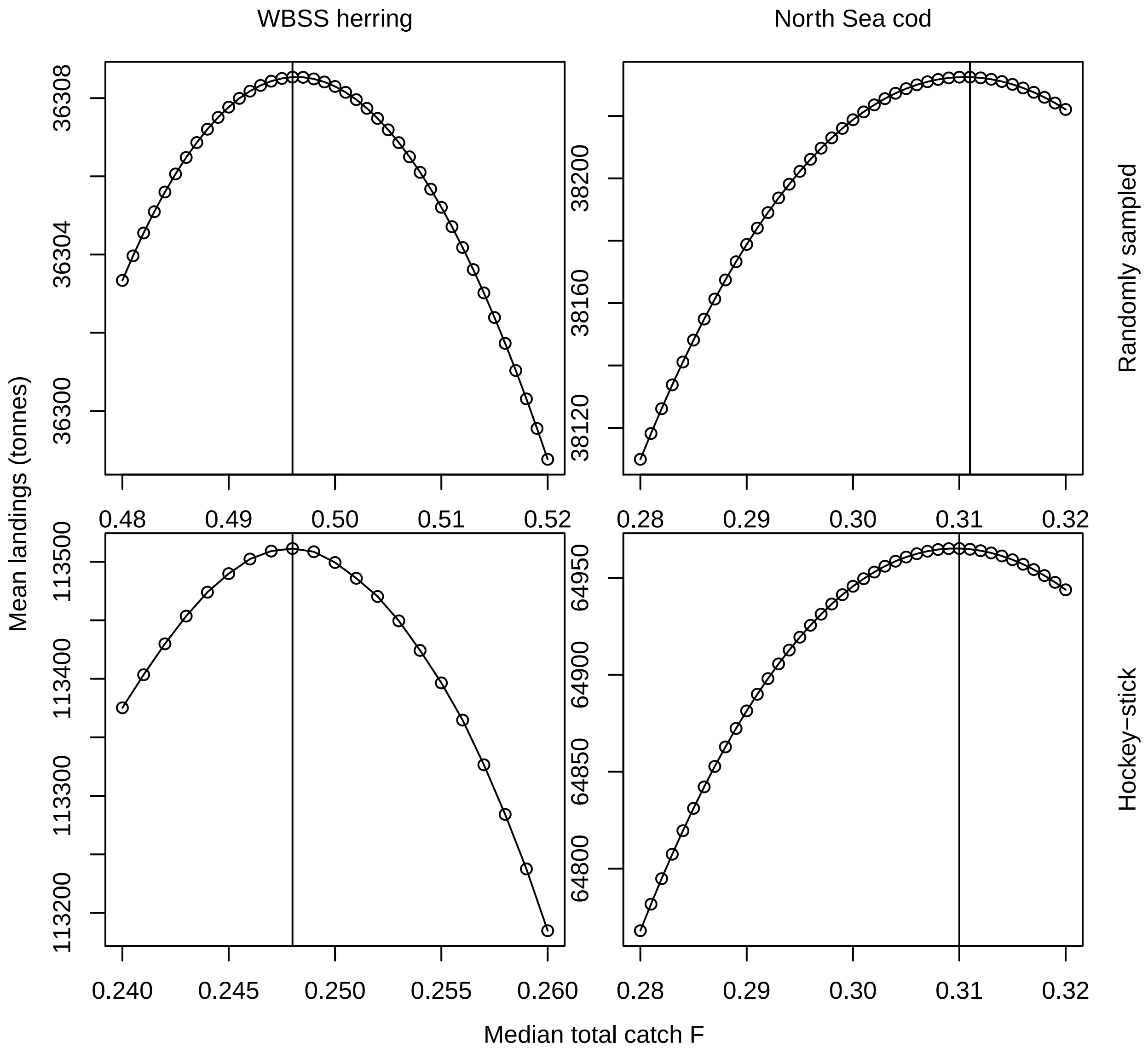


Figure A5.3: Mean landings over the 5000 replicates as a function of median total catch fishing mortality for the different recruitment assumptions and stocks. The vertical lines show the position of the $F$ that maximizes the mean landings.

### References

ICES 2018. Report of the benchmark workshop on pelagic stocks (WKPELA 2018). Tech. rep., 12–16 February 2018, ICES HQ, Copenhagen, Denmark. ICES CM 2018/ACOM:32.

ICES 2019a. Herring (Clupea harengus) in subdivisions 20–24, spring spawners (Skagerrak, Kattegat, and western Baltic). Tech. rep., ICES Advice on fishing opportunities, catch, and effort Baltic Sea and Greater North Sea Ecoregions, Report of the ICES Advisory Committee 2019. ICES Advice 2019.

ICES 2019b. Report of the Herring Assessment Working Group for the Area South of 62°N (HAWG). Tech. rep., 23-31 January 2019 and 13-21 March 2019. ICES HQ, Copenhagen, Denmark.

ICES 2019c. Working Group on the Assessment of Demersal Stocks in the North Sea and Skagerrak (WGNSSK). Tech. rep., ICES Scientific Reports, 1:7.
